# A PP2A-Integrator complex fine-tunes transcription by opposing CDK9

**DOI:** 10.1101/2020.07.12.199372

**Authors:** Stephin J. Vervoort, Sarah A. Welsh, Jennifer R. Devlin, Elisa Barbieri, Deborah A. Knight, Matteo Costacurta, Izabela Todorovski, Conor J. Kearney, Jarrod J. Sandow, Stefan Bjelosevic, Zheng Fan, Joep H. A. Vissers, Karolina Pavic, Ben P. Martin, Gareth Gregory, Isabella Y. Kong, Edwin D. Hawkins, Simon J. Hogg, Madison J. Kelly, Andrea Newbold, Kaylene J. Simpson, Otto Kauko, Kieran F. Harvey, Michael Ohlmeyer, Jukka Westermarck, Nathanael Gray, Alessandro Gardini, Ricky W. Johnstone

## Abstract

Gene expression is tightly controlled by Cyclin-dependent kinases (CDKs) which regulate the RNA Polymerase II (RNAPII) transcription cycle at discrete checkpoints. RNAPII pausing is a CDK9-controlled rate-limiting process that occurs shortly after initiation and is required for spatio-temporal control of transcription in multicellular organisms. We discovered that CDK9-mediated RNAPII pause-release is functionally opposed by a protein phosphatase 2A (PP2A) complex. PP2A dynamically competes for key CDK9 substrates, DSIF and RNAPII, and is recruited to transcription pausing sites by INTS6, a subunit of the Integrator complex. INTS6 depletion disrupts the Integrator-PP2A association and confers resistance to CDK9 inhibition. This results in unrestrained activity of CDK9 and dysregulation of acute transcriptional responses. Pharmacological PP2A activation amplifies RNAPII pausing mediated by CDK9 inhibitors and synergizes therapeutically in a model of MLL-rearranged leukemia. These data demonstrate that finely-tuned gene expression relies on the delicate balance of kinase and phosphatase activity throughout the transcription cycle.

**HIGHLIGHTS:**

- Loss of INTS6 confers resistance to CDK9 inhibition
- INTS6 recruits PP2A to Integrator and chromatin
- PP2A/INTS6 complexes functionally oppose CDK9
- PP2A/INTS6 fine-tune acute transcriptional responses
- Synergistic anti-cancer activity between PP2A activators and CDK9 inhibitors

## INTRODUCTION

Spatiotemporal control of gene expression is essential for appropriate cellular and organismal development, and is required to direct functional responses to a range of intrinsic and extrinsic cellular cues (Roeder, 2019). This essential process is tightly regulated and highly dynamic, directed by the activity of RNA polymerase II (RNAPII) (Chen et al., 2018; Roeder, 2019) and the concerted action of cyclin-dependent kinases (CDKs) and their cognate cyclins (Malumbres, 2014). Following transcription initiation, RNAPII is arrested shortly after transcribing the first 60-100 base pairs of most genes and acquires a ‘paused’ conformation, which is stabilized by the DRB sensitivity inducing factor (DSIF) and the negative elongation factor (NELF) (Chen et al., 2018; Core and Adelman, 2019; Vos et al., 2018b). Paused RNAPII is still engaged in transcription, yet unable to further elongate the nascent transcript, owing to reduced mobility and impaired binding of NTPs. Dynamic phosphorylation of RNAPII, NELF, and DSIF facilitates the transition of the paused polymerase into active elongation (Chen et al., 2018; Core and Adelman, 2019; Vos et al., 2018a).

The release of RNAPII from its paused state enables productive transcriptional elongation and requires the kinase activity of the positive transcription elongation factor (pTEFb), comprised of CDK9 and cyclin T (Chen et al., 2018; Core and Adelman, 2019). Through the activity of CDK9, pTEFb phosphorylates DSIF, NELF, and the RNAPII C-terminal domain (CTD) at serine 2 (Ser2) (Chen et al., 2018; Core and Adelman, 2019). While phosphorylation of NELF prompts its release from the transcription machinery, phospho-DSIF remains associated with the elongating polymerase but undergoes allosteric changes. RNAPII-CTD phosphorylation at Ser2 (phospho-Ser2) is spread across 52 repeats of the conserved amino acid heptad (YSPTSPS), and provides a recruitment platform for multi-protein complexes that regulate co-transcriptional processes including capping, splicing, termination, and histone methylation (Harlen and Churchman, 2017). The activity of pTEFb at the transcriptional pause site is mediated by its recruitment through a variety of protein interactions, including transcription factors such as c-MYC (Eberhardy and Farnham, 2001), the bromodomain-containing protein BRD4 that binds to acetylated histone lysine molecules (Hargreaves et al., 2009; Jang et al., 2005), and the Super Elongation Complex (SEC) that is recruited to proximal promoters by proteins such as the MLL-AF9 oncogenic fusion protein (Lin et al., 2010; Luo et al., 2012).

Small molecule inhibitors of CDK9 kinase activity, such as flavopiridol, have been effectively used to study the molecular and biochemical events that underpin transcriptional pause-release and to ascertain greater mechanistic detail of transcription dynamics (Chao and Price, 2001; Martin et al., 2020). Indeed, pharmacological inhibition of CDK9 blocks productive transcriptional elongation concomitant with accumulation of hypo-phosphorylated RNAPII at the transcriptional pause site. Given the functional interactions of pTEFb with oncogenic transcription factors such as c-MYC and MLL-fusion proteins, and that transcriptional amplification is an underlying molecular characteristic of cancer (Bradner et al., 2017), small molecule inhibitors of CDK9 have previously been applied in the oncology setting, although a selective inhibitor of CDK9 has yet to be approved as standard treatment for any cancer (Fujinaga, 2020).

Despite extensive literature on the various RNAPII-CTD kinases including the “transcriptional kinases” (CDK7, 8, 9, 11, 12, 13), relatively little is known about the phosphatases that target RNAPII-CTD and oppose pTEFb kinase activity (Jeronimo et al., 2016). A number of phosphatases including RTR1, SSu72, Cdc14b, Glc7, Fcp1 and Scp1 have been shown to de-phosphorylate RNAPII-CTD in mammals and/or lower organisms (Mayfield et al., 2016). Recently, CDK9-mediated phosphorylation of the yeast Spt5, a component of DSIF, was shown to be opposed by the PP1 phosphatase isoform Dis2 during the regulated progression from transcriptional elongation to termination (Parua et al., 2018). These data provide some evidence for functional interplay between kinases and phosphatases to dynamically regulate the function of key transcriptional machinery in lower eukaryotes, however detailed mechanistic information on the regulation of transcriptional pause-release through opposing kinase/phosphatase interactions in higher organisms remains scarce.

In this study, we used whole genome CRISPR-based screens to identify genes that regulate transcriptional pausing and MLL-rearranged leukemia cell death induced by perturbation of CDK9 (CDK9i). Knockout of INTS6, a component of the Integrator protein complex, suppressed CDK9i and allowed transcriptional elongation in human leukemia and solid tumor cells, and in untransformed *Drosophila melanogaster* cells. Integrator is a metazoan-specific, 14-subunit complex that associates with the RNAPII-CTD and cleaves nascent RNA species such as U small nuclear RNAs (snRNAs) and enhancer RNAs (eRNAs) (Lai et al., 2015; Rienzo and Casamassimi, 2016). Recently, Integrator was reported to regulate RNAPII pausing and elongation through its RNA-directed catalytic subunits INTS11/INTS9 (Elrod et al., 2019; Gardini et al., 2014). Here, we uncovered that INTS6 forms a distinct functional module of Integrator, bridging its interaction with the PP2A phosphatase. While PP2A is a well-known phosphatase complex that can act as a tumor suppressor in a wide range of human malignancies (Fowle et al., 2019), it has not been studied in association to transcriptional dynamics. We found that PP2A is recruited at actively transcribed genes through INTS6 to oppose CDK9 activity and promote pausing by controlling the phosphorylation turnover of key CDK9 substrates, including the RNAPII-CTD and DSIF. These discoveries provided a strong molecular rationale to combine CDK9i with small molecule activators of PP2A (SMAPs), resulting in enhanced transcriptional pausing and synergistic therapeutic efficacy in MLL-rearranged AML. Taken together, we reveal a previously undescribed function for Integrator in regulating transcriptional pause release through the recruitment of PP2A to chromatin via INTS6, and provide new mechanistic insights into the fine tuning of transcription through the opposing enzymatic activities of CDK9 and PP2A.

## RESULTS

### Loss of the Integrator component INTS6 confers resistance to CDK9 inhibition

Pharmacological inhibition of CDK9 (CDK9i) has been proposed to be an effective therapeutic strategy in hematopoietic malignancies, particularly MLL-rearranged leukemias. On a molecular level, CDK9 inhibition results in widespread RNAPII pausing, preventing most coding and noncoding genes from being effectively transcribed. To identify proteins that functionally antagonize CDK9 activity on both the phenotypic and transcriptional level we performed a series of genome-wide CRISPR-screens on the MLL-rearranged leukemia cell lines THP-1 and MV4;11 treated with CDK9i using the following readouts: (1) long-term cell survival and (2) nascent RNA transcription (Figure 1A). Positive selection, genome-wide CRISPR screens were initially performed in THP-1 cells transduced with a 120,000 single guide RNA (sgRNA) library (GeckoV2) under selective pressure from three chemically distinct CDK9i (AZ-5576, A159 and Dinaciclib). This revealed selective enrichment of single guide RNAs (sgRNAs) targeting the Integrator complex subunit *INTS6* in response to all three CDK9i (Figure 1B left panel). These findings were extended in analogous long-term survival screens using THP-1 and MV4;11 cells transduced with a distinct 76,000 sgRNA library (Brunello) confirming that deletion of *INTS6* conferred resistance to CDK9i (Figure 1B right panel, Figure S1A).

**FIGURE 1:**
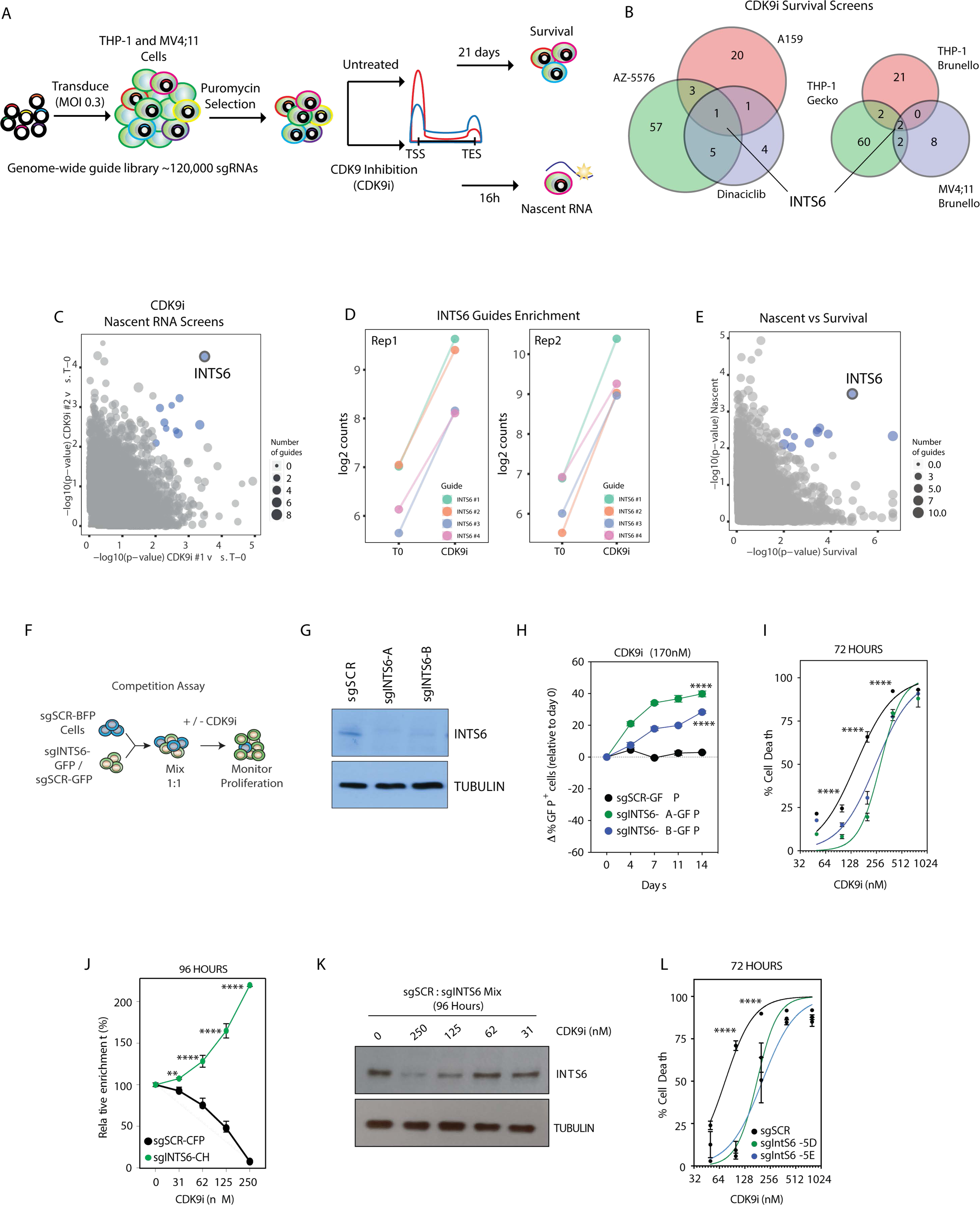
Loss of INTS6 confers resistance to CDK9 inhibition. **A.** CRISPR-Cas9 genome-wide CDK9i survival and nascent RNA screens. **B.** Venn diagrams for significant enriched sgRNAs (CDK9i versus untreated at Tend) in survival screens in THP-1-Cas9 cells (left; different CDK9 inhibitors) and using different genome-wide sgRNA libraries (adjusted p-value < 0.1 for > 3 sgRNAs in 1 or more replicate screens) **C.** Enriched sgRNAs for replicate nascent RNA screens in THP-1-Cas9 cells (significance relative to T0). **D.** Enrichment of *INTS6* targeting sgRNAs in CDK9i-treated THP-1-Cas9 cells at endpoint relative to T0. **E.** Comparison of enriched sgRNAs for genome-wide CDK9i nascent RNA and survival screens. **F.** Overview of sgRNA Competitive proliferation assays. **G.** Western blot of THP-1-Cas9 cells expressing indicated sgRNAs. **H.** Competitive proliferation assay for THP-1-Cas9 cells expressing indicated sgRNAs in the presence of CDK9i. **I.** Annexin-V analysis of THP-1-Cas9 cells expressing indicated SCR and *INTS6* targeting sgRNAs treated as indicated for 72 hours. **J.** Competitive proliferation assay for CFP-Cas9 expressing HeLa sgSCR and mCherry(CH)-Cas9 expressing HeLa sg*INTS6* cells treated with CDK9i as indicated for 96 hours. **K.** Western blot of HeLa cells expressing sgSCR or sgINTS6 (mixed) treated with CDK9i as indicated for 96 hours. **L.** Annexin-V analysis of *D.melanogaster* S2-Cas9 expressing indicated SCR and *IntS6* targeting sgRNAs and treated as indicated for 72 hours. Blue dots (Figures C and E) represent nominal p-value < 0.01. Figures G-L are representative of 3 independent experiments. Figures H, I, J, and L were analyzed by 2-way ANOVA, ** p < 0.01, **** p<0.0001.

To directly assess functional antagonism to CDK9 inhibition on a transcriptional level, a flow cytometry-based readout of nascent RNA production using click chemistry-based fluorescent 5-ethynyl uridine (EU) labelling was utilized (Figure S1B). As expected, CDK9i treatment of THP-1 cells resulted in a time- and dose-dependent reduction of EU incorporation (Figure S1C), validating the utility of this assay to determine the acute effect of CDK9i on nascent RNA production. Genome-wide CRISPR-screens in THP-1 cells were performed and assessed for nascent RNA production to select a cell population that retained the capacity to efficiently incorporate EU in the presence of AZ-5576 (Figure S1D). In concordance with the CDK9i survival screens shown in figures 1A and 1B, *INTS6* sgRNAs were enriched in two independent nascent RNA production screens (Figure 1C-D, Figure S1E), and *INTS6* loss was the only resistance mechanism identified consistently across all screens (Figure 1E). Enrichment of sgRNAs targeting Integrator subunits *INTS8* and *INTS12* was observed in both phenotypic and transcription screens, suggesting that these may also play a role in CDK9 antagonism (Figure S1F).

In competitive proliferation and cell death validation assays (Figure 1F), *INTS6* deletion conferred a strong selective survival advantage to THP-1 and MV4;11 cells treated with AZ5576 (Figure 1G-I, Figure S1G) as well as in response to a CDK9 proteolysis targeting chimera (PROTAC, 22-533) (Figure S1H-I). In agreement with results from genome-wide CDK9i survival screens, analogous assays with sgRNAs targeting *INTS3* and *INTS11* did not confer resistance to CDK9 inhibition and demonstrated the critical dependency of these proteins in vehicle treated THP-1 cells (Figure S1J and K). *INTS6* deletion did not affect proliferation of untreated cells (Figure S1L), nor did it provide resistance to the chemotherapeutic agent, cytarabine, or the inhibitor of CDK7 and CDK12, THZ1 (Figure S1M). Importantly, the resistance to CDK9i conferred by *INTS6* loss in THP-1 cells was completely rescued by ectopic expression of full length INTS6 (Figure S1N-P). As validation for the nascent RNA screens, selective *INTS6* deletion allowed for efficient EU incorporation in the presence of AZ5576, whereas RNA production was almost completely abrogated in AZ5576-treated SCR sgRNA control cells (Figure S1Q).

The importance of INTS6 in regulating biological responses to CDK9i beyond MLL-rearranged leukemia cells was demonstrated in solid human tumor cells. Indeed, CRISPR-mediated deletion of *INTS6* in HeLa cells provided a competitive proliferation and survival advantage over control cells following prolonged exposure to AZ5576 (Figure 1J-K and Figure S1R). To determine if the functional antagonism between CDK9 and INTS6 is evolutionarily conserved, *IntS6* was deleted in *D. melanogaster* derived S2 cells. As observed in solid and liquid human tumor cell lines, deletion of *IntS6* in S2 cells conferred resistance to CDK9 inhibition (Figure 1L), a phenotype that was rescued by ectopic expression of full-length IntS6 protein (Figure S1S-T). Taken together, our results reveal that depletion of INTS6 confers resistance to CDK9i across multiple model systems and organisms on both phenotypic and transcriptional levels.

### INTS6 bridges the interaction between Integrator and PP2A

To determine how INTS6 regulates the transcriptional and biological responses to CDK9 inhibition, a series of LC-MS/MS analyses were performed to investigate the extended INTS6/Integrator interactome. Immunoprecipitation (IP) of INTS6 from THP-1 and MV4;11 whole cell lysates resulted in the co-purification of the other 13 members of the Integrator complex (Figure S2A-B). Interestingly, the PPP2R1A/PPP2R1B structural subunits and the PPP2CA catalytic subunit (PP2A-C) of the PP2A phosphatase complex were highly enriched, as was the PPP1CC subunit of PP1 (Figure 2A-B, S2A-B). Members of the RNAPII complex, including the Rpb1/POLR2A catalytic core, also co-purified with INTS6 (Figure 2A-B, S2A-B). To examine the PP2A-Integrator association in a different human cell line, the endogenous INTS6 and PP2A proteomes were isolated from HeLa nuclear extracts. Affinity purification with a monoclonal antibody specific for PP2A-C identified Integrator and RNAPII as the most abundant interactors of nuclear PP2A, as measured by iBAQ values (Figure 2C).

**FIGURE 2:**
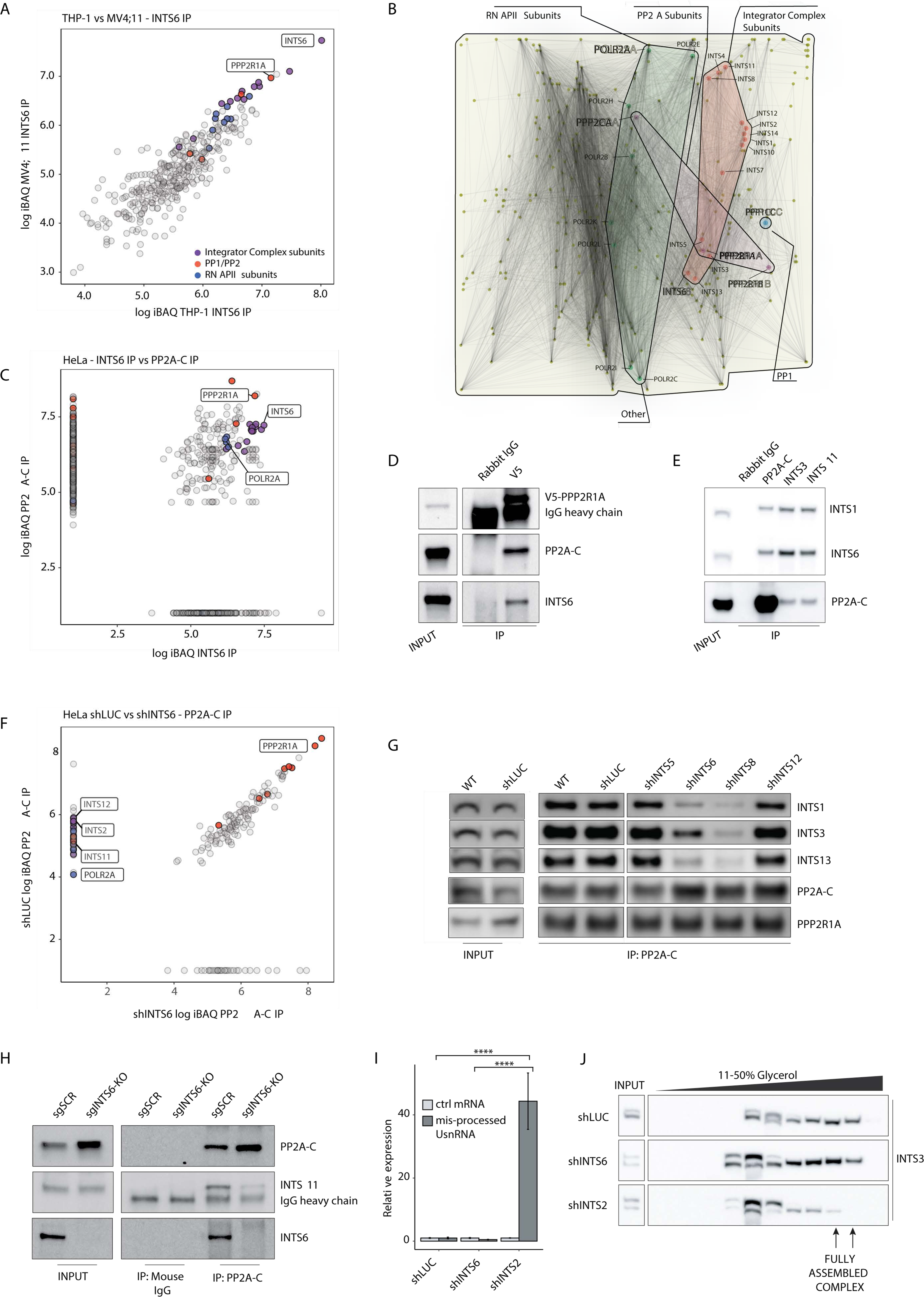
INTS6 bridges the interaction between Integrator and PP2A. **A.** Log10 iBAQ scores of proteins identified in INTS6 IP mass spectrometry experiments for THP-1 versus MV4;11 cells (filtered for proteins identified in isotype-control IP experiments). **B.** Protein-protein interaction network of INTS6, RNAPII, and PP2A interaction partners identified in THP-1 and MV4;11 INTS6 IP experiments. **C.** Log10 iBAQ scores of proteins identified in INTS6 versus PP2A-C IPs in HeLa cells. Co-IP western blot of **D.** endogenous V5-tagged PPP2R1A IP in THP-1 cells and **E.** endogenous PP2A-C, INTS3, and INTS11 IP in HeLa cells. **F.** Log10 iBAQ scores of proteins identified in PP2A-C IP mass spectrometry experiments in HeLa shLUC versus shINTS6 cells. **G.** Co-IP western blot for PP2A-C in shLUC, shINTS5, shINTS6, shINTS8, and shINTS12-infected HeLa cells. **H.** Co-IP western blot for PP2A-C in THP-1-Cas9 sgSCR and sgINTS6-KO cells. **I.** Relative expression levels of misprocessed UsnRNA from shLUC, shINTS2, or shINTS6-infected HeLa cells. Values normalized by expression of ribosomal 18S and *GUSB* is used as control (ctrl) mRNA. **J.** Western blot of glycerol gradient fractions of nuclear extracts from shLUC, shINTS2, or shINTS6-infected HeLa cells. IP mass spectrometry experiments are representative of 3 (THP-1, HeLa) or 2 (MV4;11) independent experiments. Western blots are representative of 3 independent experiments. qPCR (I) was analyzed by one-way ANOVA followed by Tukey’s HSD post-hoc test, **** p<0.0001.

As in MLL-rearranged leukemia cells, endogenous INTS6 co-precipitated all Integrator complex subunits along with RNAPII, and PP2R1A and PP2A-C subunits (Figure 2C). The interaction between endogenous PP2A and INTS6 was validated in co-IP assays in both THP-1 and HeLa cells (Figure 2D-E).

PP2A is a highly abundant protein complex found in both the nucleus and cytoplasm with the ability to dephosphorylate a wide range of protein targets (Eichhorn et al., 2009). Importantly, PP2A has never been directly implicated in the regulation of chromatin-associated transcriptional complexes. While most PP2A is cytoplasmic, sub-cellular fractionation of HeLa cells indicated that a substantial fraction of the phosphatase complex was found in both the nuclear compartment and the insoluble chromatin fraction, where the Integrator complex was also detected (Figure S2C). To assess whether INTS6 was directly implicated in recruiting PP2A to Integrator, affinity purification of chromatin-bound endogenous PP2A-C followed by LC-MS/MS was performed in HeLa cells expressing shRNA targeting INTS6, or luciferase (LUC) as a control (Figure 2F). As expected, IP of PP2A-C resulted in co-purification of Integrator subunits including INTS6, as well as RNAPII subunits including Rpb1/POLR2A (Figure 2F). However, following INTS6 depletion, PP2A-C no longer co-precipitated with Integrator or RNAPII components. To validate these results and assess specificity of the INTS6-PP2A association, INTS6 and other Integrator subunits including INST5, INTS8, and INTS12 were depleted by shRNA and IP/western blot assays were performed (Figure 2G). Depletion of INTS6 greatly affected the Integrator/PP2A association in HeLa nuclear extract, while the interaction was unperturbed by depletion of either INTS5 or INTS12 (Figure 2G). Interestingly, depletion of INTS8 also decreased the Integrator/PP2A association, consistent with the results of our genetic screens where loss of INTS8 conferred some degree of resistance to CDK9i (Figure S1F), suggesting that INTS8 may also contribute to the Integrator/PP2A functional interaction. The importance of INTS6 in mediating the Integrator-PP2A interaction was validated in THP-1 cells, with sgRNA-mediated deletion of INTS6 resulting in decreased co-IP of PP2A-C and INTS11 (Figure 2H, Figure S2D). Moreover, analysis of the Ints6 proteome in *D. melanogaster* S2 cells confirmed the evolutionary conservation of this interaction between Integrator, PP2A, and RNAPII. (Figure S2E). Finally, the effect of INTS6 depletion on the basal catalytic activity and overall assembly of the Integrator complex was assessed. A PCR-based assay to measure levels of uncleaved (misprocessed) UsnRNAs was utilized to assess catalytic Integrator activity (Figure 2I). While depletion of a large structural subunit, such as INTS2, increased misprocessed UsnRNA levels by more than 40-fold, depletion of INTS6 had little or no effect. Moreover, depletion of INTS6 did not perturb migration of the Integrator complex on a glycerol gradient, unlike INTS2, suggesting that INTS6 is dispensable for the structural integrity of the complex (Figure 2J). Collectively, these data suggest that INTS6 is part of a novel Integrator module, distinct from the catalytic module, that is responsible for recruitment of PP2A.

### INTS6 mediates the recruitment of PP2A to the transcription pause-release checkpoint

Evidence of a physical interaction between INTS6, RNAPII, and PP2A, as well as the demonstration that loss of INTS6 opposed transcriptional pausing induced by CDK9i, raised the possibility that PP2A might co-localize with Integrator and RNAPII throughout the genome. ChIP-seq assays in THP-1 cells were performed using antibodies specific for CDK9, BRD4, RNAPII, INTS6, INTS11, and PPP2R1A (Figure 3A-B, S3A). As with BRD4, RNAPII, and CDK9, Integrator subunits INTS6 and INTS11 localized to chromatin at or around transcription start sites (TSS). Treatment of cells with AZ5576 resulted in retention of RNAPII at the TSS proximal region, consistent with transcriptional pausing induced by CDK9i (Figure 3A-B). BRD4 also accumulated at the TSS following AZ5576 treatment, consistent with the physical interaction between BRD4 and RNAPII (Jang et al., 2005). Interestingly, there was a concomitant increase in CDK9 and modest change in INTS6 and INTS11 binding at the TSS (Figure 3A). Consistent with our biochemical data that revealed PP2A in the chromatin fraction of HeLa cells (Figure S2C), PPP2R1A occupied nearly all actively transcribed genes, mirroring the localization of BRD4, RNAPII, CDK9, and the INTS6 and INTS11 subunits of the Integrator complex, and was also enriched around the TSS (Figure 3A-B, S3A).

**FIGURE 3:**
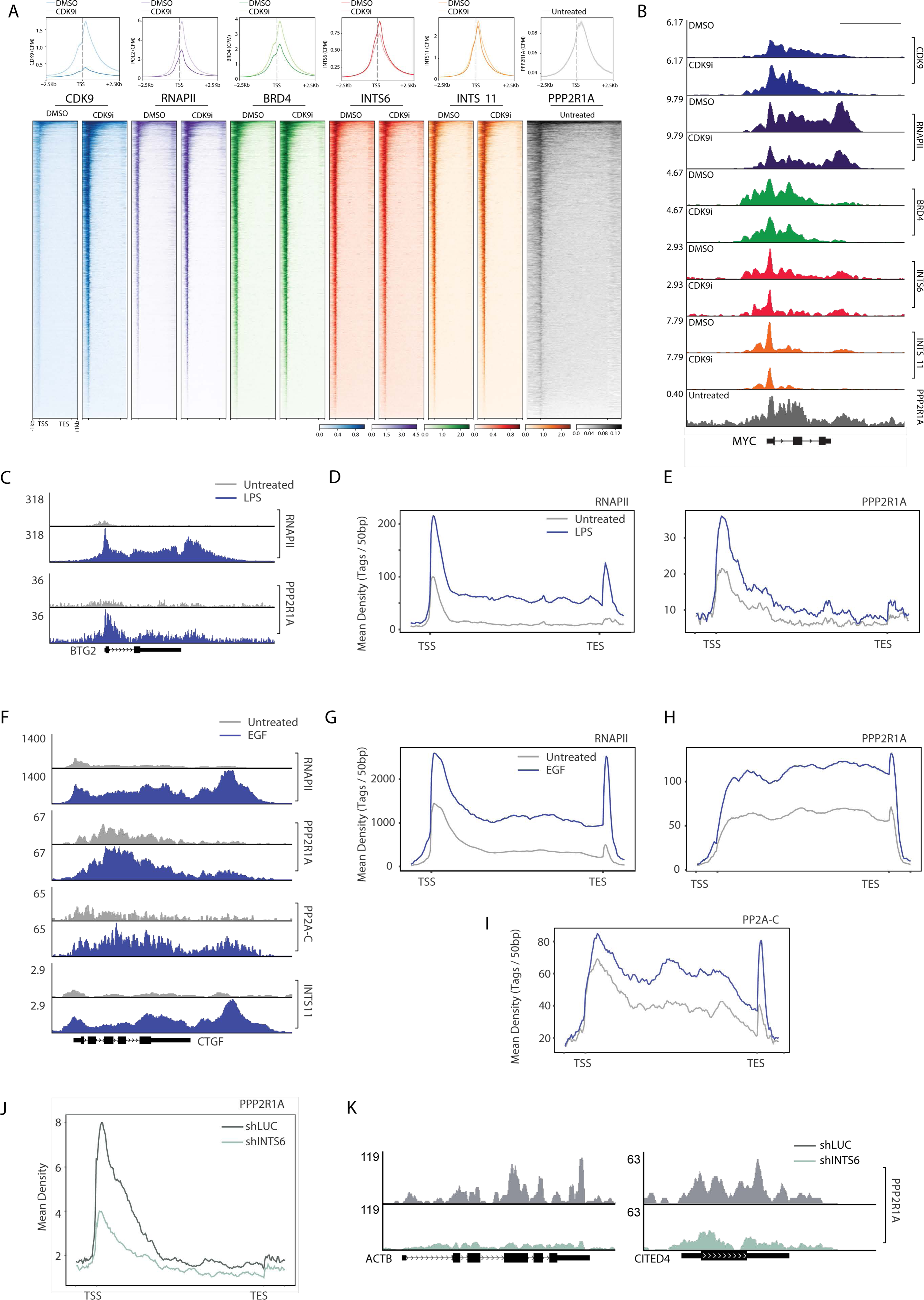
INTS6-dependent dynamic recruitment of PP2A at actively transcribed genes. **A.** Average profiles of ChIP-seq signal for CDK9, RNAPII, BRD4, INTS6, INTS11, and PPP2R1A around the TSS, plus metagene occupancy heatmaps for indicated proteins in THP-1 cells treated as indicated for 2 hours. **B.** Representative IGV ChIP-profiles for indicated proteins in THP-1 cells treated as indicated for 2 hours. **C.** Representative IGV profile for RNAPII and PPP2R1A ChIP signal in THP-1 cells untreated or acutely treated with 5.0µg/mL LPS for 3 hours. Average gene profiles for **D.** RNAPII and **E.** PPP2R1A ChIP-seq at LPS-induced genes (n=35) inTHP-1 cells under the same conditions. **F.** Representative IGV ChIP-profile for indicated proteins in HeLa cells untreated or acutely treated with EGF (0.1µg/mL) for 15 minutes. INTS11 ChIP-seq tracks are from a published dataset (Gardini et al., 2014). Average gene profiles for **G.** RNAPII, **H.** PPP2R1A and **I.** PP2A-C ChIP-seq at EGF-induced genes (n=50) in HeLa cells under the same conditions. **J.** Average gene profile at PP2A-occupied genes (n=194) and **K.** representative IGV ChIP-profiles for PPP2R1A ChIP-seq in shLUC and shINTS6 infected HeLa. Scale bar for C represents 5kb.

To assess the dynamic recruitment of PP2A in response to a coordinated and rapid transcriptional stimulus, ChIP-seq assays were performed using THP-1 cells treated for 3h with lipopolysaccharide (LPS). Treatment with LPS resulted in the expected increase of RNAPII recruitment along the entire gene body of LPS-responsive genes (Figure 3C-D). Strikingly, LPS treatment stimulated rapid recruitment of PPP2R1A at the TSS and across the gene body of LPS-responsive genes (Figure 3C-D). To further assess PP2A recruitment to chromatin in response to a physiological stimulus, Epidermal Growth Factor (EGF) was used to stimulate transcription of Immediate Early Genes in HeLa cells (Gardini et al., 2014). A dramatic recruitment of RNAPII, PPP2R1A, PP2A-C, and INTS11 along the entire gene body and the 3’ end of EGF-responsive genes was observed following 15 min treatment with EGF (Figure 3F-I). Notably, closer analysis of read alignment around the TSS revealed that PP2A recruitment gradually increased after initiation and peaked immediately 3’ of the pausing site (Figure S3B). To determine if INTS6 is important for the recruitment of PP2A to actively transcribed genes, ChIP-seq assays were performed on HeLa cells expressing shRNAs targeting LUC or INTS6 cultured with EGF for 15 min. Depletion of INTS6 dramatically reduced EGF-stimulated binding of PPP2R1A across EGF-responsive genes (Figure 3J-K).

Taken together, these findings demonstrate that pTEFb, Integrator, and PP2A localize to chromatin under basal conditions and that PP2A is actively recruited near pausing sites and across the gene body of genes selectively activated by different physiological stimuli. Moreover, these data demonstrate that INTS6 plays an important role in recruiting PP2A to chromatin.

### INTS6/PP2A dynamically controls CDK9 substrate phosphorylation levels

To investigate whether the functional antagonism between CDK9 and PP2A/INTS6 was reflected at the phosphorylation level, reverse phase protein arrays (RPPA) were performed to accurately quantify RNAPII CTD phosphorylation, in parallel with total phospho-proteomic analyses (Figure 4A). As expected, RPPA analysis of RNAPII CTD phosphorylation demonstrated that CDK9i treatment resulted in a time-dependent loss of phospho-Ser2 (pS2), phospho-Ser5 (pS5), and phospho-Ser7 (pS7) CTD levels in THP-1 cells transduced with control sgRNA (sgSCR) (Figure 4B). In contrast, THP-1 cells with stable deletion of INTS6 (sgINTS6-KO) were largely refractory to CDK9 inhibition, with no significant loss of phosphorylation observed for any of the phospho-CTD antibodies (Figure 4B). Characterization of the total phospho-proteome by mass spectrometry identified a large number of CDK9i responsive phospho-peptides in sgSCR THP-1 cells that were resistant to CDK9i treatment in sgINTS6-KO cells (Figure 4C and Figure S4A-B). Importantly, this included key CDK9 substrates such as *SUPT5H* (DSIF, SPT5; Figure 4D and S4C), *LEO1*, and degenerate heptad repeats in the RNAPII CTD (*POLR2A;* Figure 4E and S4D). Gene-ontology (GO) analysis of phosphorylated proteins sensitive to CDK9i in sgSCR cells and resistant to CDK9i in sgINTS6-KO cells revealed a strong enrichment for mRNA processing and RNA metabolic processes, suggesting that PP2A/INTS6 oppose CDK9 kinase activity on multiple levels (Figure 4F and Figure S4E).

**FIGURE 4:**
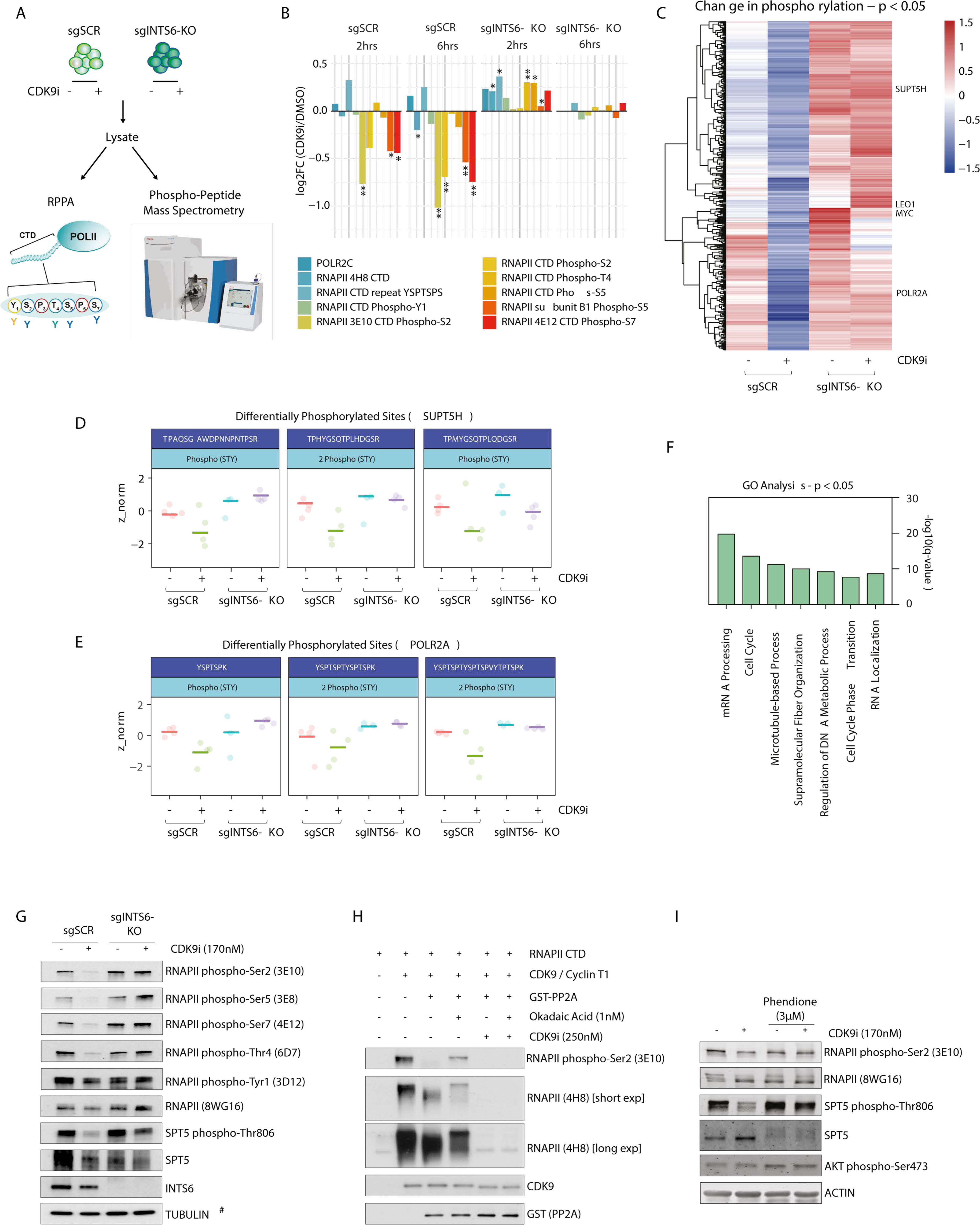
Loss of INTS6/PP2A results in decreased turnover of CDK9 substrates. **A.** Overview of CDK9-dependent phosphorylation studies in THP-1-Cas9 sgSCR and sgINTS6-KO cells. **B.** RNAPII CTD reverse phase protein array (RPPA): log fold change in relative fluorescence intensity for indicated antibodies in CDK9i-treated versus untreated THP-1-Cas9 sgSCR and sgINTS6-KO cells at indicated time points. **C.** Heatmap of Z-scores of phospho-peptides in THP-1-Cas9 sgSCR and sgINTS6-KO cells treated as indicated for 2 hours, p-value < 0.05. Identification of differentially phosphorylated peptides **D.** *SUPT5H* and **E.** *POL2RA* peptides in THP-1-Cas9 sgSCR and sgINTS6-KO cells treated as indicated for 2 hours. **F.** GO-analysis of proteins with differentially phosphorylated peptides in CDK9i-treated THP-1-Cas9 sgSCR (compared to DMSO-treated THP-1-Cas9 sgSCR and CDK9i-treated THP-1-Cas9 sgINTS6-KO cells; p < 0.05). **G.** Western blot of THP-1-Cas9 sgSCR and sgINTS6-KO cells treated as indicated for 2 hours. **H.** Western blot of *in vitro* recombinant kinase / phosphatase assay; recombinant CTD was incubated with ATP in the presence/absence of CDK9/CyclinT1 and/or PP2A as indicated for 30 minutes. **I.** Western blot of THP-1 cells treated as indicated (15 minutes pre-treatment with the PP2A inhibitor Phendione and 2 hours with CDK9i). Three biological replicates were analysed for RPPA and ten biological replicates were analysed for phospho-peptide mass spectrometry. Western blots are representative of three independent experiments. For Figure G, TUBULIN (#) is representative of individual immunoblots for phospho-CTD sites. RPPA (B) was analysed using Welch unpaired t-test, * p<0.05, ** p<0.01.

Western blot validation using antibodies targeting specific phospho-residues of the RNAPII-CTD and phospho-DSIF (SPT5) confirmed that THP-1 sgINTS6-KO cells were refractory to the CDK9i mediated loss of phosphorylation observed in sgSCR control cells (Figure 4G). To assess whether the ability of PP2A/INTS6 to oppose CDK9 kinase activity is evolutionarily conserved, analogous experiments in *D. melanogaster* S2 cells demonstrated that *IntS6* deletion conferred resistance to CDK9i-mediated loss of RNAPII-CTD phospho-Ser2 levels (Figure S4F). To determine whether PP2A can directly dephosphorylate RNAPII CTD residues that are phosphorylated by pTEFb, *in vitro* kinase/phosphatase assays were performed using purified recombinant proteins (Figure S4G). As expected, incubation of an RNAPII CTD peptide with an active pTEFb complex comprising CDK9 and Cyclin T1 resulted in CTD hyper-phosphorylation, as evidenced by western blots showing increased migration and increased signal with antibodies that recognize pan-phospho RNAPII-CTD and specific phospho-Ser2 antibodies (Figure 4H). Addition of a GST-tagged PP2A catalytic subunit (PP2A-C), greatly reduced CDK9-mediated RNAPII-CTD phosphorylation, which was partially rescued through inhibition of PP2A catalytic activity using Okadaic Acid, an effect which was lost upon simultaneous CDK9 inhibition with AZ5576 (Figure 4H). Finally, to probe the importance of PP2A mediated CDK9 antagonism in an endogenous context, the recently reported specific PP2A inhibitor Phendione (Yue et al., 2020) was used along with distinct PP1/PP2A inhibitors Calyculin A and Okadaic Acid. Consistent with the phenotype observed in sgINTS6-KO THP-1 cells, PP2A inhibition with Phendione rescued the loss of RNAPII-CTD phospho-Ser2 and SPT5 phosphorylation observed upon CDK9 inhibition (Figure 4I), which was also observed upon co-treatment with dual PP1/PP2A inhibitors (Figure S4H). These data highlight an evolutionarily conserved kinase/phosphatase antagonism between CDK9 and PP2A/INTS6 complexes that regulates the critical transcription pause-release factors RNAPII and SPT5.

### Loss of INTS6/PP2A overrides a CDK9i-dependent block of transcription elongation

To assess the impact of INTS6/PP2A loss on CDK9i-induced suppression of nascent transcription, 4-thiouridine metabolic labelling followed by RNA sequencing (4sU-seq) was performed in sgINTS6-KO and sgSCR THP-1 cells (Figure 5A-D and Figure S5A). Data were normalized through the addition of 4sU-labelled RNA from *D. melanogaster* S2 cells. Strikingly, 4sU-seq analysis revealed that the global AZ5576-dependent decrease of transcription seen in sgSCR THP-1 cells, was significantly rescued to near-baseline levels in THP-1 sgINTS6-KO cells, even after six hours of sustained inhibition of CDK9 kinase activity (Fig. 5B). Although sensitivity to AZ5576 varied across different gene subsets, including highly expressed genes, a significant rescue of nascent transcription was observed globally in sgINTS6-KO cells (Figure 5C-D), including at the *IL6R* and *MYC* loci (Figure 5E, Figure S5B-C). ChIP-seq assays further demonstrated reduced CDK9i-mediated accumulation of total RNAPII at TSS proximal regions and maintenance of RNAPII-CTD phospho-Ser2 throughout the gene body and at the transcription end site (TES) in AZ5576-treated sgINTS6-KO THP-1 cells compared to sgSCR cells (Figure 5F, Figure S5D-H). Calculation of the pausing index (the ratio of total RNAPII ChIP-seq reads at TSS vs. gene body regions) for all transcribed genes showed that INTS6 loss reduced AZ5576-induced pausing in comparison to the effect seen in AZ5576-treated sgSCR THP-1 cells, suggesting maintenance of RNAPII pause-release and elongation in the absence of full CDK9 kinase activity (Fig. 5G). The evolutionary conservation of this phenotype was validated in *D. melanogaster* S2 cells, with reduced RNAPII pausing observed in AZ5576-treated sgIntS6 S2 cells compared to the effect of AZ5576 observed in sgSCR cells (Figure S5I).

**FIGURE 5:**
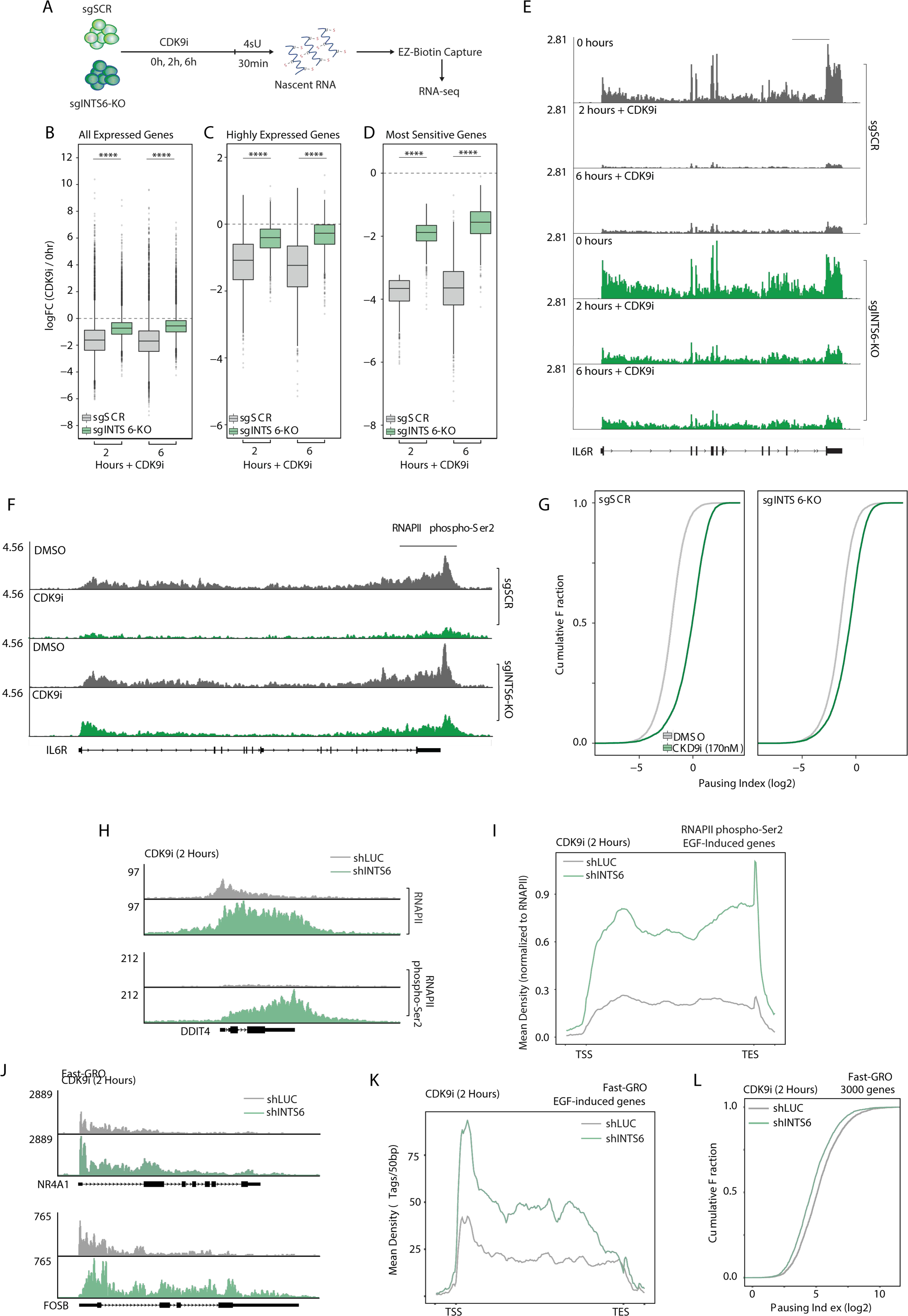
PP2A/INTS6 loss overrides CDK9i induced transcriptional pausing. **A.** Overview of 4sU labelling and analysis of nascent transcription in THP-1-Cas9 sgSCR and sgINTS6-KO cells. Log fold change in 4sU-seq signal (CPM) relative to untreated cells across in sgSCR and sgINTS6-KO THP-1-Cas9 cells for **B.** all expressed genes, **C.** highly expressed genes and **D.** CDK9i-sensitive genes. **E.** Example of 4sU-seq signal at the *IL6R* locus under indicated conditions. **F.** RNAPII phospho-Ser2 ChIP-seq signal at the *IL6R* locus in THP-1-Cas9 sgSCR and sgINTS6-KO cells treated as indicated for 2 hours. **G.** Pausing index in THP-1-Cas9 sgSCR and sgINTS6-KO cells treated for 2h with CDK9i. **H.** RNAPII and RNAPII phospho-Ser2 ChIP-seq signal at the *DDIT4* locus in shLUC and shINTS6 infected HeLa cells treated with CDK9i for 2 hours and acutely treated with EGF (0.1µg/mL) for 15 minutes. **I.** Average profile of RNAPII phospho-Ser2 ChIP-seq signal under the same conditions at EGF-response genes (n=50). **J.** Fast-GRO signal at the *FOSB* locus in shLUC or shINTS6 infected HeLa cells treated with CDK9i for 2 hours and acutely treated with EGF (0.1µg/mL) for 15 minutes. **K.** Average Fast-GRO signal across EGF response genes (n=50) under indicated conditions. **L.** Pausing index calculated by Fast-GRO for CDK9i-treated shLUC and shINTS6 infected HeLa cells at highest-expressed genes (n=2989). Pausing index defined as (TSS-region coverage)/(Gene-body coverage), where TSS-region is defined as 50bp upstream of TSS through 150bp downstream of TSS). Scale bar for E and F represents 10kb. Figures B-D were analysed by unpaired, two-sided students t-test, **** p<0.0001

To determine the effect of INTS6/PP2A loss on acutely activated genes that were suppressed by CDK9i, INTS6 was depleted in HeLa cells (Figure S5J) and cells were pre-treated with AZ5576 for two hours prior to stimulation with EGF for 15 minutes. ChIP-seq assays demonstrated that total RNAPII and RNAPII phospho-Ser2 accumulated genome-wide in INTS6-depleted HeLa cells treated with AZ5576, with accumulation most clearly observed across the gene bodies of EGF-responsive genes, in contrast to the reduced ChIP-seq signal observed in control shLUC cells (Figure 5H-I, S5K-M). The experiment was repeated with the CDK9i flavopiridol. After only one hour of treatment, INTS6-depleted cells showed broad escape of pausing as opposed to the control sample (Figure S5O-Q). Consistent with our biochemical evidence that also implicated the INTS8 subunit in the PP2A-Integrator association (Figure 2), depletion of INTS8 phenocopied INTS6 loss and also conferred resistance to CDK9i-induced pausing (Figure S5O-Q).

Consistent with the ChIP-seq data for RNAPII and RNAPII phospho-Ser2 detailed above, fastGRO assays (Barbieri et al., 2020) to assess nascent RNA production demonstrated that depletion of INTS6 in HeLa cells counteracted CDK9i-induced transcriptional pausing at all EGF-stimulated genes (Figure 5J-K). A broader analysis of the highest-expressed genes (n=2989) demonstrated that INTS6 depletion decreased the pausing index of nearly all active genes (Figure 5L), similar to the effects observed in THP-1 sgINTS6-KO cells treated with AZ5576 (Fig 5F). To determine if the counter-effects on CDK9i-induced transcriptional pausing observed following INTS6 depletion were phenocopied by depletion of PP2A, RNAPII phospho-Ser2 ChIP-seq assays were performed using HeLa cells with shRNA-mediated knockdown of PPP2R1A or control knockdown cells (shLUC) that had been treated with vehicle or AZ5576 for two hours prior to stimulation with EGF for 15 min (Figure S5Q). Depletion of PPP2R1A resulted in sustained EGF-induced RNAPII phospho-Ser2 coverage across EGF-stimulated genes in the presence of AZ5576 (Figure S5R). Collectively, data from various experimental systems revealed that loss of INTS6 promotes both basal and EGF-stimulated transcriptional elongation and decreases RNAPII susceptibility to CDK9i-induced pausing. Moreover, loss of PP2A phenocopied the transcriptional modulatory effects of INTS6 deletion/depletion.

### The INTS6/PP2A axis fine-tunes acute transcriptional responses

The fastGRO nascent RNA analysis demonstrated decreased responsiveness of HeLa cells to CDK9i-induced pausing (Figure 5J-L). Interestingly, even in the absence of CDK9i, our data suggested increased elongation and RNAPII processivity following INTS6 depletion (Figure S6A-B). To determine the role of INTS6/PP2A in transcriptional elongation and pause-release under physiological conditions of RNAPII pausing, ChIP-seq for total and phospho-Ser2 RNAPII was performed. Depletion of INTS6 in HeLa cells did not appreciably increase accumulation of RNAPII or phospho-Ser2 RNAPII near the TSS of EGF-responsive genes (Figure 6A), suggesting that the RNAPII initiation rate is unaffected by INTS6 depletion. In contrast, an increase in detectable RNAPII phospho-Ser2 across the gene bodies and 3’ ends of EGF-responsive genes was observed in INTS6-depleted HeLa compared to control cells (Figure 6A-B). The increase of RNAPII phospho-Ser2 was specifically observed following INTS6 depletion and was not observed upon depletion of the Integrator subunit INTS12 (Fig. 6B). Similar RNAPII and RNAPII phospho-Ser2 ChIP-seq results were obtained using HeLa cells depleted of PPP2R1A and treated with EGF, supporting the notion that INTS6 and PP2A functionally cooperate to regulate activated transcriptional elongation (Figure 6C-D). Further supporting the observed increase of RNAPII pause-release and elongation following perturbation of the INTS6/PP2A axis, measurement of nascent RNA production by transient-transcriptome sequencing (TT-seq) demonstrated a significant increase in the rate of EGF-induced transcription following depletion of INTS6 (Figure S6C).

**FIGURE 6:**
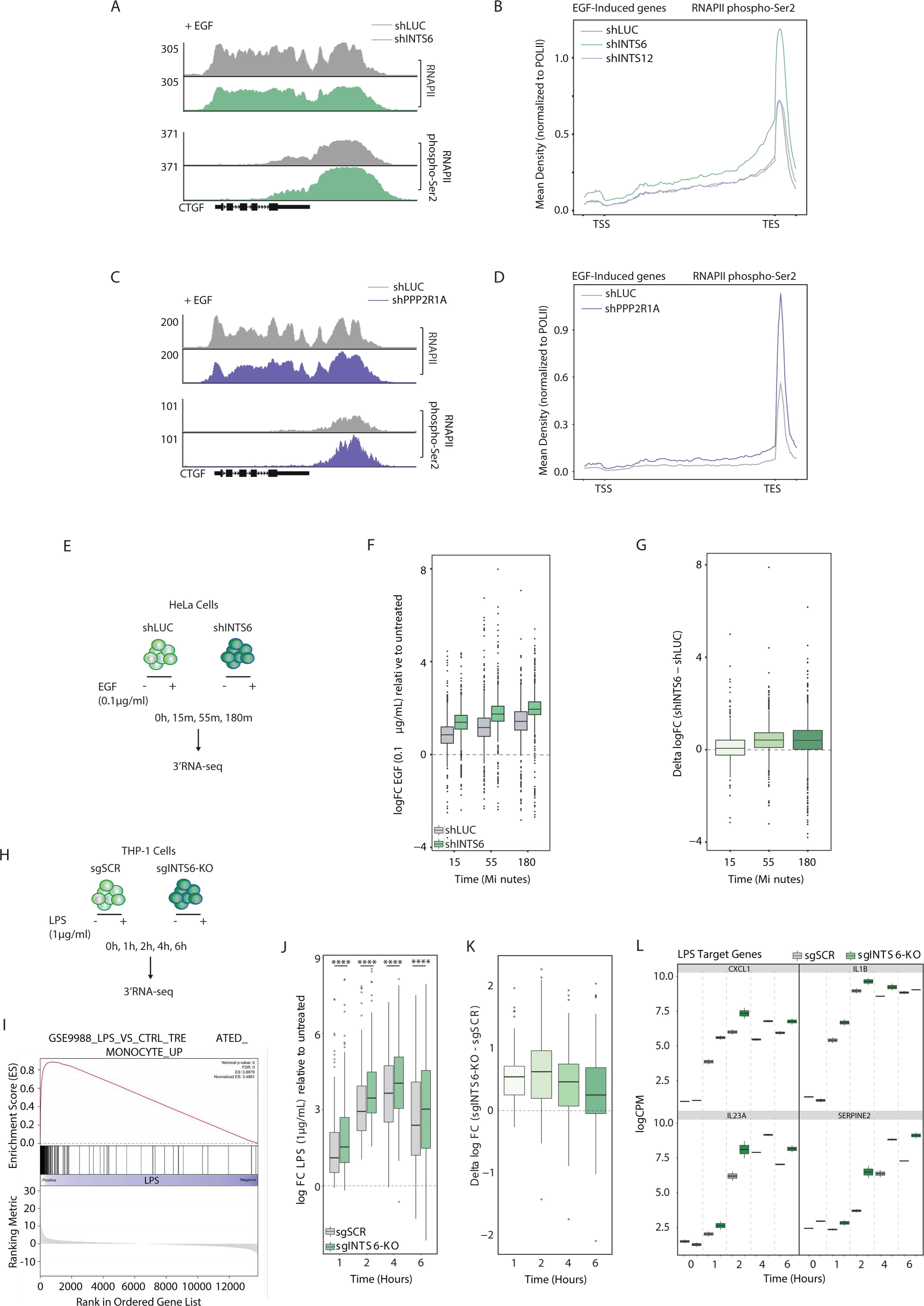
The INTS6/PP2A axis fine-tunes acute transcriptional responses. **A.** RNAPII and RNAPII phospho-Ser2 ChIP-seq signal at the *CTGF* locus after EGF treatment (0.1µg/mL; 15 minutes) in shLUC and shINTS6 infected HeLa cells. **B.** Metagene profile of RNAPII phospho-Ser2 ChIP-seq signal at EGF-response genes (n=50) in shLUC, shINTS6, and shINTS12 infected HeLa cells after EGF treatment (0.1µg/mL; 15 minutes). **C.** RNAPII and RNAPII phospho-Ser2 ChIP-seq signal at the *CTGF* locus after EGF treatment (0.1µg/mL; 15 minutes) in shLUC and shPPP2R1A infected HeLa cells. **D.** Metagene profile of RNAPII phospho-Ser2 ChIP-seq signal under the same conditions at EGF-response genes (n=50). **E.** Overview of acute EGF stimulation (0.1μg/mL; 15, 55, and 180 minutes) of shLUC and shINTS6 infected HeLa cells. **F.** Log fold change (CPM) in EGF-treated versus untreated shLUC and shINTS6 infected HeLa cells at indicated timepoints. **G.** Log fold change difference between shLUC and shINTS6 infected HeLa cells (EGF versus untreated) at indicated timepoints. **H.** Overview of acute LPS stimulation (1μg/mL; 1, 2, 4 and 6 hours) of THP-1-Cas9 sgSCR and sgINTS6-KO cells. **I.** GSEA profile of THP-1-Cas sgSCR cells treated with LPS (2 hours versus 0 hours). **J.** Log fold change (CPM) in LPS-treated versus untreated THP-1-Cas9 sgSCR and sgINTS6-KO cells at indicated time-points. **K.** Log fold change difference between THP-1-Cas9 sgINTS6-KO and sgSCR cells (LPS versus untreated) at indicated time-points. **L.** LPS-target gene expression in THP-1-Cas9 sgSCR and sgINTS6-KO cells at indicated time-points. Figure J was analysed by unpaired, two-sided students t-test, **** p<0.0001.

Examination of steady-state RNA levels by 3’ mRNA-seq (Quantseq) using an extended time course of EGF treatment (Figure 6E) demonstrated that depletion of INTS6 amplified transcriptional activation of EGF-responsive genes, up to 180 minutes after stimulation (Figure 6F and G), an effect also observed in the presence of CDK9i (Figure S6D). Since the EGF-responsive Immediate Early Genes are periodically activated in waves of approximately 30-40 minutes (Amit et al., 2007), these data suggest that human cells do not readily offset loss of the INTS6/PP2A axis and are subjected to persistent transcriptional imbalance.

The role of INTS6/PP2A in regulating transcriptional responses to physiological stimuli was further assessed by 3’ mRNA Quantseq in THP-1 cells with and without stable knockout of INTS6 that were treated over time with LPS (Figure 6H). Canonical LPS gene expression programs were induced in THP-1 cells in response to LPS treatment (Figure 6I, S6C-D), with significantly greater up-regulation of LPS-response genes measured in sgINTS6-KO cells compared to sgSCR controls at all time-points (Figure 6J and K). Expression of key inflammatory LPS-target genes, such as *CXCL1*, *IL1B*, *IL23A,* and *SERPINE2* increased in a time-dependent manner, peaking at 2–4 h post-treatment, with greater levels of gene expression observed in sgINTS6-KO cells under all conditions (Figure 6L). These data provide evidence that PP2A recruitment via INTS6 is required to fine-tune acute transcriptional responses to important pro-inflammatory or growth stimuli.

### Molecular and therapeutic synergy between CDK9 inhibition and PP2A activation

Targeting of CDK9 can be therapeutically efficacious in pre-clinical models of MLL-rearranged leukemia (Baker et al., 2016) and various other hematopoietic malignancies (Gregory et al., 2015), and CDK9i have progressed to clinical trials (Chou et al., 2020). Recently, small molecule activators of PP2A (SMAPs) have been developed, which act by stabilizing PP2A heterotrimers (Kauko et al., 2018; Leonard et al., 2020; Westermarck and Neel, 2020). SMAPs have been demonstrated to be therapeutically effective in combination with MEK inhibitors in KRAS-mutant lung cancer models (Kauko et al., 2018). The mechanistic data detailed above provided a strong rationale to target the CDK9/PP2A controlled pause-release checkpoint via concurrent CDK9 inhibition and SMAP-mediated PP2A enhancement to treat MLL-rearranged leukemias. *In vitro* drug synergy studies were performed using CDK9i and the SMAP DBK-1154 in THP-1 cells. This demonstrated that at EC50 concentrations of CDK9i, DBK-1154 greatly enhanced the induction of cell death in THP-1 and MV4;11 cells, with no single agent activity observed even at high concentrations of DBK-1154 (Figure 7A top panel and Figure S7A-B). Importantly, no synergy was observed between CDK9i and DBK-1154 in sgINTS6-KO THP-1 cells, even at high-dose CDK9 inhibitor treatment, indicating that the therapeutic synergy depends on the INTS6-mediated recruitment of PP2A to the pause-release checkpoint (Figure 7A bottom panel). The synergy between CDK9i and DBK-1154 was also reflected on the transcriptional level with markedly enhanced RNAPII pausing observed in THP-1 cells concurrently treated with both compounds, compared to single agent CDK9i (Figure 7B-C). Consistent with the phenotypic data, DBK-1154 treatment alone had a minimal impact on RNAPII pausing (Figure 7B-C), suggesting that, under steady state conditions, functional compensation is possible to counteract increased PP2A activity.

**FIGURE 7:**
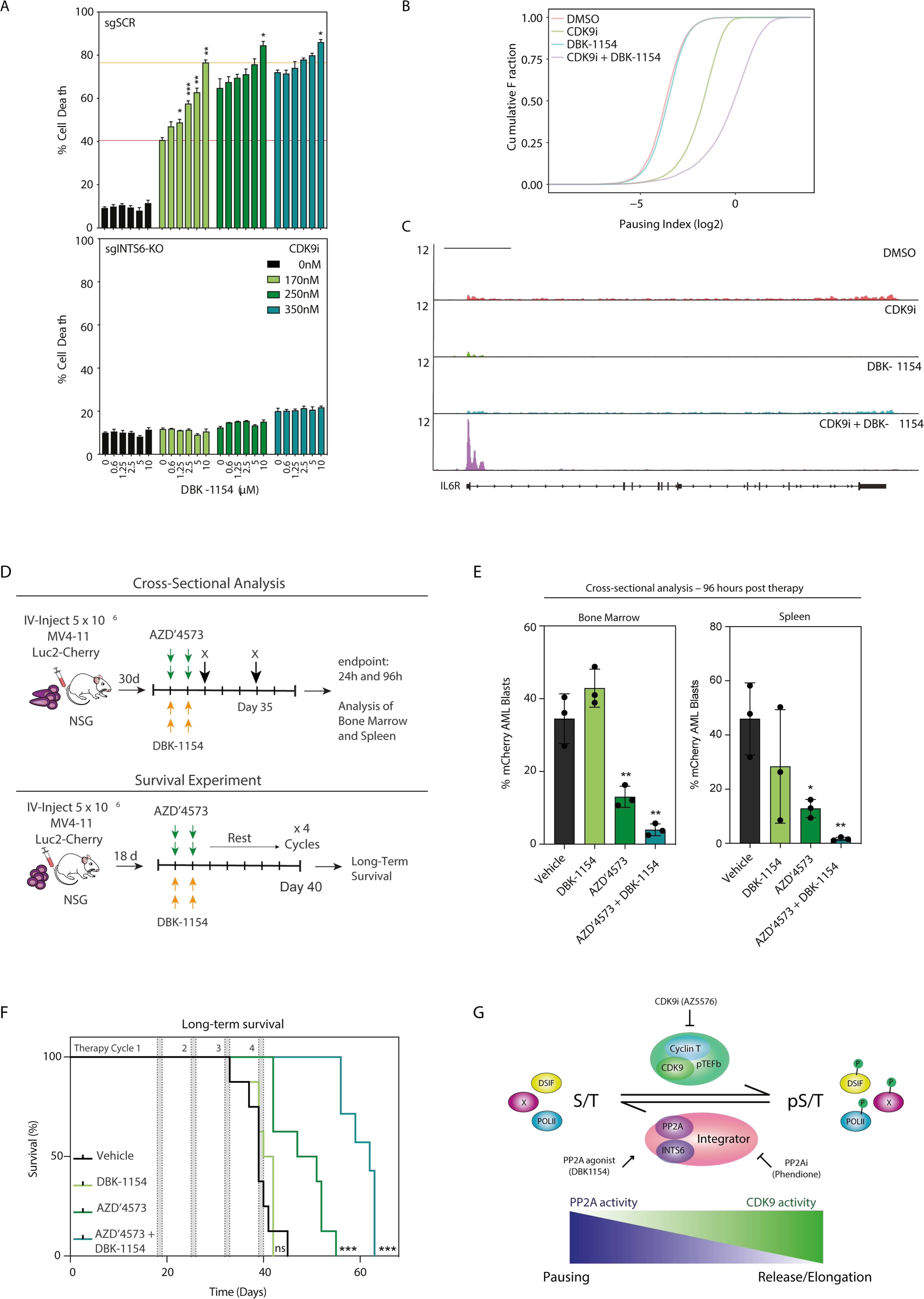
Therapeutic and molecular synergy between PP2A agonist and CDK9i. **A.** Annexin-V analysis of THP-1-Cas9 sgSCR and sgINTS6-KO cells treated as indicated for 72 hours. Orange lines indicate single agent CDK9i activity and maximal synergistic activity with DBK-1154. **B.** Pausing index in THP-1 cells treated with CDK9i (170nM for 2 hours) with or without 15 minutes pre-treatment with DBK-1154 (10μM). **C.** Representative IGV profile for RNAPII ChIP-seq signal at the *IL6R* locus under the same conditions as B. **D.** Schematic of *in vivo* experimental protocol: cross-sectional analysis of acute effects on AML progression and long-term survival in recipients treated with AZD’4573 + DBK-1154 combination therapy. **E.** Quantitation of mCherry-positive AML blasts in bone marrow and spleen at 96 hours post-therapy. **F.** Kaplan-Meier survival curve for long-term survival experiment at the conclusion of 4 rounds of therapy (*n* = 8 mice per group**). G.** Schematic of Integrator/PP2A/CDK9 axis. Figure A is representative of the mean ± SEM of 3 independent experiments. Figure E error bars represent the SD of 3 mice per group and Figure F represents 8 mice per group. Figure A was analyzed by 2-way ANOVA, Figure E was analyzed by Students *t*-Test and Figure F was analyzed by Log-rank (Mantel-Cox) test, * p < 0.05, ** p < 0.01, *** p<0.001. Scale bar for C represents 10kb.

To assess the therapeutic benefits of combined CDK9 inhibition with PP2A agonism *in vivo*, a series of experiments were performed which composed of a cross-sectional analysis that determined the effects of therapy on leukemic progression after acute treatment, and a chronic long-term survival experiment that interrogated whether therapy resulted in survival benefit (Figure 7D). NOD-*scid* IL2Rg^null^ (NSG) mice bearing mCherry/Luciferase-tagged MV4-11 leukemias were treated with the *in vivo*-optimised CDK9 inhibitor AZ4573 (Cidado et al., 2020) and the prototype tricyclic-sulfonamide PP2A activator DBK-1154 (Kastrinsky et al., 2015; Sangodkar et al., 2017). Cross-sectional analysis of mouse bone marrow and spleen 96 hours post-one cycle of combination therapy showed marked reduction in the number of mCherry-positive blast cells relative to single agent and vehicle controls (Figure 7E). These observations corresponded with concomitant reduction in overall spleen size 24 hours post therapy (Figure S7C-D), suggesting that AML blasts rapidly enter apoptosis upon combination treatment and that induction of normal erythroblastic and myeloblastic maturation occurs (Figure S7E). Long-term treatment of mice with four cycles of combination therapy resulted in a significant survival advantage relative to both single agent and vehicle controls (Figure 7F), with mice treated with both compounds yielding smaller spleen sizes at endpoint (Figure S7F). Collectively, these data demonstrate that in the context of *MLL*-rearranged AML, combined inhibition of CDK9 and agonism of PP2A results in marked acute anti-leukemic effects in the bone marrow and spleen, and significant long-term survival advantage in chronically-treated mice.

## DISCUSSION

RNAPII-driven transcription is a stepwise process comprising of distinct checkpoints, each controlled by dedicated transcriptional CDKs and their cognate cyclins (Adelman and Lis, 2012; Mayer et al., 2017). In metazoans, the pause-release checkpoint controlled by pTEFb is believed to be the rate-limiting step in the transcription cycle for most protein coding genes (Guenther et al., 2007). While the basal transcription machinery, including CDK9, is remarkably conserved through evolution, multicellular organisms boast additional multi-subunit regulatory complexes, such as SEC and Integrator, that modulate activity of the basal machinery and coordinate co-transcriptional processes in response to developmental, environmental, and immunological cues (Adelman and Lis, 2012; Baillat et al., 2005; Lai et al., 2015; Luo et al., 2012). In yeast, PP1 has recently been demonstrated to functionally antagonize Cdk9 by dephosphorylating Spt5, to control the elongation-termination transition (Parua et al., 2018). To date, the activity of CDK9 in higher organisms is believed to be unopposed, with phosphorylation of key substrates such as DSIF and the RNAPII CTD resulting in effective pause-release and productive elongation.

The study herein provides the first comprehensive, genome-wide evidence that CDK9 is functionally antagonized in metazoans by the PP2A phosphatase, which is recruited by the RNAPII-associated Integrator complex at the pause-release checkpoint. These data indicate that, in contrast to the prevailing view, CDK9 phospho-substrates are dynamically controlled prior to and during productive transcriptional elongation, providing an additional regulatory layer of transcriptional control (Figure 7G). In addition to its role in transcriptional pause-release, CDK9 has also been demonstrated to control RNAPII termination (Laitem et al., 2015) and RNA processing by phosphorylating substrates such as the exoribonuclease XRN2 (Sanso et al., 2016). It remains to be investigated whether the functional antagonism between CDK9 and PP2A/INTS6 extends beyond the pause-release checkpoint to the final steps of transcription. Although alterations in XRN2 phosphorylation were not detected in our proteomic analysis, the potent CDK9i resistance phenotype conferred by INTS6 depletion suggests that other essential CDK9 controlled pathways are intact in these cells. Recruitment of PP2A extends into the gene body and 3’ end of genes, mirroring RNAPII and suggesting that PP2A controls levels of CTD phosphorylation from end to end. It is noteworthy that, in addition to PP2A, PP1 may play a critical role in mediating the dephosphorylation of CDK9 substrates as has been described in yeast (Parua et al., 2018). In agreement with this notion, PP1 immunoprecipitated with INTS6, and loss of RNAPII and DSIF phosphorylation upon CDK9 inhibition was more potently antagonized when both PP2A and PP1 were inhibited.

The role of PP2A/INTS6 in antagonizing CDK9 seems especially important during acute transcriptional responses to LPS and EGF, as unopposed RNAPII pause-release by CDK9, amplifies the magnitude and duration of pro-inflammatory and mitogenic signals, respectively. It is interesting to speculate that the PP2A/INTS6-mediated negative feedback loop may have co-evolved with the emergence of the CDK9-mediated pause-release in early metazoans, in order to fine-tune transcription in multicellular organisms that have to respond to developmental and environmental cues. Concordantly, the multifunctional Integrator complex first arises in metazoans.

The Integrator complex, comprised of 14 subunits, appears to have a multifaceted role in transcription and RNA processing by coordinating termination of short noncoding RNAs through the INTS11/INTS9 catalytic core (Baillat et al., 2005; Lai et al., 2015; Rienzo and Casamassimi, 2016; Rubtsova et al., 2019). Additionally, Integrator’s catalytic core may help coordinate RNAPII pause-release. Specifically, depletion of the INTS11 subunit can prevent SEC recruitment and productive elongation in human cells (Gardini et al., 2014), whilst the Drosophila Ints9 subunit may impair RNAPII release by prematurely cleaving nascent transcripts at selected genes (Elrod et al., 2019; Tatomer et al., 2019). Data presented herein demonstrated that loss of INTS6 did not affect the catalytic activity of Integrator, nor did it perturb the overall complex integrity. Our data are consistent with a new functional module of Integrator that functionally associates with PP2A. While INTS6 has a central role in recruiting PP2A, functionally related subunits such as INTS8 may also be an integral part of this novel module. The modular nature of Integrator was recently revealed by a study of the INTS13 subunit, and its ability to regulate enhancer activation independent from the core complex (Barbieri et al., 2018). Our data further suggests that Integrator’s activity is the result of an intricate interplay between the core-complex, peripheral modules, and auxiliary members such as the PP2A holoenzyme.

Interestingly, *INTS6* underwent a gene duplication event in mammals with the highly homologous INTS6-like (INTS6L/DDX26B) protein, proposed to localize to mitochondria, whereas INTS6 localizes to the nucleus. Accordingly, both mitochondrial and transcriptional roles have been attributed to the ancestral INTS6 protein in lower organisms (Rienzo and Casamassimi, 2016). In agreement with the disparate roles for INTS6 and INTS6L, our screens revealed no enrichment for loss of *INTS6L*.

Interestingly, INTS6 was originally identified and named for being frequently deleted in cancer 1 (DICE1). In fact, loss or downregulation of INTS6 occurs in prostate cancer, hepatocellular carcinoma, and hematopoietic malignancies (Filleur et al., 2009; Li et al., 2015; Ropke et al., 2005). The tumor suppressive role of INTS6 may be attributed to its proposed role in the sensor of the ssRNA (SOSS) complex, which controls the double stranded DNA damage response (Skaar et al., 2009). However, we propose that INTS6 loss may also promote oncogenic transcription by facilitating pause-release, as is observed in transcriptionally addicted MLL-rearranged leukemias and MYC-driven lymphomas (Bradner et al., 2017; Luo et al., 2012). Reduction of PP2A activity through epigenetic silencing and/or somatic mutations also occurs in several tumors (Perrotti and Neviani, 2013; Shih Ie et al., 2011). Our study provides the first evidence for a role of PP2A in transcription, although it remains to be determined if this activity will significantly contribute to the tumor suppressive function of PP2A.

The recognition that perturbation of core-transcriptional components can fundamentally drive oncogenesis in a variety of cancers has spurred the development of small-molecule inhibitors targeting this process. Indeed, numerous CDK9 inhibitors have demonstrated promising results in pre-clinical studies (Chou et al., 2020). The recent development of PP2A agonists (Clark and Ohlmeyer, 2019) provided the opportunity to explore the molecular and therapeutic synergy between these compounds and CDK9i in MLL-rearranged leukemias where aberrant recruitment of the SEC fuels tumor initiation and progression (Luo et al., 2012). Concurrent PP2A activation and CDK9 inhibition resulted in enforced transcriptional pausing and synergistic cell death *in vitro,* and greatly diminished tumor burden and prolonged survival *in vivo*. These findings demonstrate the therapeutic benefit of dual CDK9/PP2A targeting in cancer and provide the basis for future pre-clinical and clinical studies.

These studies detail the discovery of an auxiliary module of the Integrator complex containing PP2A and INTS6, which is actively recruited to chromatin in response to cellular cues to functionally antagonize CDK9 in multicellular organisms. We posit that antagonism of kinase activity throughout the transcription cycle is critical to fine-tune transcription output, thus challenging the view that RNAPII checkpoints are unilaterally controlled by transcriptional CDKs.

## ACKNOWLEDGEMENTS

We thank members of the Johnstone and Gardini laboratories for helpful input and Prof. Robert Fisher (Icahn School of Medicine at Mount Sinai) for generous supply of anti-phospho SPT5 antibody. The Molecular Genomics Core and the Flow Cytometry Core Facilities at the Peter MacCallum Cancer Center, and the Genomics Core and the Proteomics Core at The Wistar Institute provided excellent technical support. R.W.J was supported by the Cancer Council Victoria, National Health and Medical Research Council of Australia (NHMRC), and The Kids’ Cancer Project. Work from the Gardini Lab (A.G.) was supported by grants from the American Cancer Society (RSG-18-157-01-DMC), the NIH (R01 HL141326), and the G Harold & Leyla Y Mathers Foundation (A.G.). S.J.V. was supported by a Rubicon fellowship (NWO, 019.161LW.017), NHMRC EL1 fellowship (GNT1178339), and The Kids’ Cancer Project. S.A.W. was supported by a training grant NIH T32-GM071339. J.R.D was supported by an Early Career Seed Grant from the Victorian Cancer Agency (VCA). S.J.H. was supported by a Postdoctoral Fellowship from the Cancer Council of Victoria (CCV). K.F.H was supported by a NHMRC Senior Research Fellowship (APP1078220). J.W and K.P were supported by Academy of Finland (294850) and Finnish Cancer Foundation and K.P. was also supported by Finnish Cultural Foundation. The Victorian Centre for Functional Genomics ACRF Translational Reverse Phase Protein Array platform (K.J.S.) is funded by the Australian Cancer Research Foundation (ACRF), University of Melbourne Collaborative Research Infrastructure Program, and the Peter MacCallum Cancer Centre Foundation. The Peter MacCallum Foundation and Australian Cancer Research Foundation provide generous support for equipment and core facilities.

## AUTHOR CONTRIBUTIONS

S.J.V. performed experiments, next generation sequencing analysis, data interpretation, and provided study supervision. S.A.W. performed experiments, next generation sequencing analysis and data interpretation. J.R.D., D.A.K., E.B., I.T., C.J.K., M.C., S.B., Z.F., J.H.A.V., B.P.M., I.Y.K., S.J.H., M.J.K. and A.N performed experiments, data analysis, next generation sequencing analysis, and data interpretation. J.J.S. performed Mass spectroscopy. K.P., O.K., K.F.H., M.O., E.D.H., J.W. K.J.S., G.G. and N.G. provided critical reagents and advice. R.W.J. and A.G. contributed to data interpretation and study supervision. S.J.V., S.A.W, J.R.D., A.G and R.W.J. wrote the manuscript, which was proof read and edited by all co-authors prior to submission.

## METHODS

### Cell Lines

THP-1 (ATCC^®^ TIB-202™) and MV4;11 (ATCC^®^ CRL-9591™) cells were cultured at 37°C and 5 percent carbon dioxide (CO_2_) in Roswell Park Memorial Institute medium (RMPI; ThermoFisher Scientific 11875093) supplemented with 10-20% fetal bovine serum (FBS; Peter MacCallum Cancer Centre), 100U/mL penicillin, 100μg/mL streptomycin (ThermoFisher Scientific 10378016) and 2mM GlutaMAX™ (ThermoFisher Scientific 35050061). Henrietta Lacks (HeLa - ATCC^®^ CCL-2™) cells were cultured at 37°C and 5 percent carbon dioxide (CO_2_) in Dulbeccos Modified Eagle’s medium (DMEM) supplemented with 10% super calf serum (GEMcell) and 2mM L-glutamine (Corning). *Drosophila melanogaster* (*D.melanogaster*) S2 cells (ATCC^®^ CRL-9591 ™) were cultured in Scheinder’s *Drosophila* medium (ThermoFisher Scientific 21720) supplemented with 10% HI-FBS, 100U/mL penicillin; 100μg/mL streptomycin and 2mM GlutaMax™ at room temperature and atmospheric CO_2_. Human embryonic kidney (HEK) 293T cells (ATCC^®^ CRL-11268™) were used for generation of lentivirus and retrovirus and were cultured at 37°C and CO_2_ in Dulbeccos Modified Eagle’s medium (DMEM; ThermoFisher Scientific 11995073) supplemented with 10% FBS, 100U/mL penicillin; 100μg/mL streptomycin and 2mM GlutaMax™.

### Reagents

AZ’5576 (CDK9i) and AZD’4573 were provided by AstraZeneca, and reconstituted in dimethyl sulfoxide (DMSO) for *in vitro* use. For *in vivo* use AZD’4573 as reconstituted in a 10:30:70 ratio of N,N-dimethylacetamide (DMA), PEG-400, and 1% Tween-80 in H_2_O. Fresh AZD’4573 stocks were prepared weekly. DBK-1154 was provided by Michael Ohlmeyer (Icahn School of Medicine, Mount Sinai Hospital) and was reconstituted in DMSO (*in vitro*) or prepared daily in 10% DMA, 10% Kolliphor-HS-15 and 80% H_2_O (*in vivo*). The CDK9 degrader, 22-533, and THZ1 were provided by Nathaneal Gray and were reconstituted in DMSO. LPS was purchased from Sigma Aldrich (L2630) and reconstituted in PBS. Phendione (1,10-Phenanthroline-5,6-dione) was purchased from Sigma Aldrich (496383) and reconstituted in DMSO. Calyculin A (ab141784) and Okadaic Acid (ab120375) were purchased from Abcam and reconstituted in DMSO. Cytarabine was obtained from the Peter MacCallum Cancer Centre pharmacy. EGF was purchased from Gibco (PHG0311) and reconstituted in PBS. Flavopiridol was purchased from Sigma-Aldrich (F3055) and reconstituted in DMSO.

### Click-EU labelling/nascent RNA

Cells treated with transcriptional inhibitors were labelled with 1mM ethynyl-uridine (EU, ThermoFisher Scientific E10345) for 1 hour prior to fixation/permeabilization (0.5% paraformaldehyde, 0.2% Tween20, 0.1% BSA, 0.1% Azide). Cells were washed once with PBS and incubated at room temperature for 30 minutes with the Click-iT reaction buffer, CuSO_4_ buffer, Alexa-Fluor-azide-647 and Click-iT reaction buffer additive from the Click-iT^®^ RNA Imaging kit as per the manufacturer’s instructions (ThermoFisher Scientific, C10329). Cells were washed twice with Click-iT kit rinse buffer, and once each with PBS plus 0.1% Tween 20 and fixation/permeabilization buffer prior to analysis using the BDFortessa flow cytometer.

### Genome-wide CRISPR-Cas9 screens

THP-1 and MV4;11 cells were engineered to stably express humanized *S.pyogenes* Cas9 endonuclease by lentiviral transduction with the FUCas9Cherry vector (Addgene 70182) and subsequent FACS-selection for mCherry-positive cells (THP-1-Cas9, MV4;11-Cas9). For THP-1-Cas9 survival screens, cells were transduced with the GeCKOV2 (125000 sgRNAs) (Sanjana et al., 2014) or Brunello (75000 sgRNAs) (Doench et al., 2016) genome-wide sgRNA libraries at a MOI of 0.3 and a fold representation of 500 for each individual sgRNA. For nascent RNA screens THP-1-Cas9 cells were transduced with the Brunello sgRNA library. Transduced cells were selected with puromycin (1μg/mL) for 7 days, at which time cells were split into relevant treatment conditions (DMSO, CDK9i) and cell pellets were collected and snap-frozen for T_0_ reference controls. For survival screens, puromycin-selected cells were cultured in AZ-5576 (150nM), A159 (50nM, GeCKOV2 only) or Dinaciclib (10nM, GeCKOV2 only) for 21 days to select for CDK9i-resistant populations and cell pellets were collected from DMSO- and CDK9i-resistant populations (Tend). For nascent RNA screens THP-1-Cas9 cells were transduced with the Brunello sgRNA library and puromycin-selected cells were cultured for 16 hours with DMSO or AZ-5576 (170nM). Cells were labelled for 1 hour with EU and Click-iT RNA detection was performed as described previously prior to FACS-selection of EU-positive cells using a BDFACSAria sorter. For MV4;11-Cas9 survival screens, cells were transduced with the Brunello sgRNA library and puromycin-selected cells were cultured for 21 days with DMSO or 350nM AZ-5576. Genomic DNA was isolated from T_0_, Tend and EU-positive cells using the DNeasy Blood and Tissue Kit (Qiagen, 69504). Sequencing libraries were generated by PCR amplification as previously described (Joung et al., 2017; Kearney et al., 2018) and were 75 base-pair single-end reads were sequenced using the Illumina NextSeq500. Sequencing files were demultiplexed (Bcl2fastq, v2.17.1.14), reads were trimmed to 20 base-pair sequences using cutadapt (v2.1) and MAGeCK (v0.5.9) was used to count reads and perform sgRNA enrichment analysis. The R package ggplot2 (v3.3.0) was employed for data visualization of screens. Venn diagrams were generated using the online web tool BioVenn (Hulsen et al., 2008); represented genes had an adjusted p-value of < 0.1 for > 3 sgRNAs in at least one replicate screens. Only genes with an adjusted p-value less than for *INTS6, INTS8* or *INTS12* were included.

### Stable expression of full-length INTS6 in THP-1 MLL-AML cells

Human full-length INTS6 ORF in pCMV6-entry vector was purchased from Origene (RC208036). The INTS6 ORF was PCR amplified with 5’ Xho I and 3’ EcoRI restriction site overhangs and the coding sequence for a C-terminal V5 epitope tag was added in frame to the 3’ end of the INTS6 ORF (Key Resources Table. The INTS6 PCR product and MSCVpuro vector (Addgene 68469) were digested with XhoI (NEB R0146) and EcoRI-HF (NEB R3101) at 37°C overnight and purified digests were ligated overnight at 16°C using T7 DNA ligase as per manufacturer’s instructions (NEB M0318). Ligation reactions were transformed into One Shot™ Stbl3™ chemically-competent *E.coli* cells (42°C heat-shock, 90 seconds; Invitrogen C737303) and positive clones were selected by Sanger sequencing. THP-1-Cas9 cells expressing FgH1t-UTG-sgSCR-GFP or FgH1tUTG-sgINTS6-GFP with stable knockout of INTS6 were engineered to express empty vector MSCVpuro or MSCVpuro-V5-INTS6 by retroviral transduction.

### Generation of mCherry-P2A-V5-PPP2R1A knock-in THP-1 cells

5 × 10^5^ THP-1 cells were resuspended in 20µL nucleofection solution (16.4µL SG nucleofector solution + 3.6µL supplement 1) from the SG Cell Line 4D-Nucleofector™ X kit (Lonza, V4XC-3032). 300pmol PPP2R1A sgRNA (Synthego, Key Resources Table) was incubated with 100pmol Alt-R^®^ S.p. HiFi Cas9 Nuclease V3 (Integrated DNA Technologies, 1081061) at room temperature for 20 minutes prior to the addition of 566ng mCherry-P2A-V5-PPP2R1A donor template DNA (Integrated DNA Technologies, Key Resources Table) on ice (6µL total volume). THP-1 cells in SG buffer were mixed with the sgRNA/Cas9/template complex and transferred to a 16-well nucleocuvette strip. Nucleofection was performed using the FF100 program of the Amaxa 4D Nucleofector X Unit (Lonza, AAF-1002X) and cells were incubated at 37°C for 10 minutes post-nucleofection before addition of culture media. Following expansion, mCherry cells were FACS-selected using a BD FACSAria™ Fusion sorter.

### Generation of *D.melanogaster* S2-Cas9 cells

Control (SCR) and *IntS6* targeting sgRNAs (Sigma Aldrich, Key Resources Table) with BspQ1 overhangs were ligated using T7 DNA ligase (NEB M0318; 16°C overnight) into BspQ1 digested (NEB R0712) pAc-sgRNA-Cas9 vector (Addgene 49330) (Bassett et al., 2014). *D.melanogaster* S2 cells were transfected with pAc-sgSCR-Cas9 or pAc-sg*IntS6*-Cas9 by Effectene transfection (Qiagen 301425) and positive clones were selected with puromycin.

### Expression of full-length Ints6-SBP in *D. melanogaster* S2 cells

The *D. melanogaster IntS6* coding sequence (FlyBase.org, Clone DmeI\SD04165. FBcl0286688) was PCR amplified with 5’ XhoI and 3’ KpnI restriction site overhangs. The *IntS6* PCR product and pMK33-SBP C-terminal vector (Yang and Veraksa, 2017) were digested with XhoI and KpnI-HF (NEB R3142) at 37°C overnight and purified digests were ligated overnight at 16°C. Ligation reactions were transformed into DH5-α chemically-competent cells (42°C heat-shock, 45 seconds; Invitrogen 18265017) and ampicillin-resistant clones were screened by sanger sequencing. *D.melanogaster* S2 clones expressing SCR sgRNA or with stable knockout of Ints6 (sg*IntS6*-5, clone D and clone E) were engineered to express pMK33-*IntS6*-SBP or pMK33-SBP by Effectene transfection and positive clones were selected using hygromycin (Yang and Veraksa, 2017) (300μg/mL; ThermoFisher Scientific 10687010). Expression of IntS6-SBP was confirmed by western blot (SBP-Tag antibody, Key Resources Table).

### Competitive Proliferation and Cell Death Assays

For THP-1/MV4;11 competitive proliferation assays 2 independent INTS6 sgRNAs or a control Scrambled (SCR) sgRNA (Sigma Aldrich, Key Resources Table) with BsmBI compatible overhangs were ligated into BsmBI (NEB R0739) digested FgH1tUTG-GFP vectors (Addgene 70183). THP-1-Cas9 and MV4;11-Cas9 cells were engineered to stably express FgH1tUTG-sgINTS6-A-GFP, FgH1tUTG-sgINTS6-B-GFP or FgH1t-UTG-sgSCR-GFP vectors by lentiviral transduction. Cells were incubated with doxycycline (1μg/mL; Sigma Aldrich D9891) for 7 days and depletion of INTS6 protein was confirmed by SDS-PAGE and western blotting. THP-1/MV4;11-Cas9 cells expressing FgH1tUTG-sgINTS6-GFP or FgH1t-UTG-sgSCR-GFP were mixed 1:1 with THP-1/MV4;11-Cas9 cells expressing FgH1t-UTG-sgSCR-BFP, incubated with doxycycline (1μg/mL) and cultured in the presence of DMSO, CDK9i or the CDK9-degrader at indicated concentrations. The relative proportions of GFP- and BFP-positive cells was measured following mixing (T_0_) and at 4, 7, 11 and 14 days post-assay initiation using a BD LSRFortessa™ flow cytometer. For HeLa competition assays, HeLa cells stably expressing either the FUCas9Cherry or FUCas9CFP vector were electroporated with 100pmol Alt-R^®^ S.p. HiFi Cas9 Nuclease V3 and 300pmol *INTS6*-A targeting sgRNA or sgSCR (Synthego, Key Resources Table) respectively using the Amaxa 4D Nucleofector X Unit as per manufacturer’s instructions. At five days post-electroporation mCherry and CFP Cells were mixed 1:1 and incubated with increasing concentrations of AZ-5576 for 96 hours. Cell populations were assessed by flow-cytometry using a BD LSRFortessa™ and SDS-PAGE/immunoblotting. For cell death analysis, THP-1-Cas9 cells expressing FgH1tUTG-sgINTS6-A-GFP, FgH1tUTG-sgINTS6-B-GFP or FgH1t-UTG-sgSCR-GFP or *D.melanogaster* S2 cells expressing pAc-sgSCR-Cas9 or pAc-sg*Ints6*-Cas9 were incubated with increasing doses of AZ-5576, cytarabine, THZ1 or DBK-1154 for 72 hours. Cells were stained with Annexin-V APC (BD Biosciences 550475) prior to flow cytometric analysis using a BD LSRFortessa™. The R package synergyfinder (v1.8.0) (He et al., 2018) was employed for Zero Interaction Potency (ZIP) synergy score computing between DBK-1154 and AZ-5576.

### Infection of HeLa cells with shRNA

shRNA vectors pLKO.1-shINTS6, pLKO.1-shINTS8, pLKO.1-shINTS12, and pLKO.1-shPPP2R1A were obtained from the Molecular Screening Facility at the Wistar Institute. shINTS2 and shINTS5 were designed with the Broad Institute algorithm (https://portals.broadinstitute.org/gpp/public/) and subsequently cloned into pLKO.1 (Addgene #10879). Sequences of all shRNAs are listed in the Key Resources Table. For infection, lentiviral particles were produced in HEK293T cells with pLKO.1-shRNA vectors as indicated. HeLa cells were incubated overnight in growing media plus virus and polybrene (Sigma-Aldrich, TR-1003). The following day, the virus-containing media was removed and replaced with growing media, and cells were selected with puromycin (Invivogen) at 2μg/mL. In all shRNA experiments, cells were treated and harvested within 72 hours of infection.

### SDS-PAGE and Immunoblotting

For whole cell THP-1 lysates, cells were washed once with cold PBS and lysed with laemmli lysis buffer (60mM Tris-HCl pH 6.8, 10% (v/v) glycerol, 2% (w/v) SDS) for 10 minutes at 95°C and protein concentration was determined using the Pierce™ BCA Protein Assay Kit (ThermoFisher Scientific 23227). Protein samples were separated by SDS-PAGE using 4-20% Mini-PROTEAN^®^ TGX™ gels (BioRad, #4561094) in 25mM Tris, 192mM Glycine, 0.1% (w/v) SDS and wet-transferred to Immobulin-P (Merck Millipore, IPVH00010) or Immobulin-FL membranes (Merck Millipore, IPFL00010) in 25mM Tris, 192mM Glycine, 5% v/v methanol (4°C, 200mA, 2 hours). Membranes were blocked at room-temperature in Tris-buffered-saline (TBS) supplemented with 0.1% (v/v) Tween20 (TBST; Sigma Aldrich, P1379) and 5% skim-milk powder (local supermarket) or Odyssey® blocking buffer (LI-COR, 927-50000) prior to incubation at 4°C overnight with primary antibodies (Key Resources Table). Membranes were washed 3 times in TBST and incubated for 1 hour at room temperature with HRP-conjugated or LI-COR secondary antibodies (Key Resources Table). Membranes were washed 3 times in TBST and incubated with Amersham ECL PLUS™ (GE Healthcare, RPN2132) prior to exposure to Super RX film (Fujifilm, 03G01), or images were acquired using the Odyssey CLx machine (LI-COR, 83µm, medium setting).

For whole cell HeLa lysates, cells were harvested and washed three times in 1X PBS and lysed in ChIP lysis buffer (150 mM NaCl, 1% Triton X-100, 0.7% SDS, 500 mM DTT, 10 mM Tris-HCl, 5 mM EDTA) supplemented with 1 µg/mL aprotinin, 1 µg/mL leupeptin (Sigma) and 1 µg/mL pepstatin (BMB). Protein samples were loaded into Bolt 4-12% Bis-Tris Plus gels (Invitrogen) and separated through gel electrophoresis (SDS-PAGE) in Bolt MES running buffer (Invitrogen). Separated proteins were wet-transferred to ImmunBlot PVDF membranes (BioRad) for antibody probing. Membranes were incubated with 10% BSA in TBST for 30 minutes at room temperature, then incubated for 2 hours at room temperature or overnight at 4°C with the suitable antibodies diluted in 5% BSA in 1X TBST. After incubation with the primary antibody, the membranes were washed with TBST, and incubated with a 1:10000 dilution of HRP-linked anti-mouse or anti-rabbit secondary antibody (Cell Signaling) for one hour at room temperature. HRP-linked antibodies were visualized using Clarity Western ECL substrate (Biorad) and imaged with Fujifilm LAS-3000 Imager (Fujifilm).

### Co-immunoprecipitation experiments

For Co-IP experiments in THP-1 cells this protocol adapted from (Gregersen et al., 2019) was used. 10 × 10^6^ cells were used per IP and all lysis and wash buffers were supplemented with Roche cOmplete™ protease inhibitors (Merck, 11873580001) and Pierce phosphatase inhibitors (ThermoFisher Scientific, A32957). Cells were washed twice with cold PBS and resuspended in 2 pellet volumes of hypotonic lysis buffer (10mM HEPES pH 7.5, 10mM KCl, 1.5mM MgCl_2_) prior to incubation on ice for 40 minutes. Nuclei were pelleted at 4°C (3900rpm, 15 minutes) and the hypotonic cytoplasmic fraction was removed and discarded. Nuclei were resuspended in 2 original pellet volumes of nucleoplasmic lysis buffer (20mM HEPES pH 7.9, 1.5mM MgCl_2_, 150mM potassium acetate, 10% (v/v) glycerol, 0.05% (v/v) IGEPAL^®^ CA-630) and incubated on ice for 20 minutes. Chromatin was pelleted at 4°C (20kg, 20 minutes) and the nucleoplasmic fraction was removed and stored on ice. Chromatin was incubated with 2 original pellet volumes of low salt chromatin digestion buffer (20mM HEPES pH 7.9, 1.5mM MgCl_2_, 10% (v/v) glycerol, 150mM NaCl, 0.1% (v/v) IGEPAL^®^ CA-630, 250 U/mL benzonase (Merck, 70746-4)) on ice for 1 hour prior to centrifugation at 4°C (20kg, 20 minutes). The low salt chromatin fraction was removed and stored on ice. The remaining undigested chromatin was incubated with 2 original pellet volumes of high salt chromatin digestion buffer (20mM HEPES pH 7.9, 3mM EDTA, 1.5mM MgCl_2_, 10% (v/v) glycerol, 500mM NaCl, 0.1% (v/v) IGEPAL^®^ CA-630) on ice for 20 minutes. 6 original pellet volumes of salt dilution buffer (20mM HEPES pH 7.9, 3mM EDTA, 1.5mM MgCl_2_, 10% (v/v) glycerol, 0.1% (v/v) IGEPAL^®^ CA-630) was added prior to centrifugation at 4°C (20kg, 20 minutes). The nucleoplasmic, low salt chromatin and diluted high salt chromatin fractions were pooled and input samples taken. Pooled fractions were tumbled at 4°C for 1 hour with 5μg of antibody (Key Resources Table) and 25μL Dynabeads™ Protein A or Protein G (ThermoFisher Scientific, 10001D and 10003D respectively). Beads were washed 5 times (20mM Tris-HCl pH 7.5, 150mM NaCl, 1.5mM MgCl_2_, 3mM EDTA, 10% (v/v) glycerol, 0.1% (v/v) IGEPAL^®^ CA-630) and incubated for 10 minutes at 95°C in 2X laemmli lysis buffer (120mM Tris-HCl pH 6.8, 4% (w/v) SDS, 1% β-mercaptoethanol, 0.02% (w/v) bromophenol blue) to elute immunoprecipitated proteins. Eluted fractions were analyzed by SDS-PAGE and immunoblotting as described previously.

For Co-IP experiments in HeLa, cells were washed twice with ice cold PBS before resuspension in buffer A (10mM Tris pH 7.9, 1.5mM MgCl2, 10mM KCl, 0.5mM DTT, 1 mg/ml each of protease inhibitors aprotinin, leupeptin, and pepstatin), and incubated at 4°C for 10 minutes. After spinning down at 2000rpm for 10 minutes, the pellet was resuspended in buffer A and subject to dounce homogenization (with B pestle), and spun down again. The supernatant was kept as cytoplasmic extract. The pellet was resuspended in buffer C (20mM Tris pH 8.0, 1.5mM MgCl2, 0.42M NaCl, 25% glycerol, 0.2mM EDTA, 0.5mM DTT, protease inhibitors) and dounce homogenized (with B pestle), followed by incubation at 4°C for 30 minutes. The resulting lysate was spun down at 12,000rpm for 30 minutes. The supernatant was kept as nuclear extract. The chromatin pellet was resuspended in Nuclease Incubation buffer (150mM HEPES pH 7.9, 1.5mM MgCl2, 150mM potassium acetate, 10% glycerol, 0.5mM DTT, and protease inhibitors) with Benzonase nuclease (Sigma-Aldrich), and incubated for 45 minutes at 4°C. The supernatant was kept as chromatin extract following centrifugation at 12,000rpm for 30 minutes. All extracts were dialyzed overnight in BC80 (20mM Tris pH 8.0, 80mM KCl, 0.2mM EDTA, 10% glycerol, 1mM B-mercaptoethanol, 0.2mM phenylmethylsulfonyl fluoride (PMSF)), cleared, and stored at −80°C. Prior to IP, 500ug of nuclear or chromatin extract was diluted in co-IP buffer (20mM Tris pH 7.9, 100mM NaCl, 0.1% NP-40, protease inhibitors). Each IP was incubated for 2 hours at 4°C with 2μg antibody (Key Resources Table) and 25μL of Dynabeads™ Protein A or Protein G. Beads were washed three times with co-IP buffer, followed by a final wash with 0.05% NP-40 in PBS. Elution was performed by agitation in 0.1M glycine pH 3.0 for one minute, and 1M Tris base pH 11.0 was added to neutralize the pH of the eluate. IP elutions were analyzed by SDS-PAGE and immunoblotting as described previously.

### Immunoprecipitation-mass spectrometry (IP-MS)

For THP-1 and MV4;11 IP-MS experiments 50-80 × 10^6^ cells were used per IP. Dynabeads™ Protein A or Protein G (Invitrogen 10002D and 10003D) were washed 5 times with 5% (w/v) BSA/PBS and tumbled at 4°C with relevant antibodies (Key Resources Table) for 1 hour. Cells were washed twice with cold PBS and incubated on 4°C roller for 10 minutes in nuclear extraction buffer (10mM HEPES, 10mM KCl, 1.5mM MgCl-_2_, 0.05% (v/v) IGEPAL^®^ CA-630, protease and phosphatase inhibitors) prior to centrifugation at 4°C (2000g, 5 minutes). This step was repeated once before isolated nuclei were incubated in 1mL mass spectrometry buffer (20mM Tris-HCl pH 8, 150mM NaCl, 2mM EDTA, 0.5% (v/v) IGEPAL^®^ CA-630, protease and phosphatase inhibitors) on ice for 10 minutes prior to sonication using the Covaris S220 Focused Ultrasonicator (8 minutes per sample). Sonicated lysates were cleared at 4°C (20000g, 10 minutes) prior to tumbling at 4°C with 50μL antibody-bound Dynabeads™ Protein A or Protein G for 1 hour. For IP-MS experiments for SBP or SPB-Ints6 in *D.melanogaster* S2 cells expressing pAc-sgSCR-Cas9 and pMK33-SBP or pMK33-IntS6-SBP, 100 × 10^6^ cells were used per IP using 50μL Pierce™ Strepavidin Agarose slurry (ThermoFisher Scientific, 20353) under the same conditions. Beads were washed 5 times (20mM Tris-HCl pH 8, 150mM NaCl, 2mM EDTA, 0.1% (v/v) IGEPAL^®^ CA-630) and 2 times in 100mM ammonium bicarbonate (AMBIC). Beads were resuspended in 50μL 100mM AMBIC and incubated overnight at 37°C with 400ng Trypsin (Promega V5280). Beads were supplemented with an additional 400ng Trypsin and incubated for a further 4 hours at 37°C. The peptide-containing supernatant was isolated from the beads and acidified through the addition of 10% formic acid. Peptides were isolated using C18 ultra micro-spin columns (Pierce #87782) which had been pre-conditioned (100% acetonitrile followed by 2 rounds of 80% acetonitrile) and equilibrated (0.5% formic acid). Columns were washed twice (0.1% formic acid) and peptides were eluted using 40DL 80% acetronitrile diluted in 0.1% formic acid. Excess acetonitrile was evaporated using a vacuum centrifuge and samples were analysed at the Bio21 Mass Spectrometry and Proteomics facility (Parkville, VIC 3052, Australia). LC MS/MS was carried out on a QExactive plus Orbitrap mass spectrometer (Thermo Scientific) with a nanoESI interface in conjunction with an Ultimate 3000 RSLC nanoHPLC (Dionex Ultimate 3000). The LC system was equipped with an Acclaim Pepmap nano-trap column (Dinoex-C18, 100 Å, 75 µm × 2 cm) and an Acclaim Pepmap RSLC analytical column (Dinoex-C18, 100 Å, 75 µm × 50 cm). The tryptic peptides were injected to the enrichment column at an isocratic flow of 5 µL/min of 2% v/v CH3CN containing 0.1% v/v formic acid for 5 min applied before the enrichment column was switched in-line with the analytical column. The eluents were 0.1% v/v formic acid (solvent A) and 100% v/v CH3CN in 0.1% v/v formic acid (solvent B). The flow gradient was (i) 0-6min at 3% B, (ii) 6-40 min, 3-25% B (iii) 40-48 min, 25-45% B (iv) 48-50 min, 40-80% B (v) 50-53 min, 85-85% B (vi) 53-54 min 85-3% and equilibrated at 3% B for 10 minutes before the next sample injection. The QExactive plus mass spectrometer was operated in the data-dependent mode, whereby full MS1 spectra were acquired in positive mode, 70 000 resolution, AGC target of 3e6 and maximum IT time of 50ms. Fifteen of the most intense peptide ions with charge states ≥2 and intensity threshold of 1.7e4 were isolated for MSMS. The isolation window was set at 1.2m/z and precursors fragmented using normalized collision energy of 30, 17 500 resolution, AGC target of 1e5 and maximum IT time of 100ms. Dynamic exclusion was set to be 30sec. Raw files were analyzed using MaxQuant (version 1.5.8.3). The database search was performed using the Uniprot *Homo sapiens* or *Drosophila melanogaster* databases plus common contaminants with strict trypsin specificity allowing up to 3 missed cleavages. The minimum peptide length was 7 amino acids. Carbamidomethylation of cysteine was a fixed modification while N-acetylation of proteins N-termini and oxidation of methionine and phosphorylation of Serine/Threonine/Tyrosine were set as variable modifications. The iBAQ quantification option was selected. During the MaxQuant main search, precursor ion mass error tolerance was set to 4.5 ppm and fragment ions were allowed a mass deviation of 20 ppm. PSM and protein identifications were filtered using a target-decoy approach at a false discovery rate (FDR) of 1% with the match between runs option enabled. The RStudio package ggplot2 (v3.3.0) was employed for data visualization: a value of 10 was added to all raw iBAQ scores to allow log10 transformation. The average value of log10(iBAQ) of each technical replicate for each condition in THP1 and MV4;11 was calculated and subsequently filtered according the following criteria: log10(isotype) < 2 & log10(IP) > 2. Interaction networks were visualized in Rstudio (v3.5.1) using the packages concaveman (v1.0.0), dplyr (v0.8.1), ggraph (v2.0.3) and ggforce (v0.3.1).

For HeLa cell IP-MS experiments, IPs were performed as described previously, but with the following modifications: for each IP, 1-2mg of nuclear or chromatin extract was incubated with 4μg antibody (Key Resources Table) and 50μL Dynabeads™ Protein A or Protein G. Eluates were prepared for SDS-PAGE and run on a Novex WedgeWell 10% Tris-Glycine Gel (Invitrogen) with Tris-Glycine-SDS buffer (Bio-Rad), at 110V for 10 minutes. The gel was stained with Colloidal Blue staining kit (Invitrogen), and further processed at the proteomics facility at the Wistar Institute. Briefly, the gel lanes were excised, digested with trypsin, and analyzed by LC-MS/MS on a Q-Exactive Plus mass spectrometer. MS/MS spectra generated from the LC-MS/MS runs were searched with full tryptic specificity against the UniProt human database (www.uniprot.org; 10/01/2018) using the MaxQuant 1.6.2.3 program. False discovery rates for protein and peptide identifications were set at 1%. The RStudio package ggplot2 (v3.3.0) was used for data visualization: a value of 10 was added to all raw iBAQ scores to allow log10 transformation.

### qRT-PCR

HeLa cells were lysed in Tri-reagent, followed by RNA extraction using Direct-zol RNA MiniPrep kit (Zymo Research). Reverse transcription of 900ng of template RNA into cDNA was done using random-hexamer primers with the Revertaid first strand cDNA synthesis kit (Thermo Scientific), according to manufacturer instructions. Real-time quantitative PCR reaction was performed with 50ng cDNA, 0.4mM primer, 10μL iQ SYBR Green Supermix (BioRAD), in a final volume of 20μL. We used the CFX96 real-time system (BioRAD), and thermal cycling parameters were as follows: 3 minutes at 95°C, followed by 40 cycles of [10s at 95°C, 20s at 63°C, 30 s at 72°C]. All samples were run in triplicate. 18S rRNA used to normalize. Primer sequences are reported in Key Resources Table.

### Glycerol gradient ultracentrifugation

Glycerol gradients were prepared fresh with 1X HEMG buffer (50mM HEPES (pH 7.9), 0.2mM EDTA, 30mM MgCl2, 200mM KCl, 0.5mM DTT, 1 mg/ml each of protease inhibitors aprotinin, leupeptin, and pepstatin) and glycerol from 11% to 50% in increments of 3%. Gradients were assembled on ice by carefully layering 320uL of 1X HEMG-glycerol in descending order of glycerol concentration, starting with 50%, into an ultracentrifuge tube (Beckman Coulter 344057). Prior to centrifugation, 500ug of nuclear extract (brought to 300uL) was layered on top. Gradients were ultracentrifuged in a swinging-bucket rotor at 40,000rpm for 16 hours (brake=1, accel=9). After centrifugation, gradient fractions were collected by piercing a hole in the bottom of the tube and collecting fractions of 320uL each. Protein was precipitated from each fraction with 1/10th volume trichloroacetic acid (TCA, Sigma-Aldrich), incubated overnight at −20°C, washed twice with cold acetone (Sigma-Aldrich), and resuspended in 70uL of 1X BOLT laemmli sample buffer (Invitrogen) for SDS-PAGE analysis.

### Reverse Phase Protein Array

THP-1-Cas9 cells expressing FgH1t-UTG-sgSCR-GFP or FgH1tUTG-sgINTS6-A-GFP with stable knockout of INTS6 were treated with DMSO or AZ-5576 for 2 or 6 hours. 3 biological replicates were assessed for each treatment condition. Treated cells were lysed with CLB1 buffer (Zeptosens, Bayer) and quantified using a Pierce™ BCA Protein Assay Kit (ThermoFisher Scientific 23227). Samples were diluted (100%, 63%, 40%, 25%) with 9:1 CSBL1:CBL1 buffer (Zeptosens, Bayer) using a Sciclone Caplier ALH3000 liquid handling root (Perkin Elmer). Diluted samples were spotted onto ZeptoChips (Zeptosens; 2 technical replicates) using a Nano-plotter-NP2 non-contact microarray system (GeSim). Chips were blocked with BB1 buffer (Zeptosens, 1 hour) prior to incubation with primary antibodies (1:500 dilution for 2 hours, Key Resources Table) and Alexa-Fluor® conjugated secondary antibodies (1:1000 dilution for 4 hours, anti-rabbit Alexa-Fluor®-647, ThermoFisher Z25308; anti-mouse Alexa-Fluor®-647, ThermoFisher Z25008; anti-rat Alexa-Fluor®-647, ThermoFisher A21247). Alexa-Fluor® signal intensity was measured using a ZeptoREADER and relative fluorescence intensity (RFI) values were calculated using ZeptoVIEW software (Zeptosens, version 3.1). RFI values for each sample were normalized to secondary-antibody only controls.

### Phospho-peptide mass spectrometry

THP-1-Cas9 cells expressing FgH1t-UTG-sgSCR-GFP or FgH1tUTG-sgINTS6-A-GFP with stable knockout of INTS6 were treated with DMSO or AZ-5576 for 2 hours. 4 biological replicates were used for each condition. Cells were pelleted in Eppendorf^®^ protein LoBind tubes (Sigma Aldrich Z666505) washed twice with ice cold PBS and snap-frozen. Thawed cell pellets were incubated at 95°C for 10 minutes in SDS-lysis buffer (5% SDS, 10mM TCEP, 40mM 2-CAA, 10mM Tris-HCl pH 7.5) prior to DNA hydrolysis using 1% TFA and sample neutralisation through the addition of 3M Tris-HCl pH 7.5 to a final concentration of 160mM. Samples were incubated with PureCUBE carboxy agarose magnetic beads (Cube Biotech 50201) and 70% acetonitrile at room temperature for 20 minutes prior to 2 washes with 70% ethanol and 1 wash with 100% acetonitrile. Beads were lyophilised to remove residual acetonitrile and were then incubated with lysis buffer (50μL per sample; 10% trifluoroethanol, 100mM ammonium bicarbonate), Trypsin (Promega V5280) and Lysyl Endopeptidase (Wako 125-05061) at 1:25 ratio of enzyme:substrate. Beads were incubated for 2 minutes at room temperature in an ultrasonic water bath (Unisonics Australia) prior to incubation at 37°C for 4 hours with agitation (1200rpm). The lysis supernatant was isolated and the beads were washed with 1 volume of ultrapure H_2_O. The lysis supernatant and wash supernatant were pooled and TFA was added to a final concentration of 1%. Samples were centrifuged (20000g) and the supernatant was isolated. TFA-supplemented acetonitrile was added to the supernatant for a final concentration of 80% acetonitrile and 0.1% TFA prior to incubation at room temperature with gentle shaking for 30 minutes with PureCUBE Fe(III)-NTA beads (Cube Biotech 31501-Fe). Beads were washed 3 times with 80% acetonitrile / 0.1% TFA and transferred to C8 stage tips pre-wetted with 100% acetonitrile. Isolated phospho-peptides were eluted using 50% acetonitrile / 2.5% ammonium hydroxide pH 10 (2 × 20μL) and collected into tubes containing 30μL 10% TFA. Samples were lyophilized and peptides were resuspended in 50μL 5% formic acid prior to being transferred to C18 stage tips pre-wetted with 10% isopropyl alcohol, 60% acetonitrile and 5% formic acid. C18 stage tips were centrifuged (500g, 1 minute) prior to washing twice with 5% formic acid. Phospho-peptides were eluted in 60% acetonitrile and 5% formic acid (50μL) and were lyophilised to dryness prior to storage at minus 80°C. For analysis, phospho-peptides were resuspended in 2% acetonitrile and 1% formic acid and separated by reverse phase liquid chromatography using an ThermoFisher Scientific Easy-nLC 1200 UHPLC system (1.6μm C18 packed emitter tip column, 250mm × 75μm, Ion Opticks) with a linear 120 minute gradient at 400nL/min flow rate from 98% solvent A (0.1% formic acid / ultrapure H_2_O) to 37% solvent B (0.1% formic acid / acetonitrile). The UHPLC was coupled online to a Q-Exactive HF Orbitrap mass spectrometer equipped with a nano-electrospray ionization source (ThermoFisher Scientific) and a column oven at 50°C (Sonation, Germany). The Q-Exactive HF was operated in a data-dependent mode, switching automatically between one full-scan and subsequent MS/MS scans of the ten most abundant peaks. The instrument was controlled using Exactive series version 2.9 and Xcalibur 4.1.31.9. Full-scans (m/z 350–1,850) were acquired with a resolution of 60,000 at 200 m/z. The 10 most intense ions were sequentially isolated with a target value of 10,000 ions and an isolation width of 1.4 m/z and fragmented using HCD with NCE of 27. Maximum ion accumulation times were set to 100ms for full MS scan and 110ms for MS/MS. Raw files were analyzed using MaxQuant (version 1.5.8.3). The database search was performed using the Uniprot *Homo sapiens* database plus common contaminants with strict trypsin specificity allowing up to 3 missed cleavages. The minimum peptide length was 7 amino acids. Carbamidomethylation of cysteine was a fixed modification while N-acetylation of proteins N-termini, oxidation of methionine and phosphorylation of Serine/Threonine/Tyrosine were set as variable modifications. During the MaxQuant main search, precursor ion mass error tolerance was set to 4.5 ppm and fragment ions were allowed a mass deviation of 20 ppm. PSM and protein identifications were filtered using a target-decoy approach at a false discovery rate (FDR) of 1% with the match between runs option enabled.

Further analysis was performed using a custom pipeline developed in R, which utilizes the MaxQuant output file evidence.txt. A feature was defined as the combination of peptide sequence, charge and modification. Features not found in at least 80% of the replicates in one group were removed. To correct for injection volume variability, feature intensities were normalized by converting to base 2 logarithms and then multiplying each value by the ratio of maximum median intensity of all replicates over median replicate intensity. Missing values were imputed using a random normal distribution of values with the mean set at mean of the real distribution of values minus 2.2 s.d., and a s.d. of 0.15 times the s.d. of the distribution of the measured intensities. The probability of differential peptide expression between groups was calculated using the Limma R package (version 3.34.9). Probability values were corrected for multiple testing using Benjamini–Hochberg method. The R package pheatmap (v1.0.12) was used to compute heatmaps of phospho-sites with the indicated significance cutoff. The values in the heatmaps represent the median of z-norm values of biological replicates for each condition. Filtering conditions were applied to phospho-sites in addition to significance cutoffs. To be plotted, z-norm values of phospho-sites were to meet the following criteria: SCR_CDK9i – SCR_UT < −0.3 & SCR_UT – INTS6_KO_UT < 0.3 & SCR_CDK9i *≤* INTS6_KO_UT & SCR_CDK9i *≤* INTS6_KO_CDK9i. Kmean cluster line plots and phospho-peptides scatter plots were generated by using the R package ggplot2 (v3.3.0). To be plotted, z-norm values of phospho-sites in kmean cluster and in single site plots were to meet the following criteria: SCR_CDK9i < SCR_UT & SCR_CDK9i *≤* INTS6_KO_UT & SCR_CDK9i *≤* INTS6_KO_CDK9i. In the phospho-peptides scatterplots, each bar represents the median of z-norm values of different biological replicates for the same condition. Gene ontology (GO) analysis was performed using Metascape (release 3.5) (Zhou et al., 2019).

### Recombinant protein kinase / phosphatase assay

Reactions were performed in kinase reaction buffer (25mM Tris-HCl pH 7.4, 2mM dithiothreitol (DTT), 10mM MgCl-_2_) supplemented with 200mM Mg^+^-ATP (Sigma Aldrich A9187) in 40μL final reaction volumes. 0.5μg human RNA polymerase II CTD repeat YSPTSPS protein (Abcam ab81888) was incubated with/without 0.4μg human 6X-His-CDK9/Cyclin T1 (Merck, 140685) and/or 0.4μg human GST-tagged PP2A-α (Sigma, SRP5336) in the presence or absence of AZ’5576 or okadaic acid (Abcam ab120375) for 45 minutes at 30°C (300rpm). Reactions were stopped through the addition of 20μL 2X laemmli lysis buffer (120mM Tris-HCl pH 6.8, 20% (v/v) glycerol, 4% (w/v) SDS) and incubation at 95°C for 10 minutes. Reactions were analyzed by SDS-PAGE and immunoblotting as described previously.

### 3’RNA Quant-Seq

THP-1-Cas9 cells expressing FgH1t-UTG-sgSCR-GFP or FgH1tUTG-sgINTS6-A-GFP with stable knockout of INTS6 were treated with PBS or LPS (1µg/mL) for 1, 2, 4 and 6 hours (2 × 10^6^ cells per condition in duplicate). Cells were lysed with 600µL TRIzol® (ThermoFisher Scientific 15596026) and total RNA was isolated using the Diret-zol RNA mini-prep kit (Zymo Research R2051). Sequencing libraries were prepared using the QuantSeq 3’-mRNA Seq Library Prep Kit for Illumina (Lexogen) from 500ng total RNA and 75 base-pair single-end reads were sequenced on the Illumina NextSeq 500. Sequencing files were demultiplexed (Bcl2fastq, v2.17.1.14) and QC was performed on FASTQ files using FASTQC (v0.11.6). Sequencing reads were trimmed (cutadapt v2.1) and aligned to the Hg19 human reference genome using HISAT2 (v2.1.0). Read counting across genomic features was performed using FeatureCounts (Subread, v2.0.0) and differential gene expression analysis was performed using Voom-Limma in R (v3.42.2). Gene set enrichment analyses were performed using the Broad Institute GSEA software (Subramanian et al., 2005).

HeLa cells infected with shLUC or shINTS6 were treated with PBS or EGF (0.1µg/mL) for 15, 55, and 180 minutes. Cells were lysed in Tri-reagent and total RNA was isolated using the Direct-zol RNA mini-prep kit (Zymo). For spike-in, 200ng of RNA from *D.melanogaster* S2 cells was added to 1µg of total RNA for each timepoint. Sequencing libraries were prepared at the Wistar Genomics Facility using the QuantSeq 3’-mRNA Seq Library Prep Kit for Illumina (Lexogen) and 75 base-pair single-end reads were sequenced on the Illumina NextSeq 500. Reads were aligned to hg19 and dm5 reference genomes using STAR v2.5 (Dobin et al., 2013) in 2-pass mode with the following parameters: –quantMode TranscriptomeSAM –outFilterMultimapNmax 10 –outFilterMismatchNmax 10 –outFilterMismatchNoverLmax 0.3 –alignIntronMin 21 –alignIntronMax 0 –alignMatesGapMax 0 –alignSJoverhangMin 5 –runThreadN 12 –twopassMode Basic –twopass1readsN 60000000 –sjdbOverhang 100. BAM files were filtered based on alignment quality (q = 10) using Samtools v0.1.19 (Li et al., 2009). FeatureCounts (Liao et al., 2014) was used to count reads mapping to each gene, and differential gene expression analysis was performed using Voom-Limma in RStudio (v3.42.2) with normalizationFactors calculated from the *D.melanogaster* spike-in. Data was visualized using ggplot2 (v3.3.1).

### Chromatin Immunoprecipitation Sequencing (ChIP-Seq)

For THP-1 parental, THP-1-Cas9 FgH1t-UTG-sgSCR-GFP or FgH1tUTG-sgINTS6-A-GFP, 20 – 50 × 10^6^ THP-1 cells were incubated with DMSO, AZ-5576 or DBK-1154 for indicated time-points. Cells were washed once with cold PBS prior to cross-linking at room temperature for 10 minutes with 1/10^th^ volume fresh formaldehyde solution (11% formaldehyde, 0.05mM EGTA, 1mM EDTA, 100mM NaCl, 50mM Hepes-KOH pH7.5). To quench the cross-linking reaction, 1/20^th^ volume of 2.5M glycine was added and cells incubated at room temperature for a further 5 minutes. Cells were washed with ice-cold PBS and nuclei were isolated by 3 successive 10 minute incubations ice with cold nuclear extraction buffer (20mM Tris-HCl pH 8, 10mM NaCl, 0.5% IGEPAL^®^ CA-630, 2mM EDTA). Cell nuclei were resuspended in sonication buffer (20mM Tris-HCl pH 7.5, 150mM NaCl, 2mM EDTA, 0.3% SDS, 1% IGEPAL^®^ CA-630) and sonicated for 18 minutes using the Covaris S2 instrument (20% Duty Cycle, 1000 cycles/burst, 10 Intensity). Fragmented chromatin was diluted 1:1 with dilution buffer (20mM Tris-HCl pH 8, 150mM NaCl, 2mM EDTA, 1% TritonX-100). For each immunoprecipitation, diluted chromatin was incubated overnight at 4°C with Protein A and Protein G Dynabeads (ThermoFisher Scientific 10002D, 10004D; 25µL each) and indicated antibodies (Key Resources Table). Post-incubation, Dynabeads were washed once each with dilution buffer, ChIP wash buffer 1 (20mM Tris-HCl pH 8, 500mM NaCl, 2mM EDTA, 0.1% SDS, 1% TritonX-100) and ChIP wash buffer 2 (20mM Tris-HCl pH 8, 250mM LiCl, 2mM EDTA, 0.5% deoxycholate, 0.5% IGEPAL^®^ CA-630) and washed twice with TE buffer (10mM Tris-HCl pH 7.5, 1mM EDTA). Washed beads were incubated with shaking at 55°C for 1 hours in reverse crosslinking buffer (200mM NaCl, 100mM NaHCO_3_, 1% SDS, 300µg/mL Proteinase-K) followed by incubation of the supernatant overnight at 65°C. DNA was isolated using the Zymogen ChIP DNA Clean and Concentrator Kit (Zymo Research D5205). Sequencing libraries were prepared using the NEBNext Ultra II DNA Library Prep Kit (NEB E7645) and 200 – 500 base-pair size selection was performed using a Pippin Prep system (Sage Science). 75 base-pair single end reads or 40 base-pair paired end reads were sequenced on the Illumina NextSeq500. Sequencing files were demultiplexed (Bcl2fastq, v2.17.1.14) and QC was performed on FASTQ files using FASTQC (v0.11.6). Sequenced reads were aligned to the Hg19 human or dm3 *D. melanogaster* reference genomes using Bowtie2 (v2.3.4.1) and generated SAM files were converted to BAM files using Samtools (v.1.9) using the view function. Samtools was used to further sort, index and remove potential PCR duplicates (rmdup) from BAM files which were converted to BigWig files using Deeptools (v3.0.0; bamCoverage, --nomalizeUsing CPM --smoothLength 150 --binSize 50 -e 200, scaleFactor 1). BigWig files were visualized using the Integrative Genomics Viewer (IGV; Broad Institute). Average read density across defined genomic intervals was computed using the Deeptools computeMatrix function and the resulting matrices were used to generate chromatin occupancy heatmaps using the plotHeatmap function. Pausing indices for Hg19 and dm3 were calculated as a ratio TSS read density (+/- 250bp) over gene body density (+500 base pairs from TSS and −250 base pairs from TES). In more detail, subread (v2.0.0; featureCounts -O -M -T 16 -F SAF) was used to count reads within TSS and gene body intervals generated using bedtools (v2.27.1) slop function that included Hg19 or dm3 chromosome sizes (downloaded from UCSC). Read density was calculated as RPKM in Rstudio (v3.6.1) using limma (v3.40.6) and edgeR (v3.26.8) packages and visualized using ggplot2 (v3.3.1).

For wild-type HeLa, cells were serum-starved (with 0.3% SCS) for 72 hours prior to induction of gene expression with 0.1µg/mL recombinant EGF (Gibco) or PBS for 15 minutes. For knockdowns, HeLa cells infected with indicated shRNAs were placed under serum-starvation (0.3% SCS) the night before harvesting. Cells were treated for 2 hours with 300nM AZ5576, 1 hour with 2µM Flavopiridol, or time-matched DMSO control, and subsequently treated with PBS or 0.1µg/mL EGF for 15 minutes. ChIP-seq was performed as previously described (Lai et al., 2015), with some modifications. To allow for accurate normalization between treatment conditions, tandem ChIP-seq was performed with a *D.melanogaster-* specific antibody similar to that described by (Egan et al., 2016). For each replicate, 10×10^6^ HeLa cells were harvested and cross-linked for 5 minutes at room temperature with 1% formaldehyde for 5 minutes at room temperature and washed twice with PBS. The chromatin pellet was resuspended in ChIP lysis buffer (150mM NaCl, 1% Triton X-100, 0.7% SDS, 500mM DTT, 10mM Tris-HCl, 5mM EDTA), and supplemented with chromatin from 2×10^6^ formaldehyde-crosslinked *D.melanogaster* S2 cells. The chromatin was sheared to an average length of 200–400 base pairs using a Covaris S220 Ultrasonicator. The chromatin lysate was diluted with SDS-free ChIP lysis buffer. For ChIP-seq, the following were added to the sheared chromatin and incubated 4°C overnight: 10 µg of human antibody, 2 µg Drosophila spike-in antibody, and Protein A or Protein G Dynabeads (Invitrogen). After incubation, beads were washed twice with each of the following buffers: Mixed Micelle Buffer (150 mM NaCl, 1% Triton X-100, 0.2% SDS, 20 mM Tris-HCl, 5 mM EDTA, 65% sucrose), Buffer 500 (500 mM NaCl, 1% Triton X-100, 0.1% Na-deoxycholate, 25 mM HEPES, 10 mM Tris-HCl, 1 mM EDTA), LiCl/detergent wash (250 mM LiCl, 0.5% Na-deoxycholate, 0.5% NP-40, 10 mM Tris-HCl, 1 mM EDTA), followed by a final wash with 1X TE. Finally, beads were resuspended in 1X TE with 1% SDS and incubated at 65°C for 10 min to elute. Elution was repeated twice, and the samples were incubated overnight at 65°C to reverse cross-link, alongside the untreated input (5% of the starting material). After treatment with 0.5 mg/ml proteinase K for 3 hours, DNA was purified with Zymo ChIP DNA Clean Concentrator kit (Zymo research) and quantified by QUBIT. Barcoded libraries were made with NEB Ultra II DNA Library Prep Kit for Illumina, and sequenced on Illumina NextSeq 500, producing 75 base pair single-end reads or 40 base pair paired-end reads. Sequences were aligned to the human reference hg19 and *D.melanogaster* dm5 genome using Burrows Wheeler Alignment tool (BWA), with the MEM algorithm (Li, 2013). Using Samtools, aligned reads were filtered based on mapping quality (MAPQ>10) and PCR duplicates removed (rmdup). Data was visualized on the UCSC Genome Browser (http://genome.ucsc.edu/) with Bigwig files generated with Deeptools (v2.4.2): bamCoverage --binSize 10 --normalizeTo1x 3137161264 --ignoreForNormalization chrX. Peaks were called using MACS2 (Zhang et al., 2008) at 5% FDR, with default parameters (Zhang et al., 2008). For *D.melanogaster* spike-in normalization, the number of hg19 filtered reads was divided by the lowest number of dm5 filtered reads for each set of experiments, resulting in a downsampling factor for each sample. Normalized BAM files for each sample were generated using Samtools view –s with the downsampling factor, and normalized Bigwigs were generated from the BAM files using Deeptools bamCoverage --binSize 10. Average read density across defined genomic intervals was computed using seqMINER 1.3.3 package (Ye et al., 2011) with the following parameters: left and right extensions = 5.0 kb; internal bins = 160; flanking region bins = 20. Read density matrices were exported to generate average profiles in R with ggplot2 (v3.3.1). Genomic intervals were generated using valr (v0.6.1) (Riemondy et al., 2017). Pausing index was calculated as a ratio of TSS read density (−150bp/+50bp) over gene body density (+50 bp from TSS through the TES).

### 4sU Nascent RNA sequencing

20 × 10^6^ THP-1-Cas9 cells expressing FgH1t-UTG-sgSCR-GFP or FgH1tUTG-sgINTS6-A-GFP with stable knockout of INTS6 were treated with DMSO (6 hours) or AZ-5576 for 2 or 6 hours. Cells were labelled with 500μM 4-thiouridine (4sU; Sigma Aldrich T4509) for the final 30 minutes of the treatment time and were lysed in TRIzol. *D.melanogaster* S2 cells (20 × 10^6^) were also labelled with 4sU as above. Lysates were mixed with 1/5^th^ volume of chloroform and the aqueous phase was collected after centrifugation at 4°C (13000g, 15 minutes). Total RNA was precipitated in isopropanol (room temperature for 10 minutes, 4°C for 10 minutes at 13000g) and washed once with 70 percent ethanol (4°C for 15 minutes at 13000g). RNA was resuspended in ultrapure nuclease-free H_2_O and denatured at 65°C for 10 minutes. 80μg human RNA (with 5% S2 RNA spike-in) was incubated for 1.5 hours with constant rotation at room temperature with 10mM Tris pH7.4, 1mM EDTA, 200μg/mL dimethylformamide (Sigma Aldrich, 227056) and 300μg EZ-Link™ HPDP Biotin (ThermoFisher Scientific 21341). The ‘biotin-labeling reaction’ was supplemented with 1 volume chloroform and vortexed well prior to centrifugation at 4°C (20000g, 5 minutes). The aqueous phase was isolated and chloroform isolation was repeated twice, prior to the addition of 5M NaCl to a final concentration of 0.5M and 1 volume of isopropanol. Precipitated RNA was centrifuged at 4°C (20000g, 20 minutes), washed with 75% ethanol (4°C, 20000g, 20 minutes) and resuspended in nuclease-free H_2_O (1μg/1μg RNA input) prior to incubation at 65°C for 10 minutes. Biotinylated RNA was incubated with constant rotation for 15 minutes at room temperature with μMACs streptavidin beads and was isolated using μMACs columns (μMACs Strepavidin Kit, Miltenyi Biotec 130-074-101) that had been previously equilibrated with wash buffer (100mM Tris-HCl pH 7.4, 10mM EDTA, 1M NaCl, 0.1% Tween-20). Columns were washed with wash buffer pre-warmed to 65°C (5 × 0.9mL washes) and room temperature wash buffer (5 × 0.9mL washes) and RNA was eluted using DTT (100mM, 2 × 100μL) and collected into 700μL RLT buffer from the RNeasy MinElute Cleanup Kit (Qiagen 74204). RNA was isolated using the RNeasy MinElute kit as per manufacturer’s instructions and was quantified using the Agilent Tapestation High Sensitivity RNA kit (Agilent 067-5576). RNA sequencing libraries were prepared using the NEBNext Ultra II Directional RNA Library Prep Kit for Illumina and 75 base-pair single-end reads were sequenced using the Illumina NEXTseq 500. Sequencing files were demultiplexed (Bcl2fastq, v2.17.1.14) and QC was performed on FASTQ files using FASTQC (v0.11.6). Sequenced reads were aligned to a combined Hg19/dm3 reference genome using Bowtie (v2.3.4.1) and BAM files were generated and processed as described for ChIP-seq. Normalization to the *D.melanogaster* S2 RNA spike-in was performed similarly to that described previously (Orlando et al., 2014), with reads mapping to Hg19 human and dm3 *D.melanogaster* genomes calculated using FeatureCounts (Subread v2.0.0) and a scaleFactor determined by calculating the proportion of reads mapping to the dm3 genome relative to the combined Hg19/dm3 genome. BigWig files were generated using Deeptools (v3.0.0) bamCoverage function using the appropriate scale Factor (--normalizeUsing CPM --smoothLength 150 --binSize 50 –e 200 scaleFactor #). Scaled BigWig files were visualized in IGV and the bigwigCompare function (Deeptools). Data was visualized in RStudio using ggplot2 (v3.3.1), ggrepel (v0.8.2) and ggfortify (v0.4.10) packages.

### TT-seq

HeLa cells infected with indicated shRNAs were placed under serum-starvation (0.3% SCS) the night before harvesting. Cells were treated with EGF (0.1μg/mL) and concurrently labelled with 500μM 4-thiouridine (4sU; Sigma Aldrich T4509) for 15 minutes. Cells were lysed in QIAzol lysis reagent (QIAgen, #79306), mixed with 1/5^th^ volume of chloroform, and the aqueous phase collected after centrifugation at 4°C (13000g, 15 minutes). Total RNA was precipitated in isopropanol (room temperature for 10 minutes), centrifuged at 4°C for 10 minutes at 13000g, and washed twice with 75 percent ethanol. Pellet was air-dried and then re-suspended in ultrapure nuclease-free H_2_O on ice for 10 minutes, then 60°C for 5 minutes. 300μg human RNA (with 5% S2 4sU-labeled RNA spike-in) was fragmented with the Bioruptor Plus for one cycle of 30s/30s ON/OFF at high settings. Fragmentation was checked by TapeStation analysis. 150μg fragmented RNA was incubated at 65°C for 10 minutes, then on ice for 5 minutes before adding biotin-labeling reagents (100mM Tris pH 7.5, 10mM EDTA, 200μL dimethylformamide (DMF), 200μg EZ-link™ HPDP Biotin (ThermoFisher Scientific 21341)) and incubating in the dark at 24°C and 800rpm for 2 hours. Chloroform was added to the reaction and mixed well. After centrifugation, the upper aqueous phase was collected and the RNA was precipitated by the addition of 1/10^th^ volume of 5M NaCl and 1 volume of isopropanol, centrifuged at 4°C (20000g, 30 minutes), and washed with ice cold 75% ethanol. RNA was resuspended in ultrapure nuclease-free H_2_O, incubated on ice for 10 minutes, then 65°C for 5 minutes. Biotin-labeled RNA was enriched using Streptavidin beads (Invitrogen). Beads were equilibrated with wash buffer (100mM Tris pH 7.5, 10mM EDTA, 1M NaCl, 0.1% (v/v) Tween-20) before incubation with biotinylated RNA at 4°C and 800rpm for 15 minutes. Beads were washed 3x with 65°C wash buffer and 3x with room temperature wash buffer. RNA was eluted twice by resuspending the beads in 100mM DTT buffer, and incubating for 5 minutes. RNA was purified using the RNA Clean and Purification kit (Zymo research) with DNAse step following manufacturer’s instructions. Sequencing libraries were prepared with the NEBNext Ultra II Directional RNA Library Prep Kit for Illumina and 75bp single-end or 40bp paired-end reads were sequenced using the Illumina NEXTseq 500. Reads were aligned to hg19 and dm5 reference genomes using STAR v2.5 (Dobin et al., 2013) in 2-pass mode with the same parameters described above for HeLa 3’-Quantseq. BAM files were filtered based on alignment quality (q = 10) using Samtools v0.1.19 (Li et al., 2009). The latest annotations were obtained from Ensembl to build reference indexes for the STAR alignments. FeatureCounts (Liao et al., 2014) was used to count reads mapping to each gene. RSEM (Li and Dewey, 2011) was used to obtain TPM and FPKM (Fragments Per Kilobase of exon per Million fragments mapped). Normalization to the *D.melanogaster* S2 RNA spike-in, and subsequent normalized BAM and BigWig file generation was performed as described above for HeLa ChIP-seq. Average read density across defined genomic intervals was computed on the normalized BAM files using seqMINER 1.3.3 package (Ye et al., 2011) with the following parameters: left and right extensions = 5.0 kb; internal bins = 160; flanking region bins = 20. Read density matrices, featureCounts, and RSEM outputs were exported for plotting in RStudio with ggplot2 (v3.3.1).

### FastGRO

FastGRO was performed as described (Barbieri et al., 2020). HeLa cells infected with shLUC or shINTS6 were treated with DMSO or 300nM CDK9i AZ5576 for 2 hours, then subsequently treated for 15 minutes with 0.1μg/mL EGF. Per condition, 20×10^6^ cells were harvested and washed twice with ice-cold PBS before adding swelling buffer (10 mM Tris-HCL pH 7.5, 2mM MgCl2, 3 mM CaCl2, 2U/ml Superase-in (Invitrogen)). Cells were swollen for 5 min on ice, washed with swelling buffer with 10% glycerol, and then lysed in lysis buffer (10 mM Tris-HCL pH 7.5, 2mM MgCl2, 3mM CaCl2, 10% glycerol, 1mL Igepal (NP-40), 2 U/ml Superase-in) to isolate nuclei. Nuclei were washed twice with lysis buffer and resuspended in freezing buffer (40% glycerol, 5mM MgCl2, 0.1mM 0.5M EDTA, 50mM Tris-HCL pH 8.3) to a concentration of 2×10^7^ nuclei per 100 μL. Nuclei were frozen on dry ice and stored at −80 °C. For the run-on reaction, nuclei were thawed on ice, an equal volume of pre-warmed nuclear run-on reaction buffer (10mM Tris-HCl pH 8, 5mM MgCl2, 300mM KCl, 1mM DTT, 500μM ATP, 500μM GTP, 500μM 4-thio-UTP, 2μM CTP, 200 U/ml Superase-in, 1% Sarkosyl (N-Laurylsarcosine sodium salt solution) was added. This was incubated for 7 min at 30°C for the nuclear run-on. Nuclear run-on RNA was extracted with TRIzol LS reagent (Invitrogen) following the manufacturer’s instructions and precipitated with ethanol. RNA was then resuspended in water and the concentration was determined with Qubit High Sensitivity Assay kit (Invitrogen). Spike-in *D.melanogaster* RNA was added at 10%, and RNA was fragmented with a Bioruptor UCD-200 for 1-5 cycles of 30s ON/30s OFF, high settings. Fragmentation efficiency was analyzed by running fragmented and unfragmented RNA on Agilent 2200 TapeStation using High Sensitivity RNA ScreenTapes following manufacturer’s instructions. Fragmented RNA was incubated in Biotinylation Solution (100 mM Tris pH 7.5, 10 mM EDTA pH 8.0, 40% dimethylformamide, 200 g/mL EZ-link HPDP Biotin (Thermo Scientific)) for 2h in the dark at 25 °C and 800 rpm. After ethanol precipitation, the biotinylated RNA was resuspended in water and separated with M280 Streptavidin Dynabeads (Invitrogen). 100 μl/sample of beads were washed twice with 2 volumes of freshly prepared wash buffer (100 mM Tris pH 7.5, 10mM EDTA pH 8.0, 1M NaCl, 0.1% (v/v) Tween-20) and resuspended in 1 volume of wash buffer and added to the biotinylated-RNA. After 15 minutes in rotation at 4°C, beads were washed three times with wash buffer pre-warmed at 65°C and three times with room temperature wash buffer. 4-S-UTP containing RNA was eluted in 100mM DTT buffer and purified with RNA Clean and Purification kit (Zymo Research) with in-column DNAse reaction to eliminate traces of genomic DNA. The eluted RNA was quantified with Qubit High Sensitivity Assay kit (Invitrogen) and used to produce barcoded RNA sequencing libraries using the NEBNext Ultra II Directional RNA Library Prep kit (New England Biolabs). Libraries were sequenced on Illumina NextSeq 500. Reads were aligned to hg19 and dm5 reference genomes using STAR v2.5 (Dobin et al., 2013) in 2-pass mode with the same parameters described above for HeLa 3’-Quantseq. BAM files were filtered based on alignment quality (q = 10) using Samtools v0.1.19 (Li et al., 2009). The latest annotations were obtained from Ensembl to build reference indexes for the STAR alignments. FeatureCounts (Liao et al., 2014) was used to count reads mapping to each gene. RSEM (Li and Dewey, 2011) was used to obtain TPM and FPKM (Fragments Per Kilobase of exon per Million fragments mapped). Normalization to the *D.melanogaster* S2 RNA spike-in, and subsequent normalized BAM and BigWig file generation, was performed as described above for HeLa ChIP-seq. Average read density across defined genomic intervals was computed on the normalized BAM files using seqMINER 1.3.3 package (Ye et al., 2011) with the following parameters: left and right extensions = 5.0 kb; internal bins = 160; flanking region bins = 20. Read density matrices, featureCounts, and RSEM outputs were exported for plotting in RStudio with ggplot2 (v3.3.1).

### *In vivo* Methods

All *in vivo* experiments performed in this study were approved by the Peter MacCallum Cancer Centre (PMCC) Animal Ethics Committee under ethics approval numbers E555, and all NOD-*scid* IL2Rg^null^ (NSG) mice utilised were bred in-house (PMCC). For disseminated MV4-11 therapy experiments, 4-6-week-old female NSG mice were IV-injected with 5 × 10^6^ MV4;11 cells expressing the MSCV-Luc2-mCherry vector (Bots et al., 2014). For long-term survival experiments, treatment commenced when peripheral blood was detected to be 1% mCherry positive as determined by flow cytometry (day 18). Mice were administered 15 mg/kg AZD’4573 twice daily via intraperitoneal injection (IP; 100 mL) two hours apart, with 40 mg/kg DBK-1154 (100 mL) administered via oral gavage two hours prior to, and two hours post, the AZD’4573 injections. This regimen was implemented in a 2 days on, 5 days off manner and repeated for four consecutive cycles of therapy. For Cross-sectional analysis, therapy commenced 30 days post inoculation, and mice were treated as per the above regimen. Mice were culled 24 hours and 96 hours post the conclusion of 1 cycle of therapy. At endpoint in all experiments, blast cells were harvested from femur and tibial/fibial bone and spleen, and assessed for mCherry percentage via flow cytometry using a BD FACSymphony™.

## KEY RESOURCES TABLE

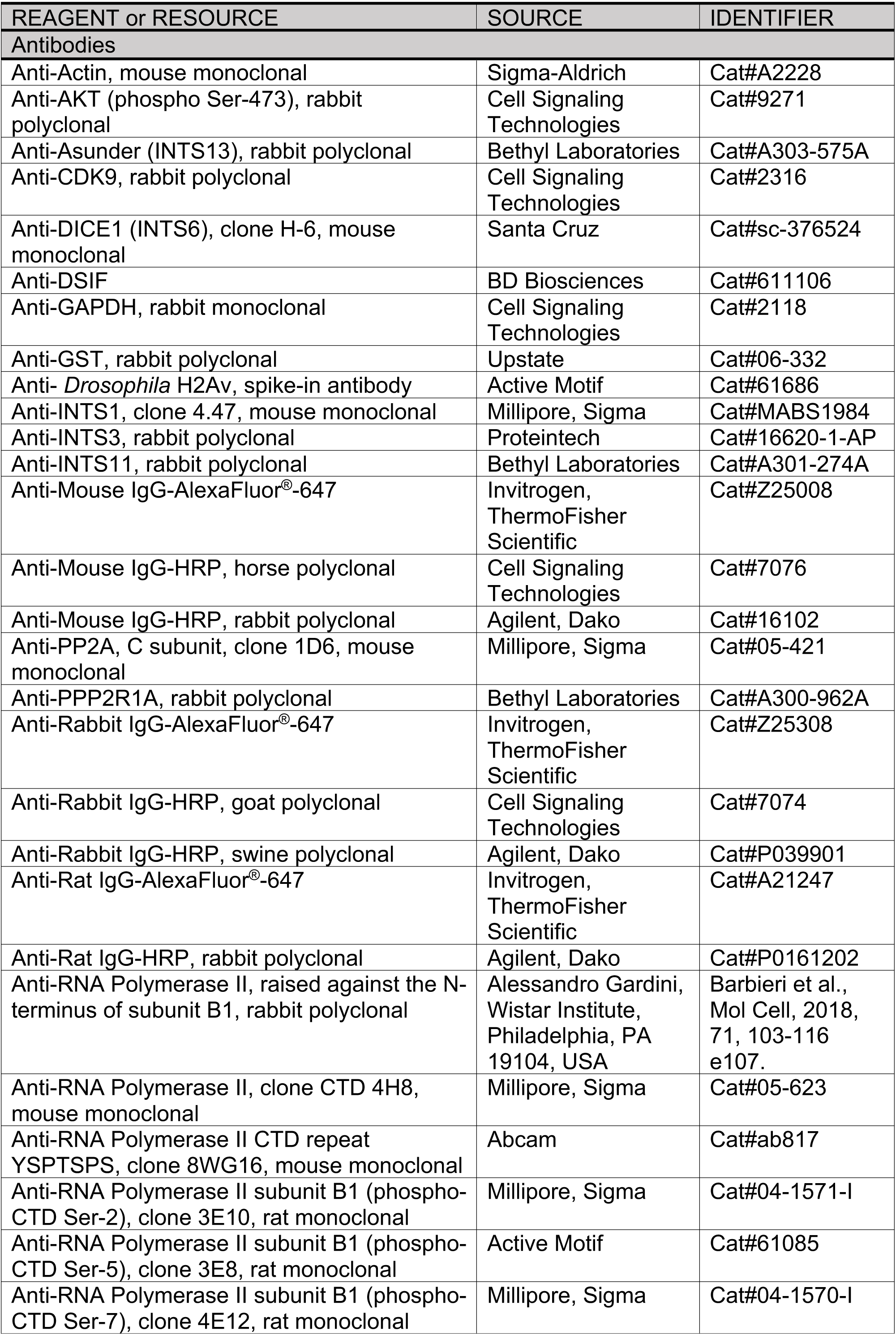
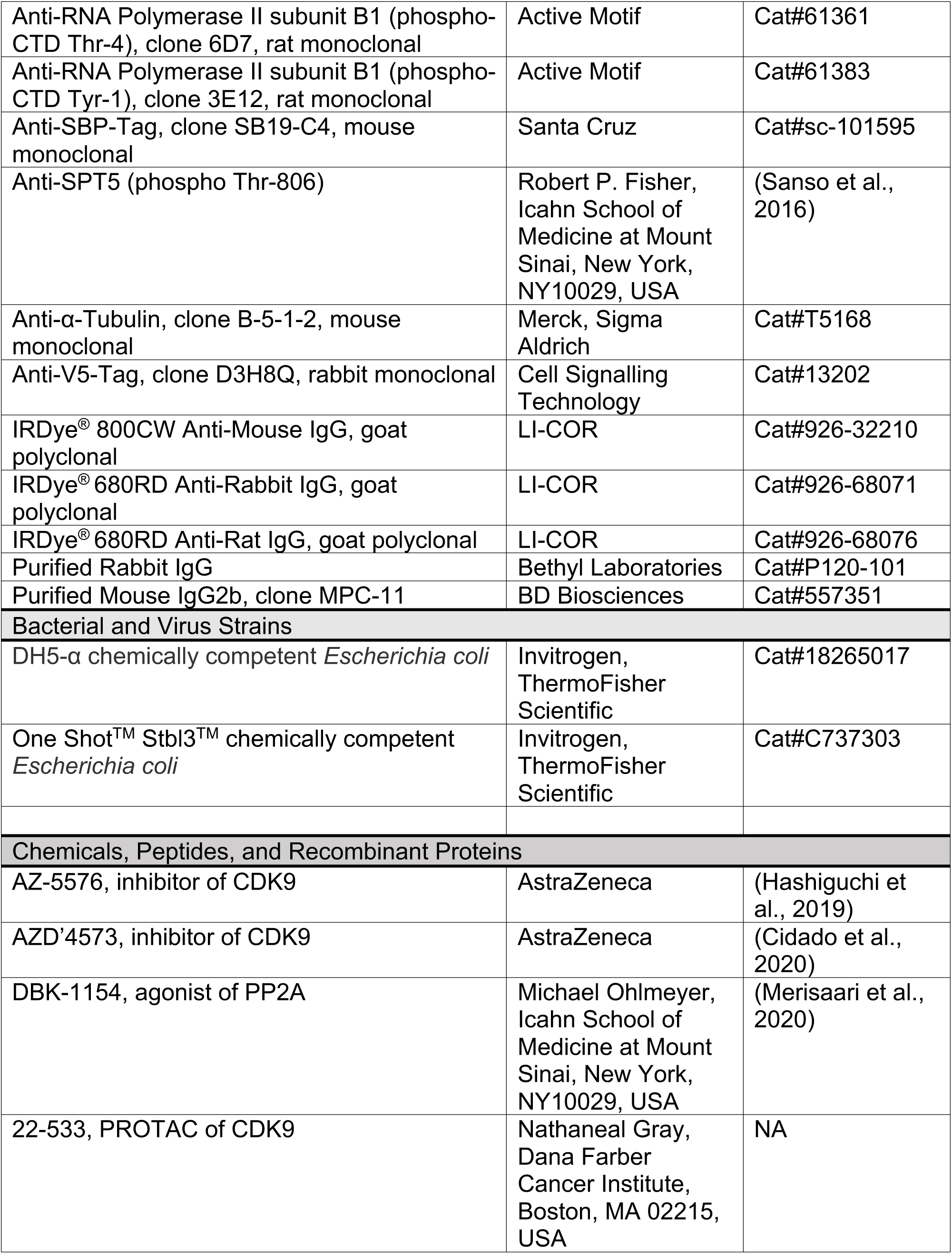
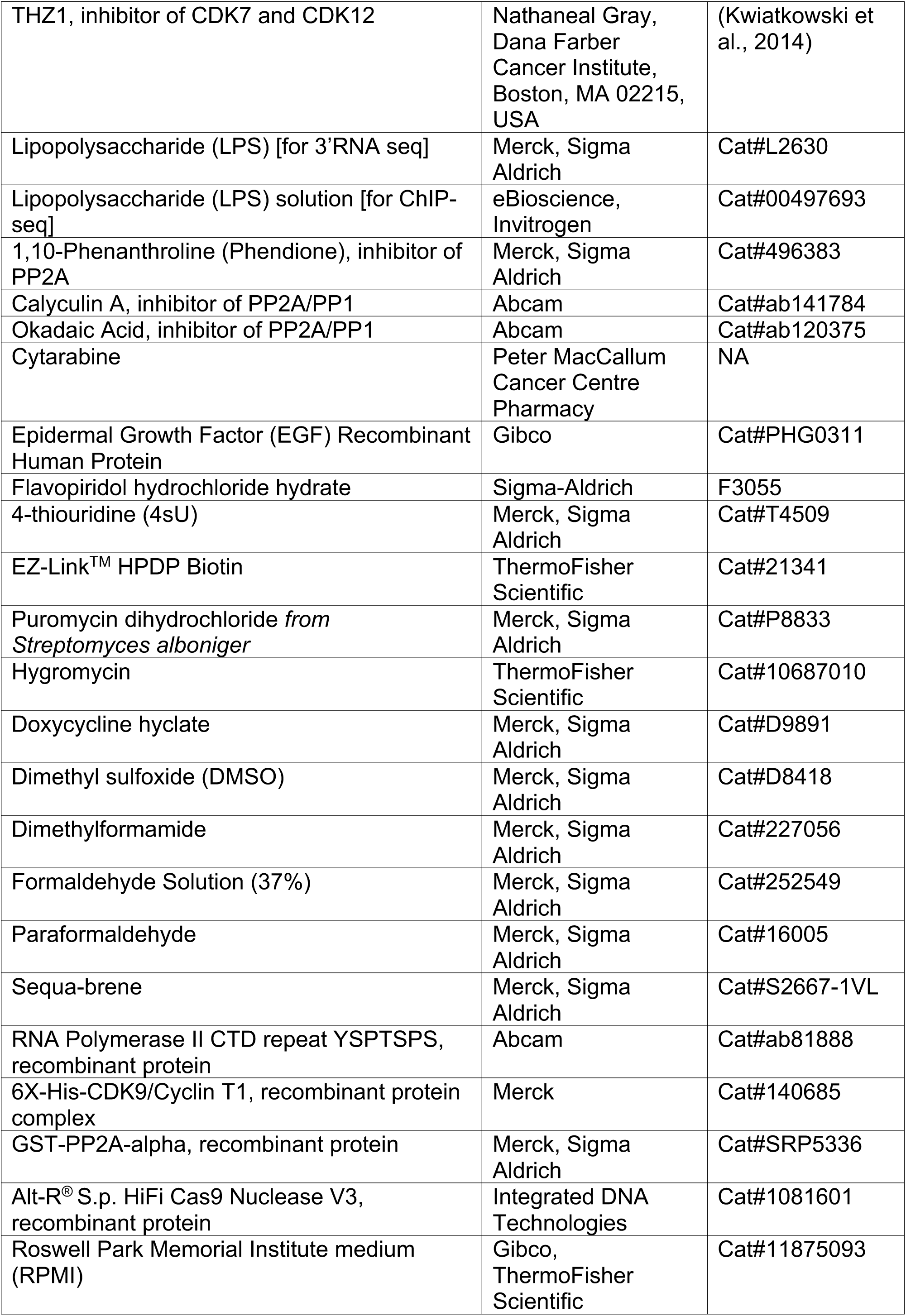
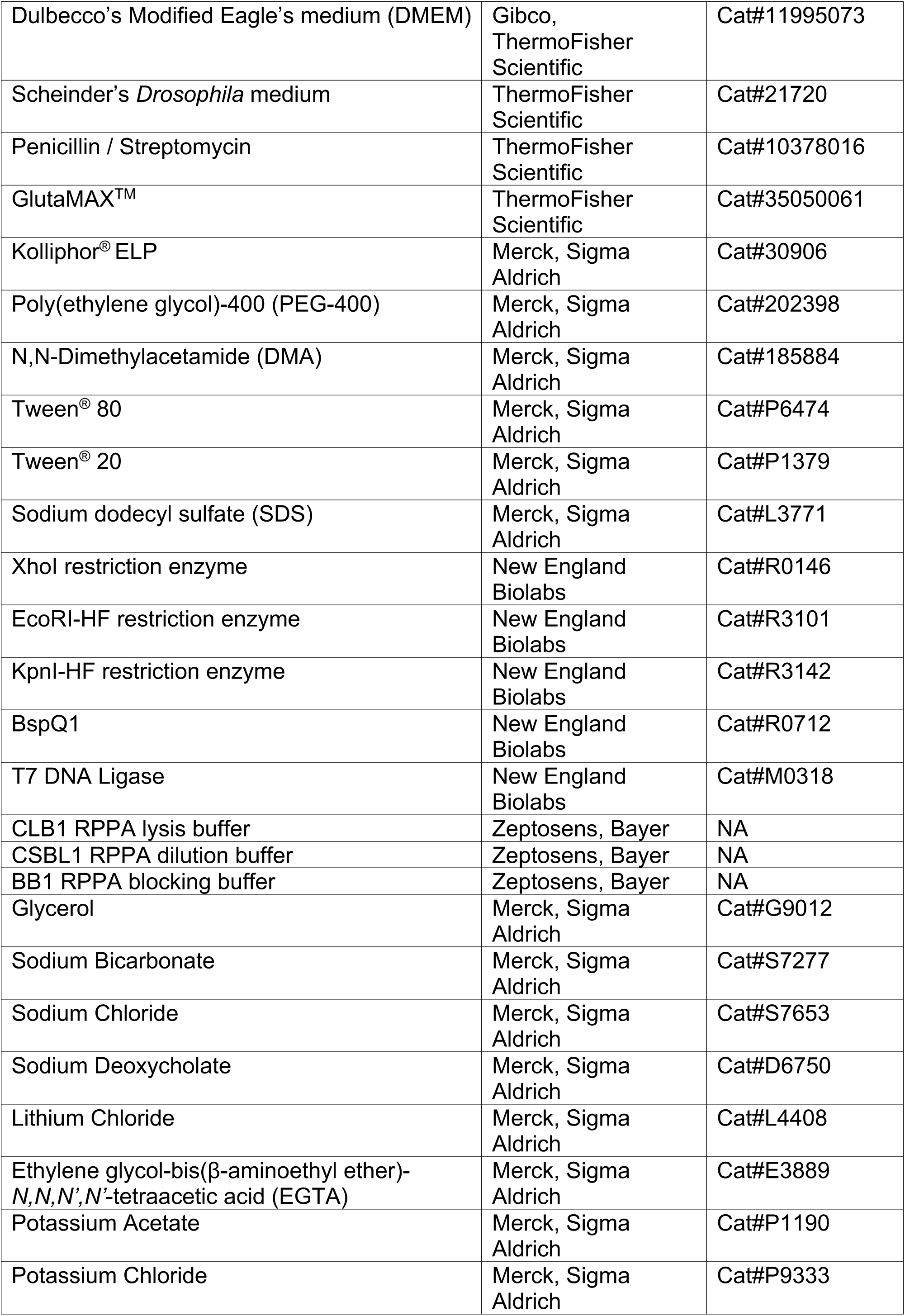
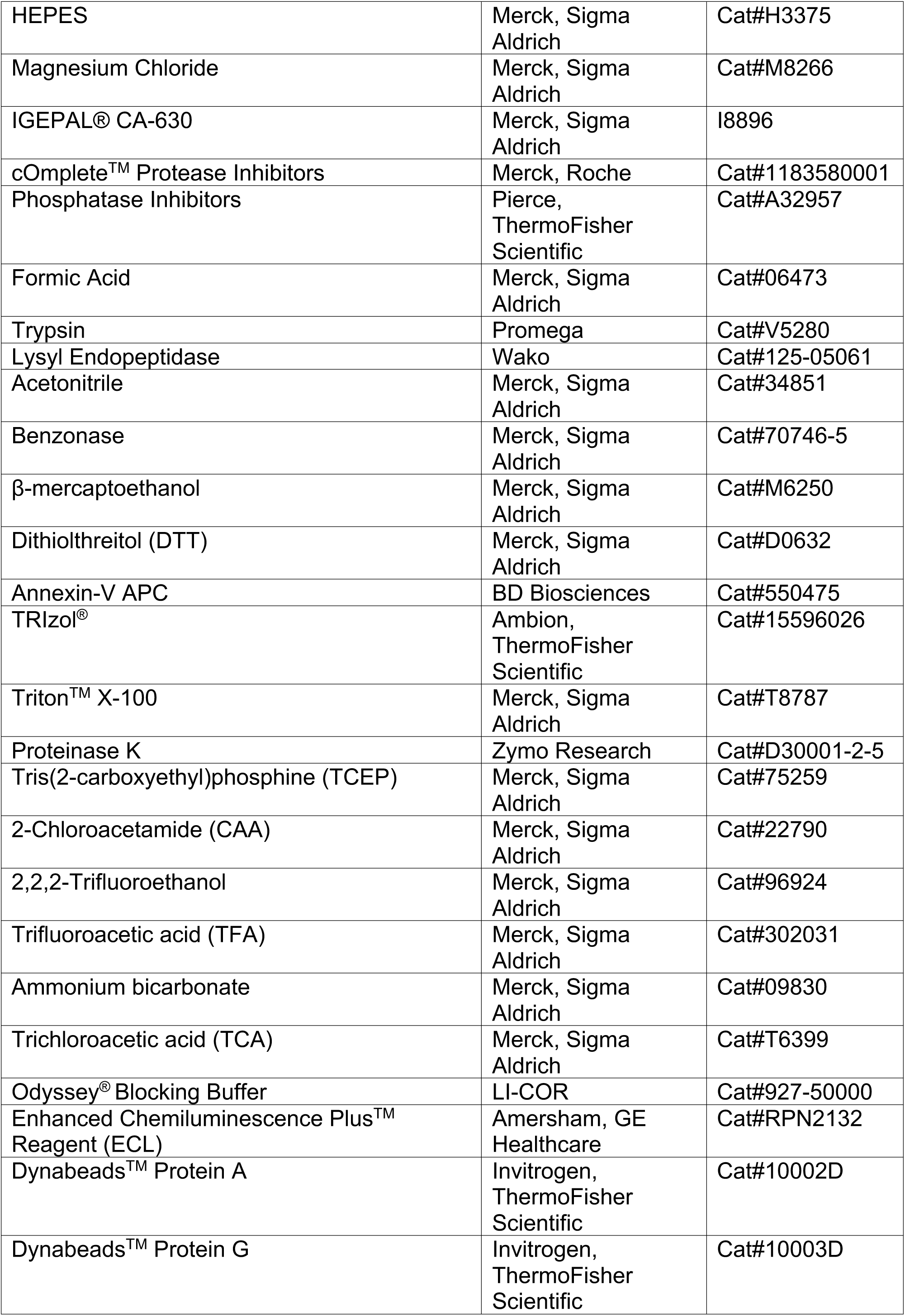
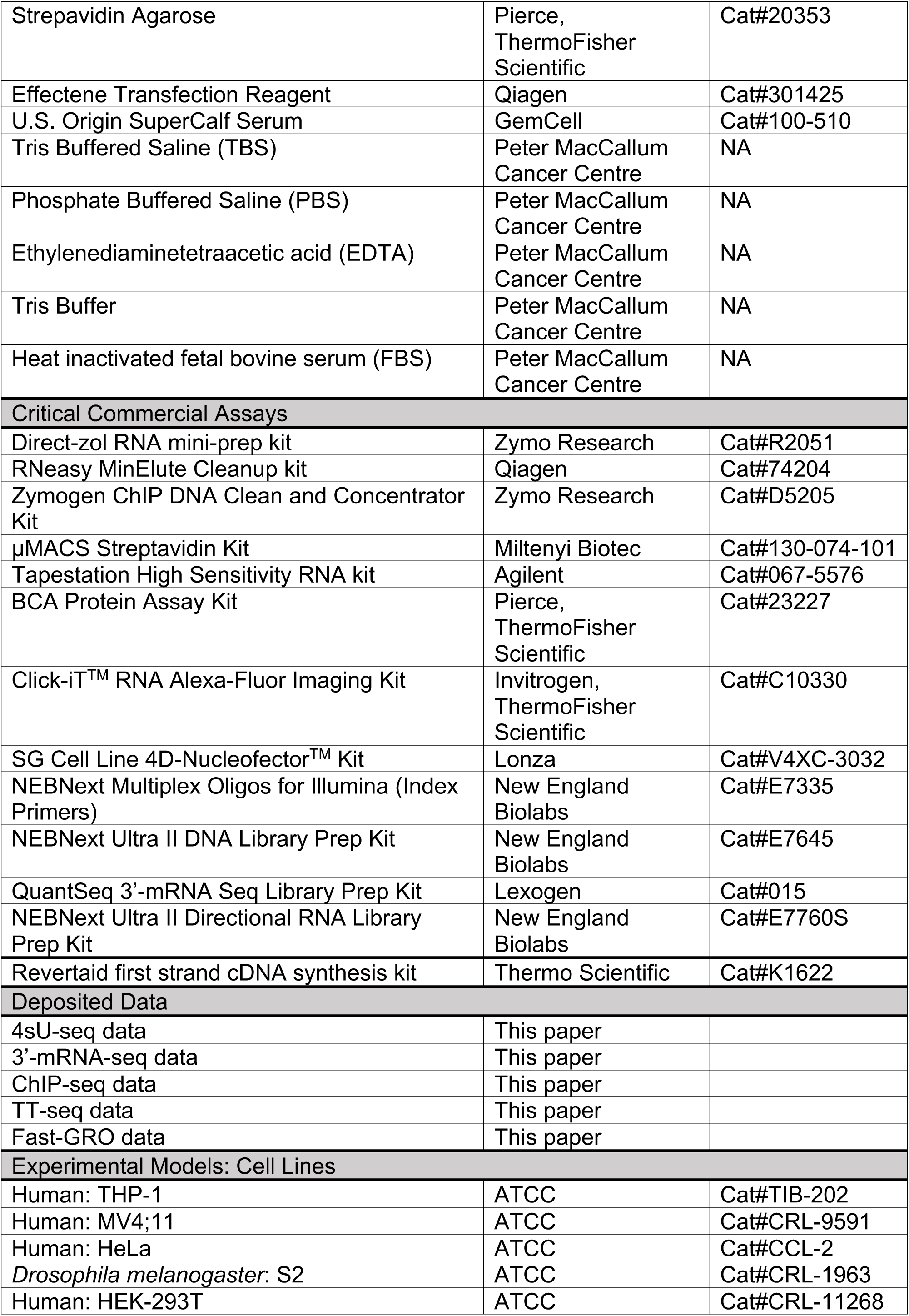
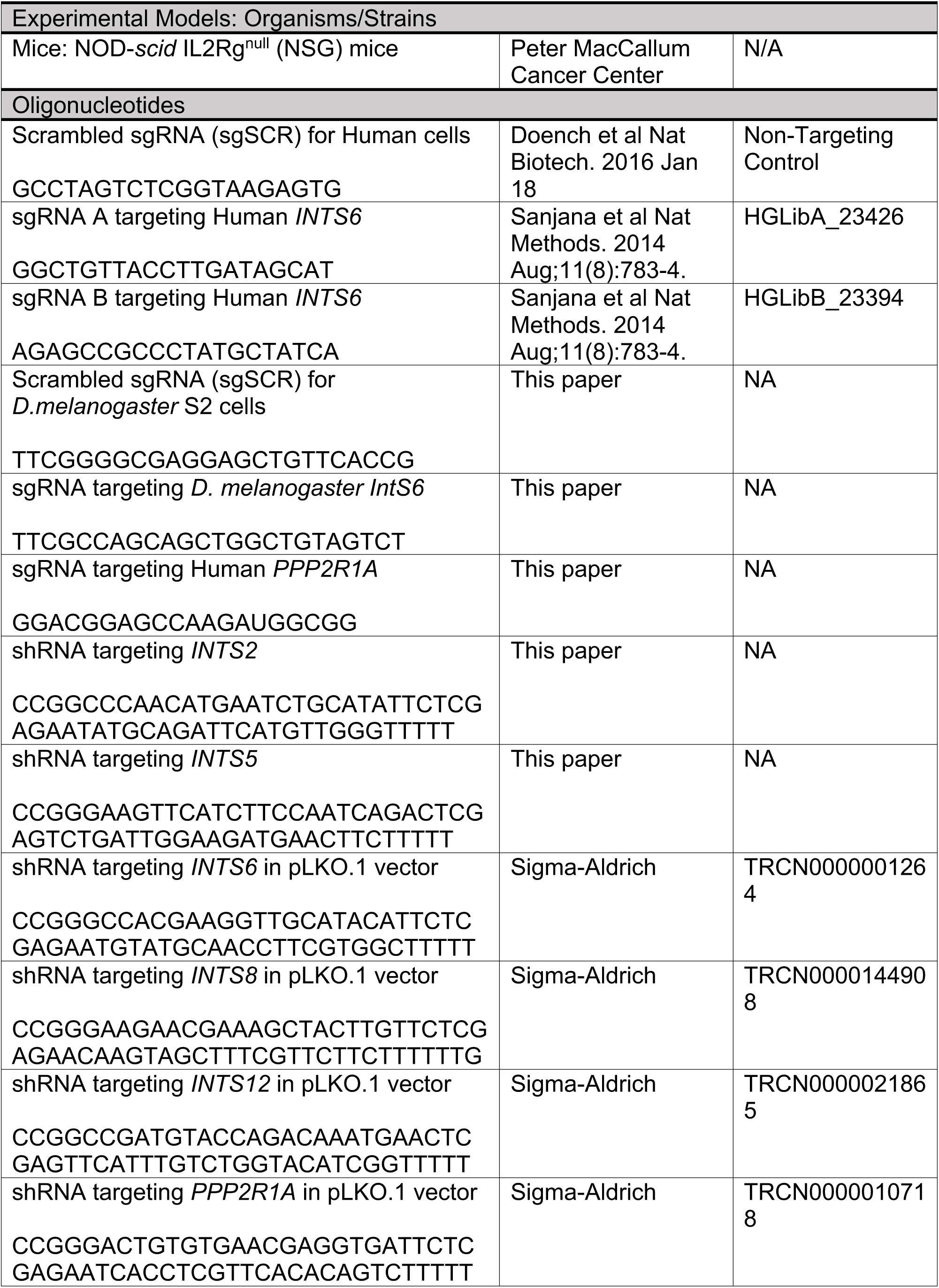
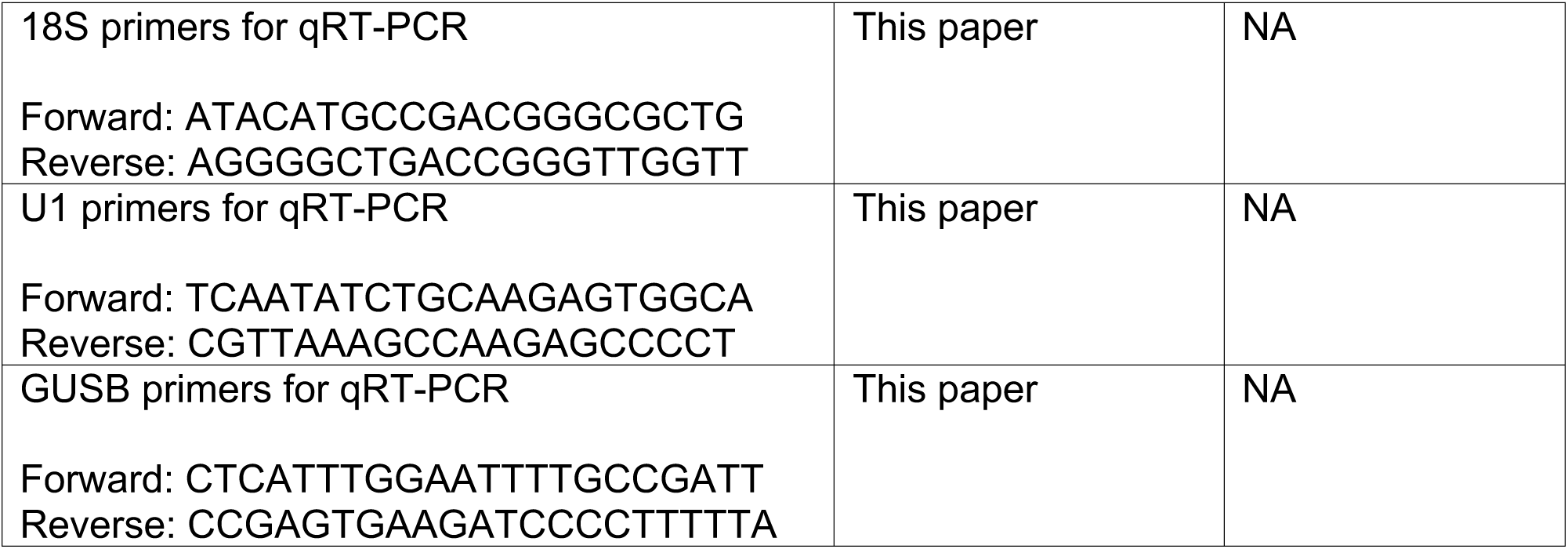
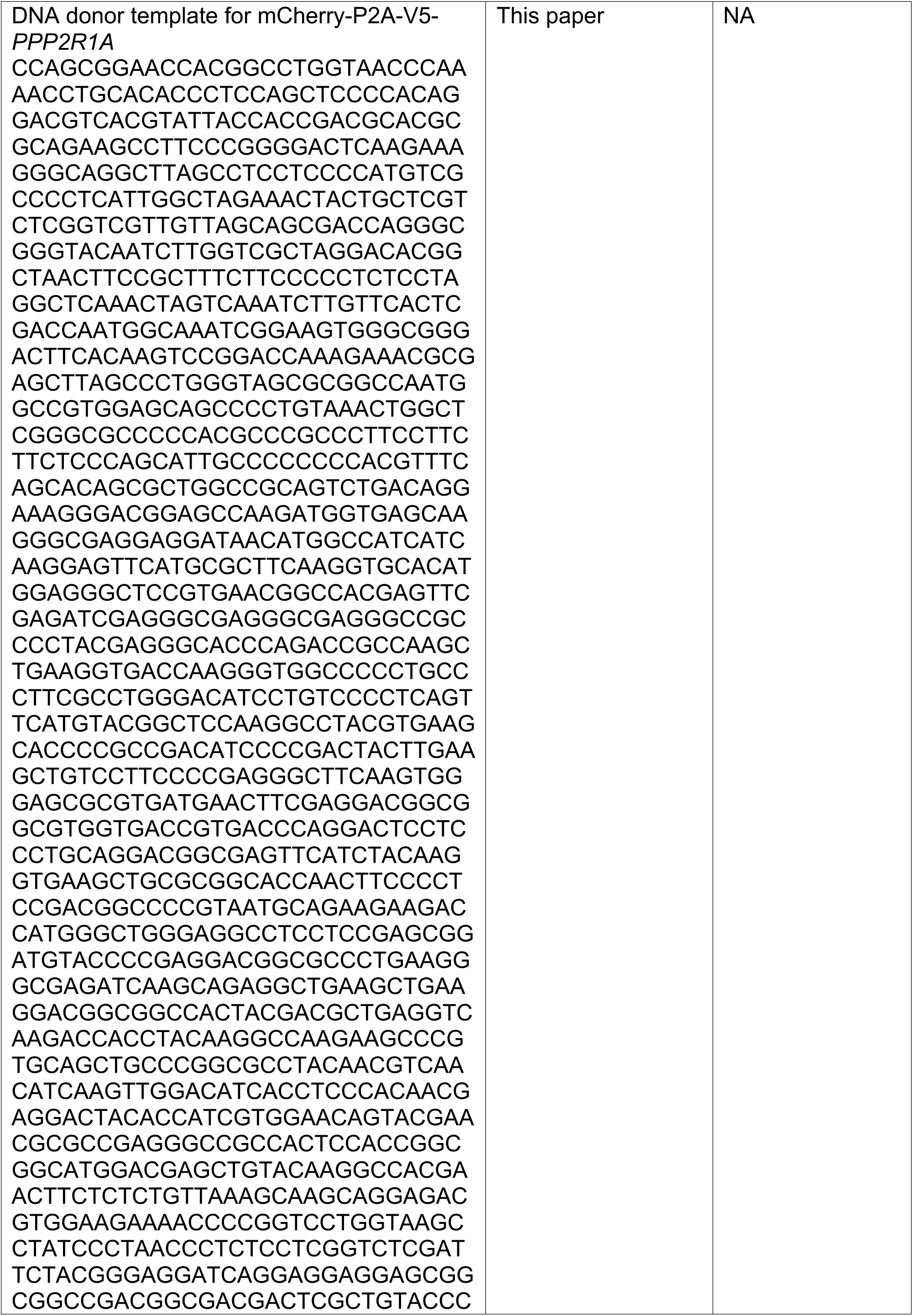
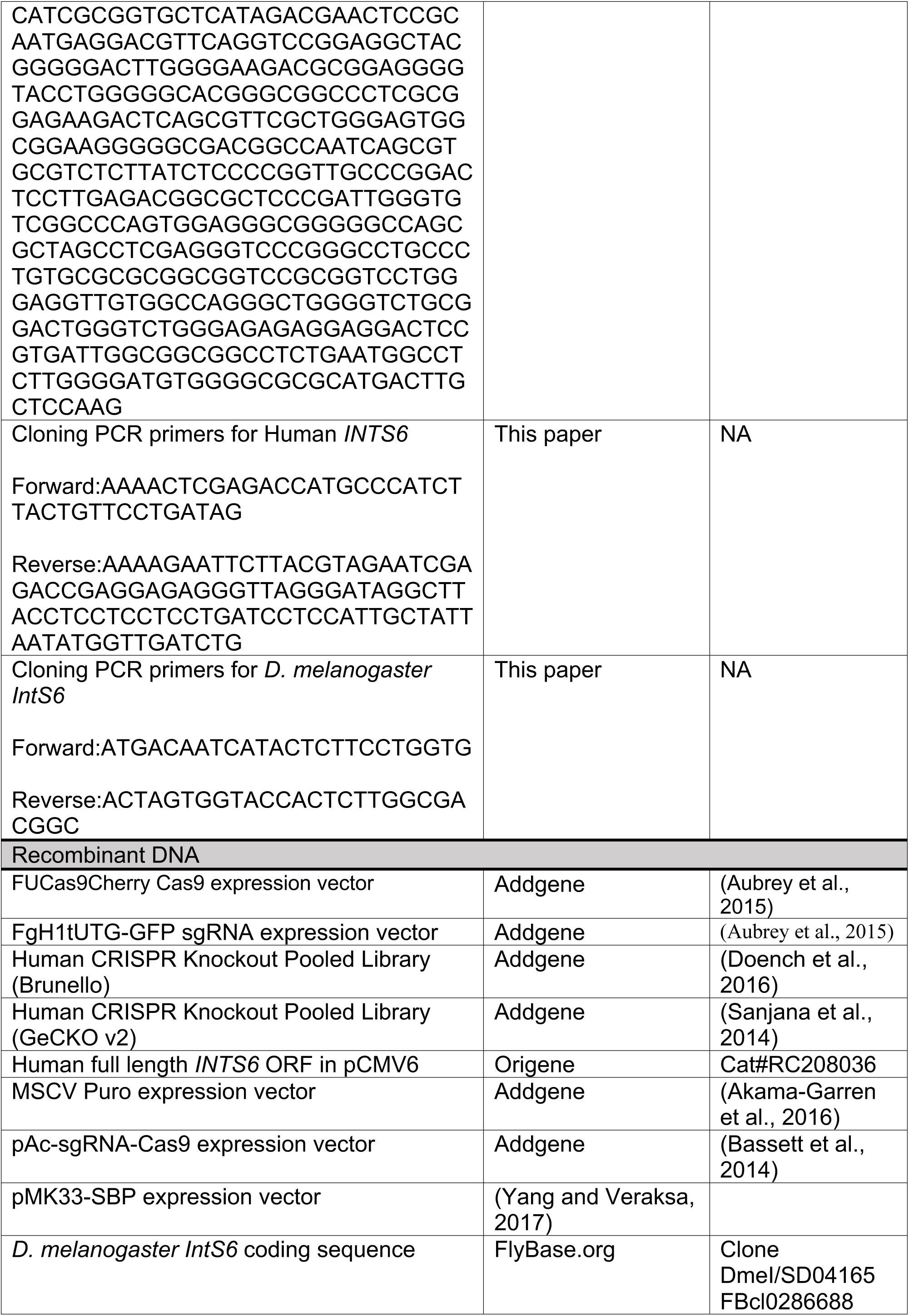
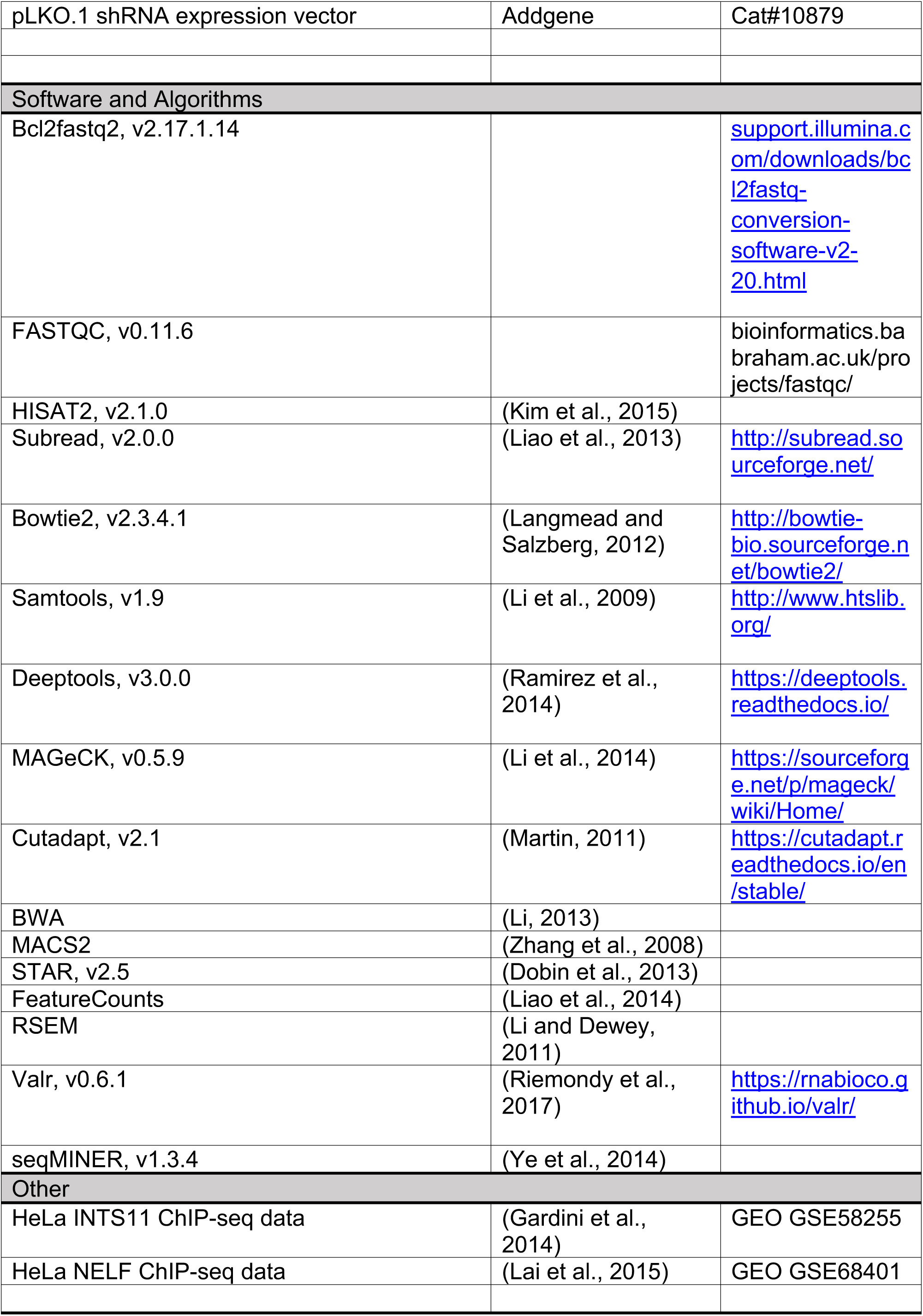

---

**SUPPLEMENTARY FIGURE 1:**
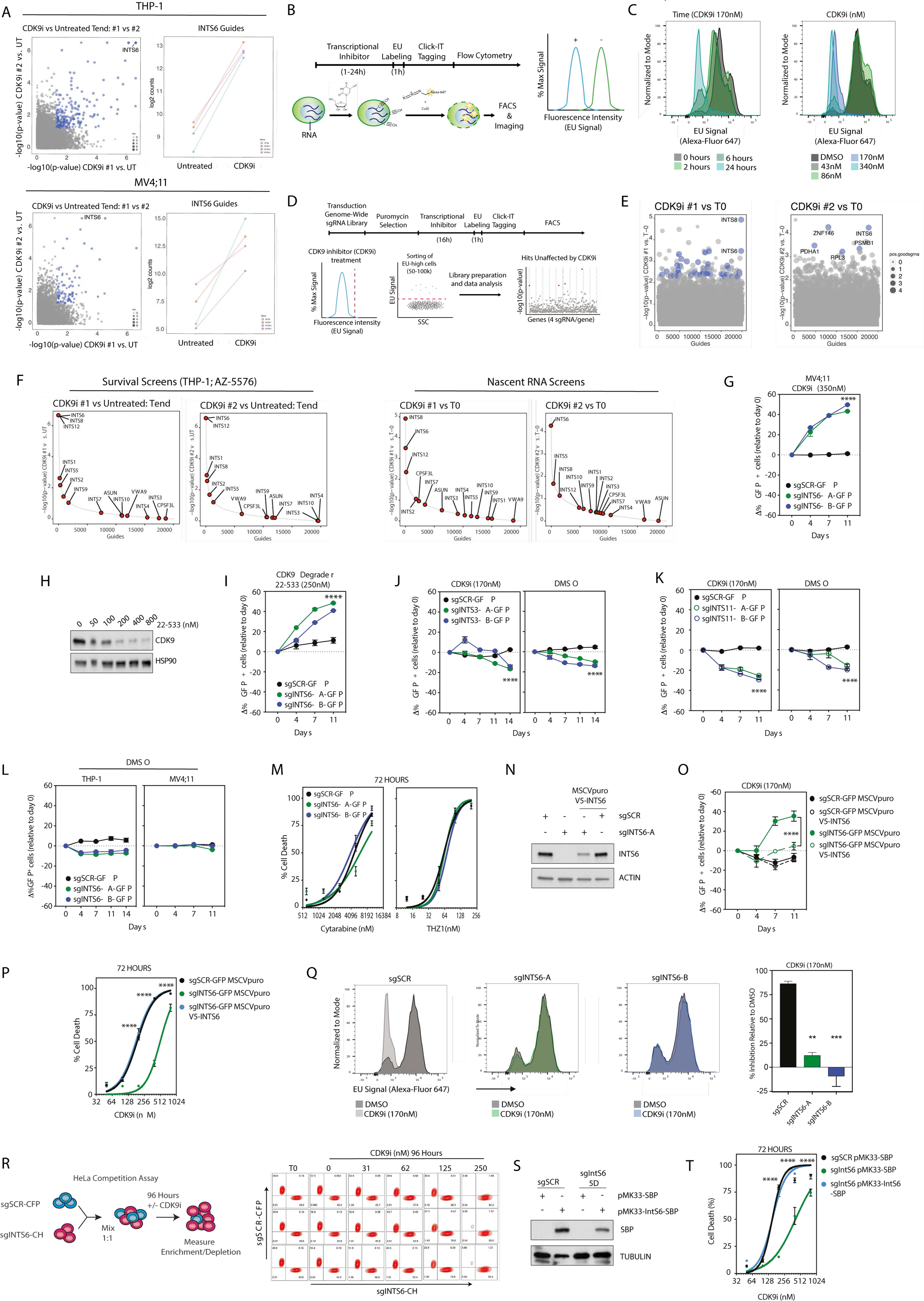
Loss of INTS6 confers resistance to CDK9 inhibition. **A.** Comparison of enriched sgRNAs for replicate survival screens in THP-1-Cas9 and MV4;11-Cas9 cells (CDK9i versus untreated at Tend) plus enrichment of INTS6 sgRNAs in CDK9i-treated versus untreated cells at Tend. **B.** Overview of nascent RNA analysis using ClickIT-EU assay and flow cytometry. **C.** Representative flow cytometry profiles of THP-1 cells treated with CDK9i (170nM) for the indicated time-points (left) or for 24 hours with the indicated dose of CDK9i (right). **D.** Overview of nascent RNA CRISPR-Cas9 genome-wide screen in THP-1-Cas9 cells; resistant cells were selected by FACS for the maintenance of high-EU signal in the presence of CDK9i. **E.** Identification of significantly enriched guides in EU-high cells (CDK9i treated relative to T0). **F.** Enrichment of sgRNAs for Integrator complex subunits in THP-1-Cas9 cells for individual replicate survival screens (significance relative to DMSO-treated cells at Tend) and nascent RNA screens (significance relative to T0). **G.** Competitive proliferation assay for MV4;11-Cas9 cells expressing indicated *INTS6* targeting sgRNAs in the presence of CDK9i. **H.** Western blot of THP-1 cells treated with 22-533 for 6 hours. **I.** Competitive proliferation assay for THP-1-Cas9 cells expressing indicated SCR or *INTS6* targeting sgRNAs in the presence of 22-533. **J.** Competitive proliferation assay for THP-1-Cas9 cells expressing indicated SCR or *INTS3* targeting sgRNAs in the presence of CDK9i or DMSO. **K.** Competitive proliferation assay for THP-1-Cas9 cells expressing indicated SCR or *INTS11* targeting sgRNAs in the presence of CDK9i or DMSO. **L.** Competitive proliferation assay for THP-1-Cas9 or MV4;11-Cas9 cells expressing indicated SCR or *INTS6* targeting sgRNAs in the presence of DMSO. **M.** Annexin-V analysis of THP-1-Cas9 cells expressing indicated *INTS6* targeting sgRNAs treated as indicated for 72 hours. **N.** Western blot of THP-1-Cas9 cells expressing SCR or *INTS6*-A targeting sgRNAs plus/minus V5-INTS6 overexpression. **O.** Competitive proliferation assay and **P.** Annexin-V analysis of THP-1-Cas9 cells expressing indicated SCR or *INTS6*-A targeting sgRNAs plus/minus V5-INTS6 overexpression in the presence of A5576 (CDK9i). **Q.** Representative flow-cytometry profiles of THP-1-Cas9 cells expressing indicated SCR or *INTS6* targeting sgRNAs and treated as indicated for 24 hours, plus relative inhibition of nascent RNA production following CDK9i treatment. **R.** Overview and representative flow cytometry profiles of HeLa cell competitive proliferation assay: cells expressing sgSCR (CFP-Cas9) and sgINTS6 (mCherry-Cas9) were mixed 1:1 and incubated for 96 hours with indicated doses of CDK9i. **S.** Western blot of *D. melanogaster* S2-Cas9 sgSCR and sg*IntS6*-5D cells plus/minus SBP-IntS6 over-expression. **T.** Annexin-V analysis of *D. melanogaster* S2-Cas9 cells expressing indicated SCR or *IntS6* targeting sgRNAs plus/minus SBP-IntS6 overexpression treated as indicated for 72 hours. Blue dots (Figures A and E) represent nominal p-value < 0.05 or p-value < 0.01 respectively. Figures G, H, I J, K, L, M, N, O, P, S, Q, T are representative of 3 independent experiments. Figures G, I, J, K, M, O, P, Q and T were analyzed by 2-way ANOVA, ** p < 0.01, *** p< 0.01, **** p<0.0001.

**SUPPLEMENTARY FIGURE 2:**
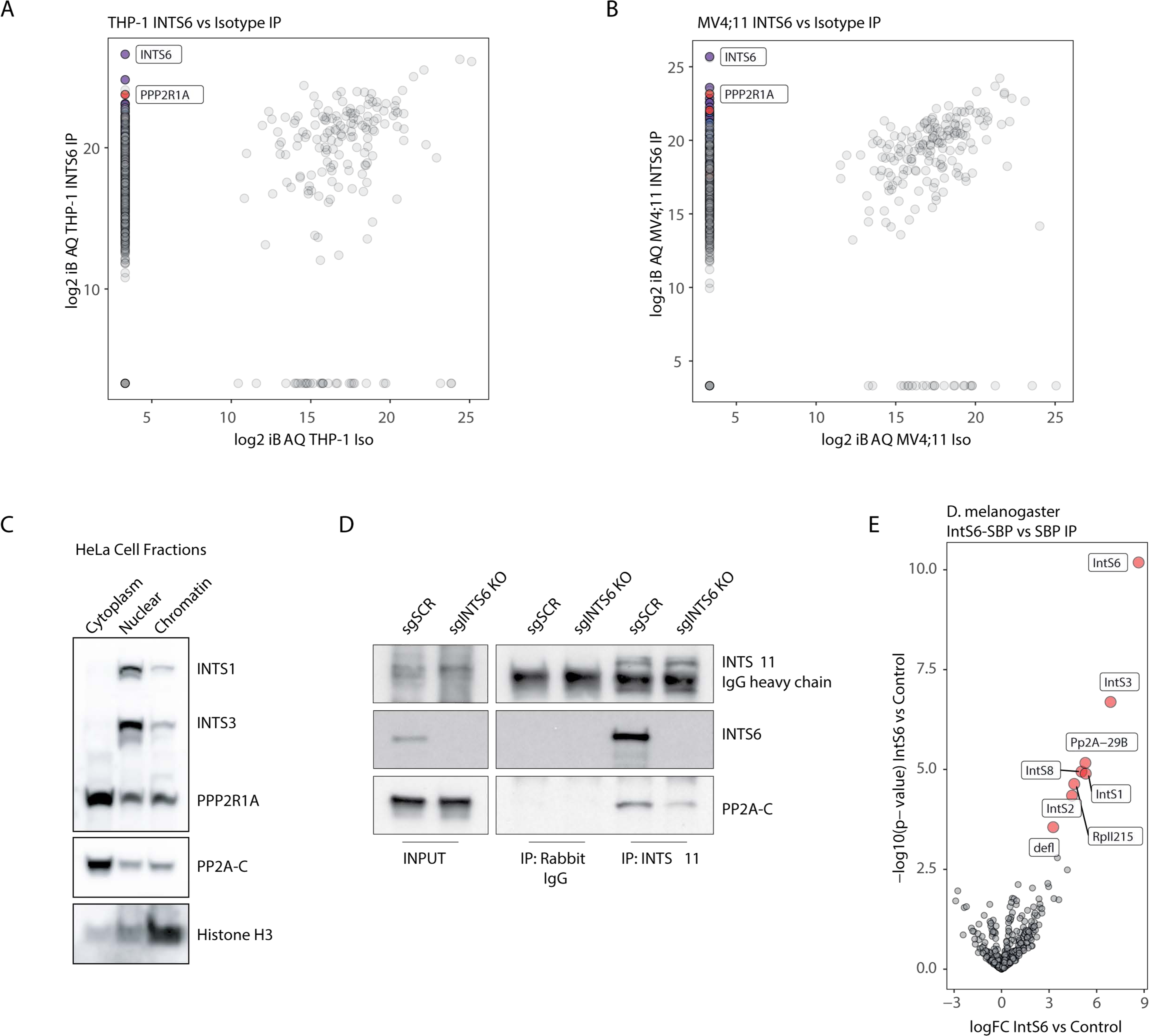
INTS6 bridges the interaction between Integrator and PP2A. Log2 iBAQ scores of proteins identified in INTS6 IP versus mouse IgG isotype control IP mass spectrometry experiments for **A.** THP-1 and **B.** MV4;11 cells. **C.** Western blot of HeLa cell cytoplasmic, nuclear, and chromatin fractions. **D.** Co-IP western blot for INTS11 in THP-1-Cas9 sgSCR and sgINTS6-KO cells. **E.** Volcano plot of differentially enriched proteins in SBP-IntS6 versus SBP (isotype control) streptavidin-IP mass spectrometry experiments in *D. melanogaster* S2 cells.

**SUPPLEMENTARY FIGURE 3:**
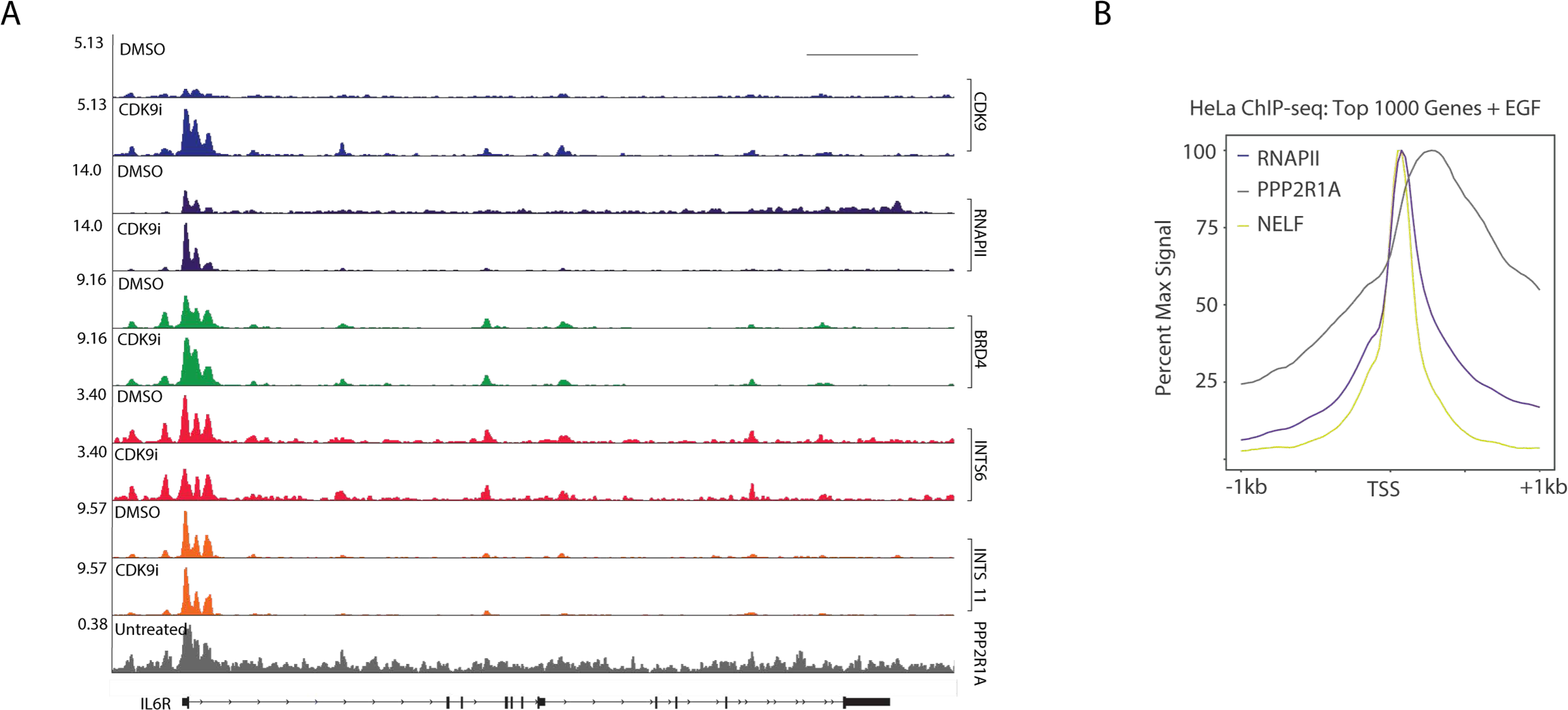
INTS6-dependent dynamic recruitment of PP2A at actively transcribed genes. **A.** Representative IGV ChIP-profiles for indicated proteins in THP-1 cells treated as indicated for 2 hours. **B.** Average profiles of ChIP-seq signal for RNAPII, PPP2R1A, and NELF around TSS profiles at highest-expressed genes (n=990) in HeLa cells treated acutely with EGF (0.1µg/mL; 15 minutes). NELF ChIP-seq tracks are from a previously published dataset (Lai et al., 2015). Scale bar for A represents 10kb.

**SUPPLEMENTARY FIGURE 4:**
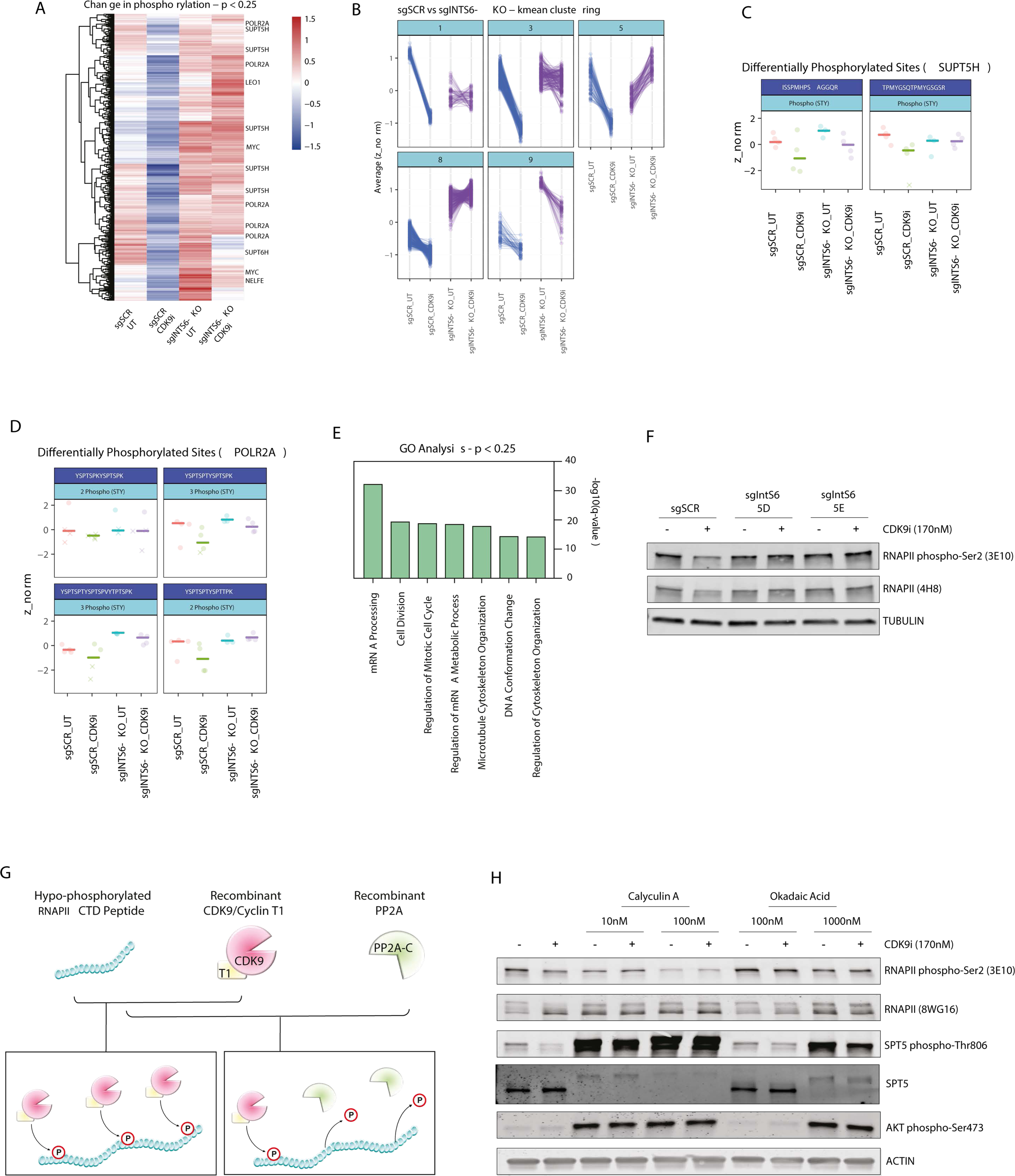
Loss of INTS6/PP2A results in decreased turnover of CDK9 substrates. **A.** Heatmap of Z-scores of phospho-peptides in THP-1-Cas9 sgSCR and sgINTS6-KO cells treated as indicated for 2 hours, p-value < 0.25. **B.** K-means clustering of differentially phosphorylated peptides identified by phospho-peptide mass spectrometry in THP-1-Cas9 sgSCR and sgINTS6-KO cells treated as indicated for 2 hours. Additional differentially phosphorylated peptides **C.** *SUPT5H* and **D.** *POL2RA* peptides in THP-1-Cas9 sgSCR and sgINTS6-KO cells treated as indicated for 2 hours. **E.** GO-analysis of proteins with differentially phosphorylated peptides in CDK9i-treated THP-1-Cas9 sgSCR (compared to DMSO-treated THP-1-Cas9 sgSCR and CDK9i-treated THP-1-Cas9 sgINTS6-KO cells; p < 0.25). **F.** Western blot of *D. melanogaster* S2-Cas9 cells expressing indicated sgRNAs and treated as indicated for 2 hours. **G.** Overview of *in vitro* recombinant CDK9/Cyclin-T1 and PP2A kinase / phosphatase assay. **H.** Western blot of THP-1 cells treated as indicated (15 minutes pre-treatment with Calyculin A or Okadaic acid and 2 hours with CDK9i).

**SUPPLEMENTARY FIGURE 5:**
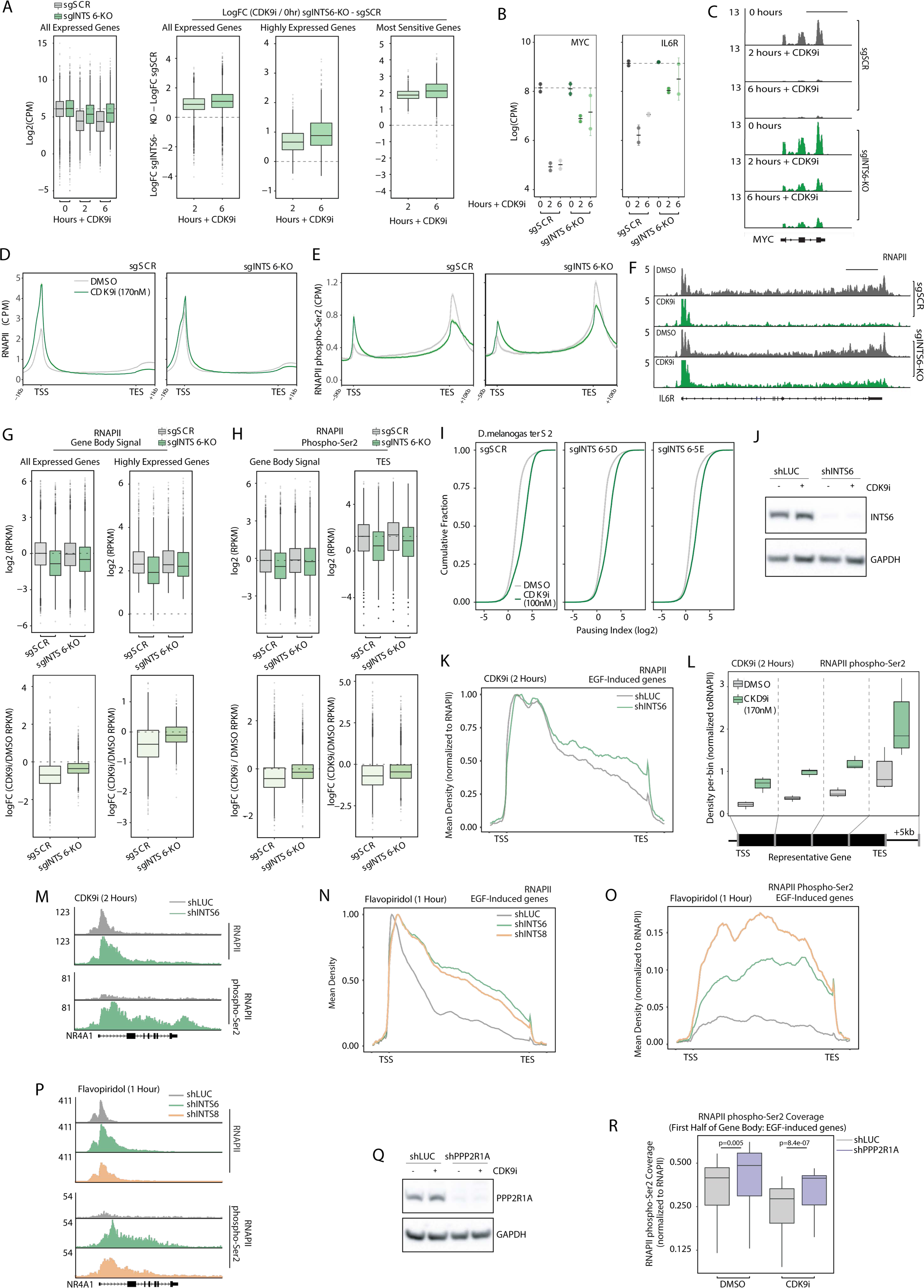
PP2A/INTS6 loss overrides CDK9i induced transcriptional pausing. **A.** 4sU-seq signal (log2CPM) across all expressed genes and difference in log fold change of 4sU-seq signal (Figure 5B-D) between THP-1-Cas9 sgINTS6-KO and sgSCR cells for all expressed genes, highly expressed genes and CDK9i-sensitive genes as indicated. **B.** 4sU-seq signal for *MYC* and *IL6R* genes in THP-1-Cas9 sgSCR and sgINTS6-KO cells treated as indicated. **C.** Example of 4sU-seq signal at the *MYC* locus under indicated conditions. Average profile of **D.** RNAPII and **E.** RNAPII phospho-Ser2 in THP-1-Cas9 sgSCR and sgINTS6-KO cells treated as indicated for 2 hours. **F.** Representative IGV ChIP-profiles for RNAPII under the same conditions. **G.** RNAPII ChIP-seq total and log fold change gene body signal at all genes and highly expressed genes in THP-1-Cas9 sgSCR and sgINTS6-KO cells treated as indicated for 2 hours. **H.** RNAPII phospho-Ser2 ChIP-seq total and log fold change gene body signal (left) and seq total and log fold change TES signal (right) for the same conditions. **I.** Pausing ratio in *D. melanogaster* S2-Cas9 cells expressing indicated sgRNAs and treated as indicated for 2 hours. **J.** Western blot of shLUC and shINTS6 infected HeLa cells treated with CDK9i for 2 hours. **K.** Average profile of RNAPII ChIP-seq signal at EGF response genes (n=50) in shLUC and shINTS6 infected HeLa cells treated with CDK9i for 2 hours and acutely treated with EGF (0.1µg/mL; 15 minutes). **L.** Quantification of Fig. 5I, RNAPII phospho-Ser2 ChIP-seq signal across gene body quartiles. **M.** RNAPII and RNAPII phospho-Ser2 ChIP-seq signal at the *NR4A1* locus under the same conditions. Average profile of **N.** RNAPII and **O.** RNAPII phospho-Ser2 ChIP-seq signal at EGF-response genes (n=50) in shLUC, shINTS6 or shINTS8 infected HeLa cells treated with Flavopiridol (2µM) for 1 hour and acutely treated with EGF (0.1µg/mL; 15 minutes). **P.** RNAPII and RNAPII phospho-Ser2 ChIP-seq signal at the *NR4A1* locus under the same conditions. **Q.** Western blot of shLUC and shPPP2R1A infected HeLa cells treated with CDK9i for 2 hours. **R.** RNAPII phospho-Ser2 ChIP-seq signal across the first half of the gene body for EGF-response genes (n=50), in EGF-treated shLUC or shPPP2R1A HeLa cells, and treated as indicated for 2 hours. Scale bar for C represents 5kb. Scale bar for F represents 10kb.

**SUPPLEMENTARY FIGURE 6.**
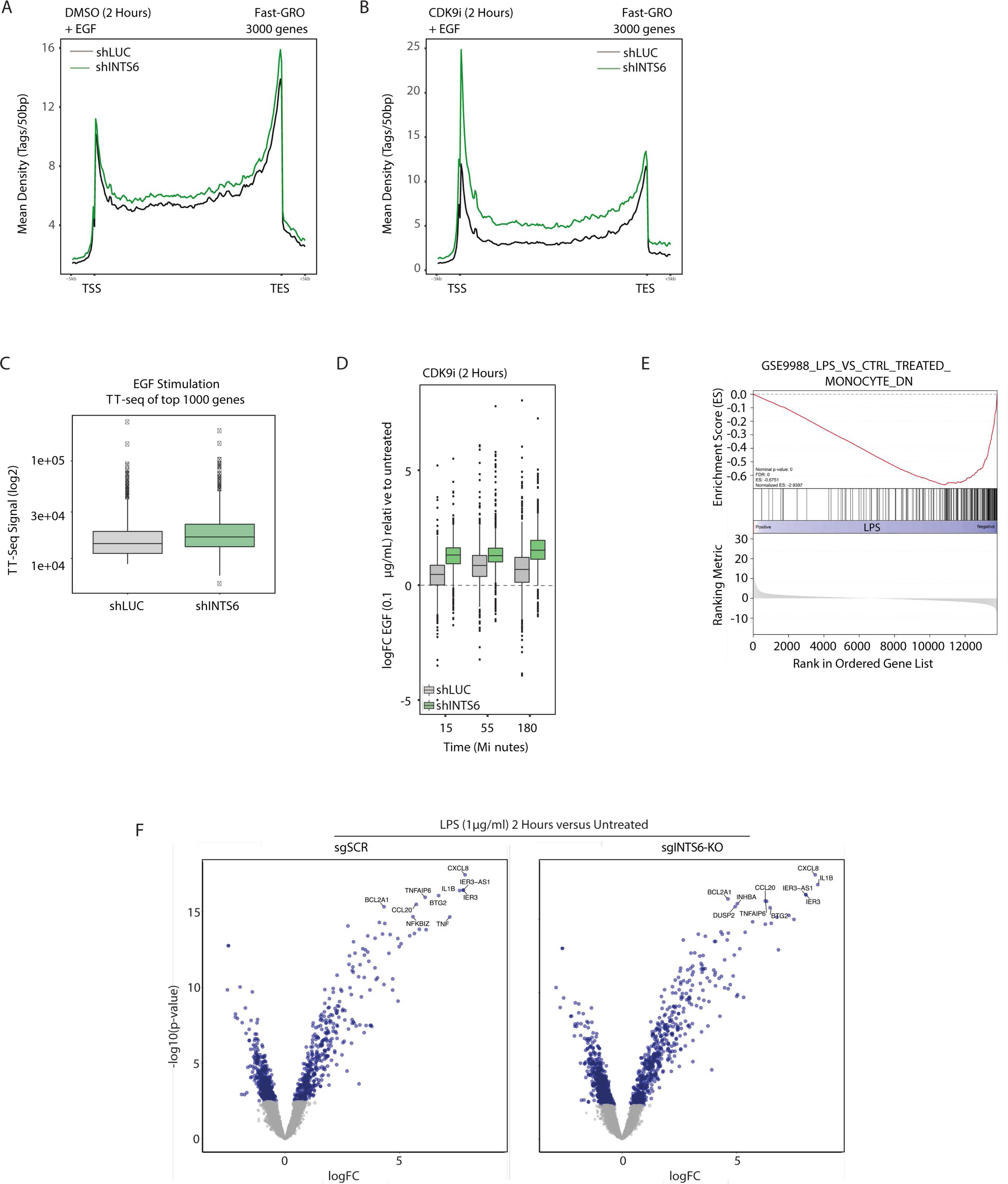
The INTS6/PP2A axis fine-tunes acute transcriptional responses. Average Fast-GRO signal for highest expressed genes (n=2989) in shLUC and shINTS6 infected HeLa cells treated with EGF (0.1µg/mL; 15 minutes) and **A.** DMSO or **B.** CDK9i for 2 hours. **C.** GSEA profile of THP-1-Cas sgSCR cells treated with LPS (2 hours versus 0 hours). **C.** TT-seq signal in HeLa shLUC or shINTS6 cells for highest expressed genes (n=990) after EGF treatment (0.1µg/mL; 15 minutes). **D.** Expression of EGF-induced genes (n=50) in HeLa shLUC or shINTS6 cells treated with CDK9i for 2 hours followed by treatment with 0.1μg/mL EGF for the indicated time-points. **E.** GSEA profile of THP-1-Cas sgSCR cells treated with LPS (2 hours versus 0 hours). **F.** Volcano plot of differentially expressed genes in THP-1-Cas9 sgSCR and sgINTS6-KO cells treated with LPS for 2 hours (relative to untreated control).

**SUPPLEMENTARY FIGURE 7:**
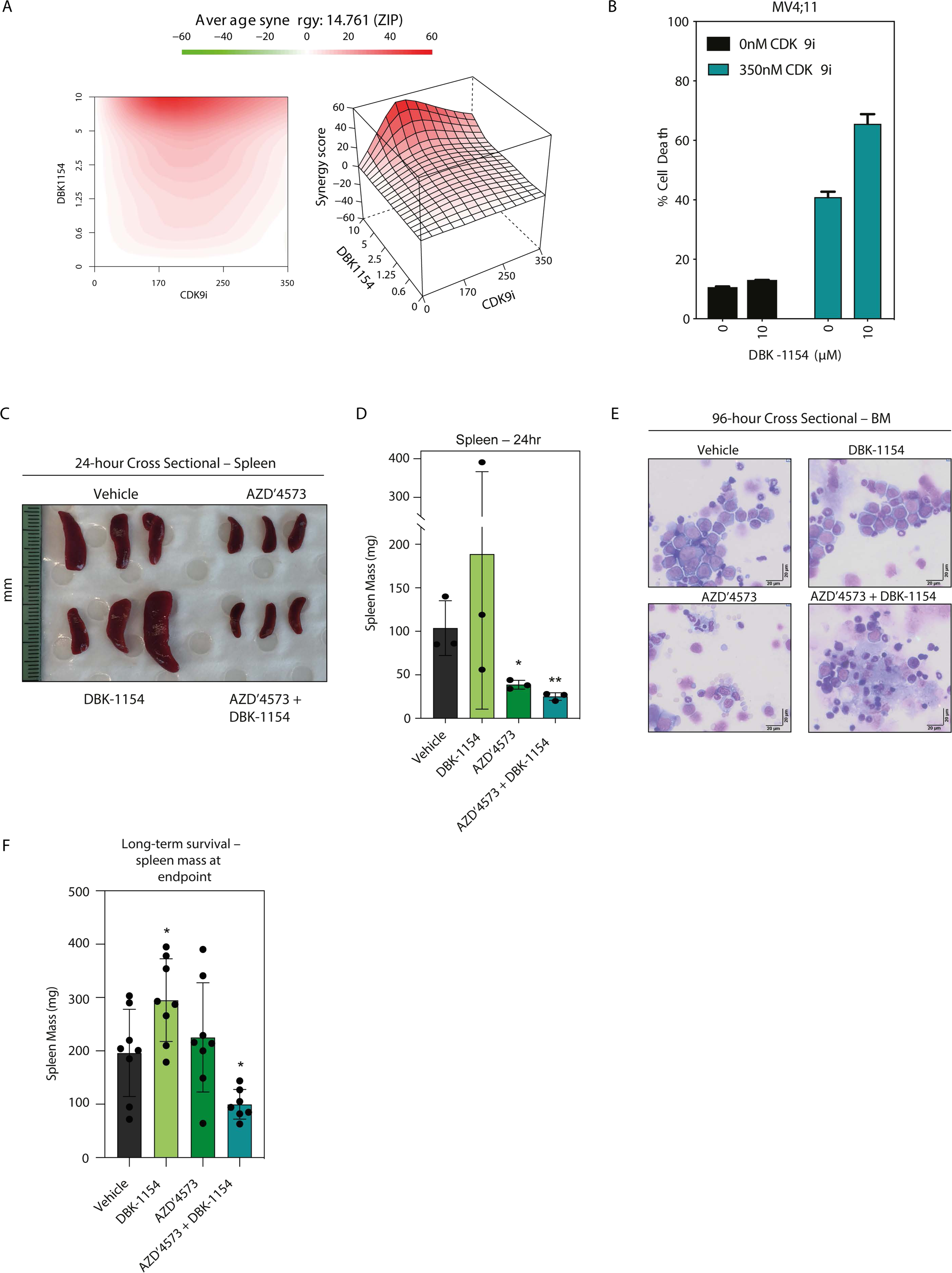
Therapeutic and molecular synergy between PP2A agonist and CDK9i. **A.** Synergy for combined CDK9i and DBK-1154 treatment of THP-1-Cas9 sgSCR cells was determined using the SynergyFinder computational package and the ZIP synergy index and is denoted as regions of red in the graphs. **B.** Annexin-V analysis of MV4;11 cells treated as indicated for 72 hours. **C.** Spleens and D. spleen masses from mice treated as indicated at 24 hours post-therapy. Scale bar in millimetres (mm). **E.** May-Grünwald Giemsa histological staining of bone marrow cytospins from mice treated as indicated 96 hours post therapy. Scale bar represents 20 μm. **F.** Spleen masses of long-term survival experiment mice at endpoint.

